# Evolution of mutational fitness effects in island populations

**DOI:** 10.64898/2026.07.14.738531

**Authors:** Emma K. Howell, Lauren E. Nolfo-Clements, Felix Baier, Bret A. Payseur

## Abstract

The distribution of fitness effects (DFE) quantifies the selective consequences of newly arising mutations. Theoretical and empirical investigations of the DFE suggest that it is highly context-specific, shaped by both intrinsic properties of an organism (e.g., biological complexity) and extrinsic properties of a population (e.g., environment). Despite recent comparisons of the DFE between populations and species, little is known about how this distribution changes over shorter evolutionary timescales. Islands provide a powerful framework for understanding the impact of recent shifts in selection on the DFE, as founding populations often experience abrupt environmental changes. Using whole-genome, population-level sequence data, we investigate how such extreme transitions shape the DFE in two island radiations of *Peromyscus* mice: white-footed mice (*P. leucopus*) in Massachusetts’s Boston Harbor and deer mice (*P. maniculatus*) in the Gulf Islands of British Columbia. To measure the extent to which the selective effects of mutations have diverged between island and mainland populations, we leverage recent advances that extend DFE inference to multiple populations. By reconstructing the “joint DFE”, we estimate both the strength of selection acting on distinct mutational classes and the correlation in mutational fitness effects between populations. We find that mutational fitness effects have diverged between island and mainland populations, despite the relative recency of these radiations. Comparisons between island populations, which reveal higher fitness effect correlations, suggest that this feature of the joint DFE captures broad-scale divergence in the environment populations inhabit. Together, our discoveries provide a rare empirical example of divergent environments shaping genome-wide patterns of fitness-affecting genetic variation in natural populations.

## INTRODUCTION

The movement of organisms onto islands demarcates a major evolutionary transition. Populations on islands face unusual environments that differ from the mainland across multiple ecological axes, including predation, competition, climate, habitat diversity, and resource availability (Crowell, 1983; Lomolino, 1985; Adler & Levins, 1994; Lomolino, 2005). In addition, the unique geographic setting of islands constrains dispersal and migration, which in turn influences broad-scale patterns of species richness (MacArthur & Wilson, 1963; MacArthur & Wilson, 1967a). Evolution on islands also carries severe genetic consequences for inhabitant populations. Initial founding bottlenecks can sharply reduce genetic variation (MacArthur & Wilson, 1967b; Clegg, 2009). Subsequent prolonged periods of small population size can amplify the effects of genetic drift and increase population-level inbreeding (Frankham, 1997; Frankham, 1998). Limits on dispersal and migration can inhibit gene flow and increase the genetic differentiation of island populations (MacArthur & Wilson, 1967b; Clegg, 2009). When combined with the environmental and demographic stochasticity characteristic of islands, this precarious genetic state likely contributes to higher extinction rates of island populations (Frankham, 1997; Frankham, 1998).

Though these vulnerabilities portray islands as evolutionary “dead-ends” for the populations that occupy them, island biota abound with examples of extreme morphological, behavioral, and life-history evolution. Comparisons between island and mainland body sizes across diverse taxa have demonstrated broad adherence to the “island rule”, a pattern in which large-bodied species become smaller on islands and small-bodied species become larger (Foster, 1964; Van Valen, 1973; Lomolino, 1985; Benítez-López et al., 2021). In addition to larger body sizes, island rodents demonstrate reduced reproductive output, lower dispersal, higher population densities, and decreased aggression, a suite of characteristics that have been termed the “island syndrome” (Adler & Levins, 1994). These striking ecogeographical trends suggest that the myriad ecological differences between island and mainland present divergent phenotypic optima for inhabitant populations. Such broad-scale shifts in the selective regime situate islands as powerful settings for understanding how genetic variation affects fitness.

The impacts of mutations on organismal fitness are captured by the distribution of fitness effects (DFE), which describes the selective consequences of mutations that occur within a population (reviewed in Eyre-Walker & Keightley, 2007). The DFE in natural populations can be reconstructed by following the logic that the burden and frequencies of mutations are shaped by their relative contributions to fitness (Eyre-Walker & Keightley, 2007; Chen et al., 2021; Johri et al., 2022). Since observed allele frequencies reflect the combined action of natural selection and genetic drift, most inference frameworks rely on contrasts in allele frequencies between putatively neutral variants and classes of mutations presumed to experience selection (Williamson et al., 2005; Eyre-Walker et al., 2006; Keightley & Eyre-Walker, 2007; Boyko et al., 2008; Galtier, 2016; Tataru et al., 2017; Kim et al., 2017; Huang et al., 2021). The fitness effects of some mutations will depend on the environmental and genetic background on which they arise (Eyre-Walker & Keightley, 2007). This context-dependent property of the DFE, combined with the capacity of existing inference frameworks to account for demographic change, position the DFE as an ideal metric for assaying the broad-scale selective shifts that accompany transitions from mainland to island environments.

To understand how the concurrent environmental and demographic transitions underlying island evolution shape patterns of fitness-affecting variation, we leverage two expansions of endemic *Peromyscus* mice to near-shore islands along the west and east coasts of North America: *P. leucopus* in Massachusetts’ Boston Harbor, and *P. maniculatus* in the Gulf Islands of British Columbia. Continental island systems such as these, which were formed by Pleistocene-era glacial cycles, have more recent origins and clearer ties to their mainland sources than do remote oceanic islands (Martínková et al., 2013). Despite the recency of island formation and the proximity between island and mainland sites, populations of *Peromyscus* in both systems exhibit key phenotypic hallmarks of island evolution. In the Boston Harbor, wild-caught island *P. leucopus* are 40-50% larger in size than mainland mice (Nolfo-Clements et al., 2017). Field studies in the Gulf Islands suggest that *P. maniculatus* adhere to multiple island syndrome characteristics, including larger body sizes, higher population densities, lower dispersal, and reduced intraspecific aggression (Redfield, 1976; Sullivan, 1977; Halpin & Sullivan, 1978; Berg & Nietlisbach, 2025). Common garden experiments support a heritable basis of body size differences in this system, with laboratory-raised *P. maniculatus* weighing 35% more than mainland mice (Baier & Hoekstra, 2019).

Island and mainland populations in each system differ in both the temporal and spatial scale of isolation. These contrasts make it possible to examine the generality of evolutionary dynamics across species, continental island systems, and “replicate” island populations. Leveraging this hierarchical comparative framework, we show that fitness-affecting variation exhibits both shared and system-specific signatures of natural selection and demographic history. Disentangling the contributions of these two determinants by reconstructing multi-population DFEs yields strong evidence for the divergence of mutational fitness consequences between island and mainland populations.

## RESULTS

To understand how mutations affect fitness in the context of island evolution, we harnessed two continental island radiations of *Peromyscus*: *P. leucopus* (white-footed mice) in Massachusetts’ Boston Harbor and *P. maniculatus* (deer mice) in the Gulf Islands of British Columbia (Figure 1). Situated just east of Boston’s city center, the Boston Harbor archipelago is comprised of 34 small islands and peninsulas. Our sample of *P. leucopus* includes wild-caught mice from two of the inner harbor islands: Bumpkin (0.122 km^2^) and Peddocks (0.746 km^2^), and one nearby mainland site, World’s End. The Gulf Islands, situated between Vancouver Island and mainland British Columbia in the southern Strait of Georgia, encompass much larger islands located further from the mainland. Here, our sample of wild-caught *P. maniculatus* spans two of the southern islands, Saturna (31 km^2^) and Pender (34 km^2^), and a representative mainland site, Maple Ridge.

**Figure 1.**
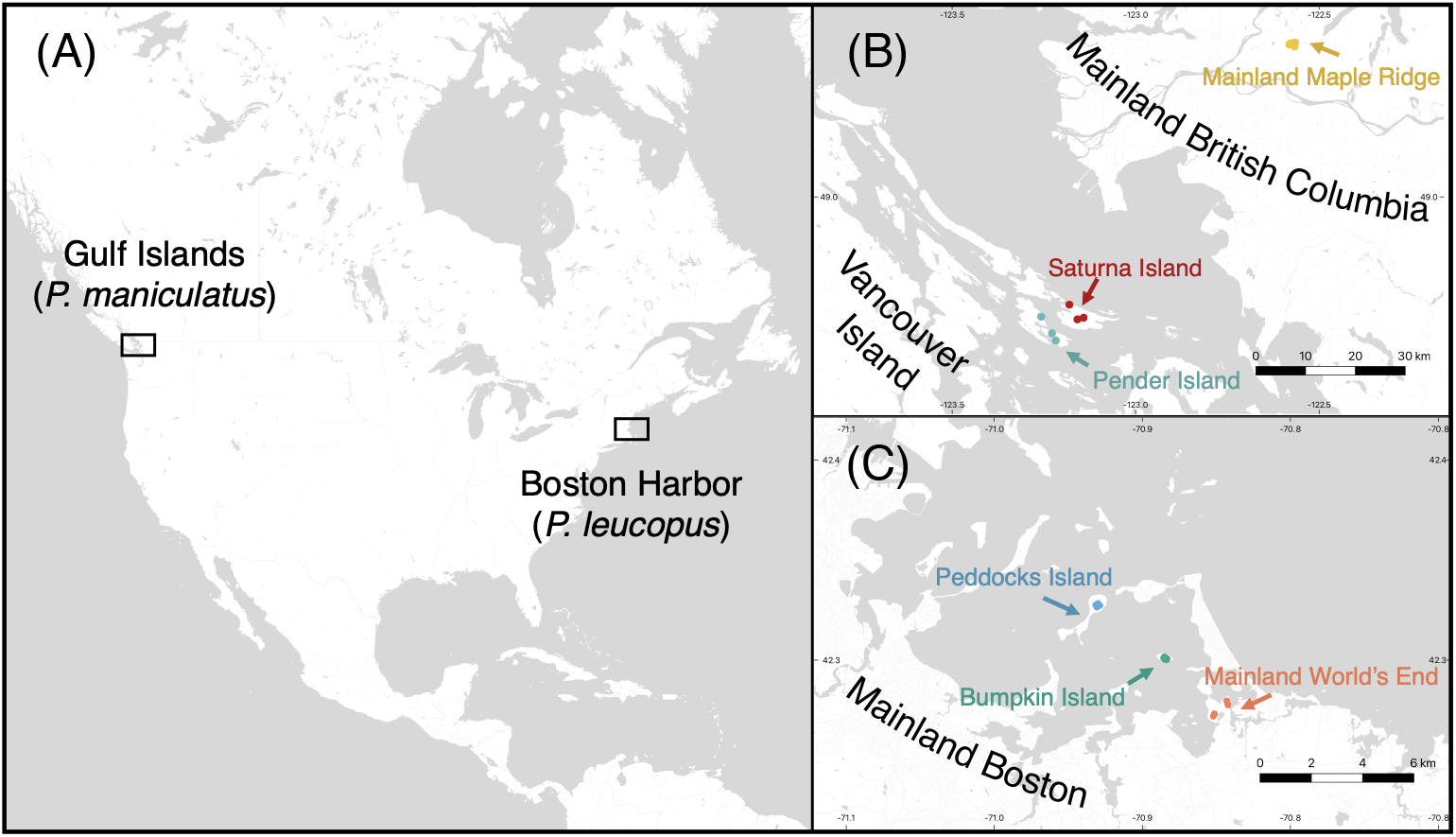
Continental island systems of *Peromyscus* mice. Boxes in panel **A** depict the locations of the two continental island systems characterized in this study: *P. maniculatus* in the Gulf Islands of British Columbia and *P. leucopus* in Massachusetts’ Boston Harbor. Panel **B** depicts the Gulf Islands in greater detail, with sampling sites on Saturna Island (red), Pender Island (light blue), and mainland Maple Ridge (yellow) labeled. Panel **C** depicts the Boston Harbor in greater detail, with sampling sites on Peddocks Island (dark blue), Bumpkin Island (green), and mainland World’s End (orange) labeled. Scale bars measure distance in kilometers. Land areas are depicted in white. Water areas are depicted in gray.

Previous demographic modeling in these systems suggest that splits between island and mainland populations in the Boston Harbor were mediated by rises in sea level following the Last Glacial Maximum (LGM) and subsequent isolation of land areas (Howell et al., 2025a). Although the timescale of divergence between island populations in the Gulf Islands is also consistent with post-LGM vicariance, island and mainland populations exhibit significantly deeper divergence that pre-dates the establishment of contemporary land areas (Howell et al., 2025b). Genomic analyses of each species presented in this study are based on high-coverage (∼30X), short-read, whole genome sequences generated for a subset of unrelated mice sampled from each location (n=13 for Bumpkin Island, n=13 for Peddocks Island, n=17 for mainland World’s End, n=20 for Saturna Island, n=10 for Pender Island, and n=17 for mainland Maple Ridge).

To understand how different classes of mutations affect fitness in island and mainland populations, we contrast patterns of variation within protein-coding genes with those observed in intergenic regions, which are assumed to evolve in a largely neutral fashion (Figure 2A). We partition each gene into 5’ UTR elements, introns, exons, and 3’ UTR elements to capture differences in the functional consequences of mutations occurring in each element type. Within exons, we separately consider the dynamics of mutations that result in synonymous versus nonsynonymous changes to protein sequence.

**Figure 2.**
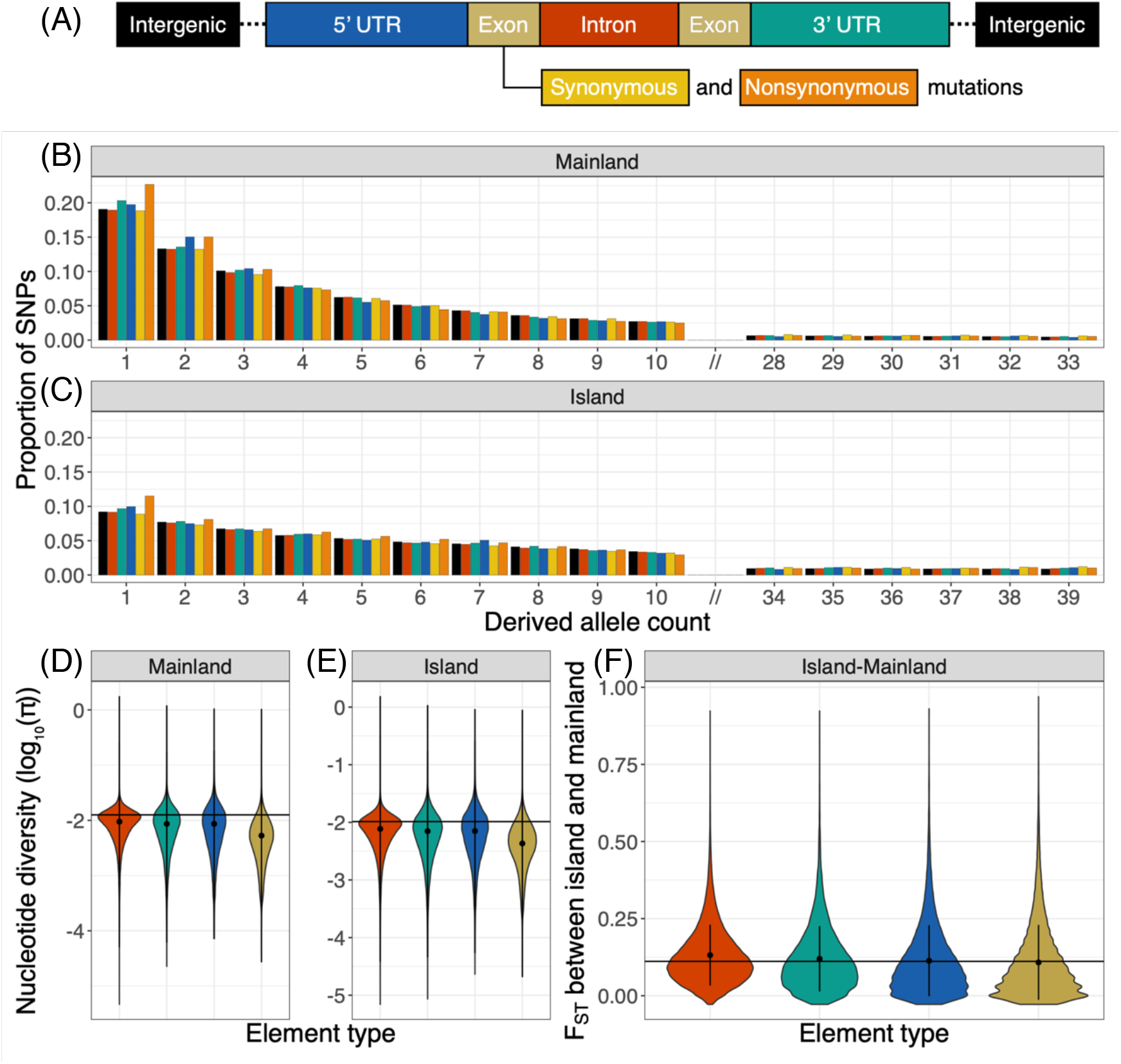
Distinct signatures of purifying selection across genic elements. The schematic in **A** depicts the source of mutation types contrasted in this study. Exons, depicted in khaki for the element-wise summaries of variation in **D**-**F** are further decomposed into synonymous (yellow) and nonsynonymous (orange) mutation types for SFS comparisons in **B**, **C**, and downstream inferences. Panels **B** and **C** depict differences in the unfolded site frequency spectra across mutation types and between mainland **B** and island **C** populations. X-axes show the derived allele count, and y-axes display the proportion of SNPs within a given mutation type that fall in each allele count bin. Panels **D** and **E** depict the distributions of nucleotide diversity (measured as log(π)) across genic element types in the mainland **(D)** and island **(E)** population. Black circles denote the mean of each distribution and vertical black lines denote +/-1 standard deviation around the mean. Horizontal black lines indicate the mean log(π) measured across disjoint intergenic windows. Panel **F** depicts the distributions of F_ST_ measured between island and mainland populations across each element type. As with **D** and **E**, black circles denote the mean of each distribution and vertical black lines denote +/-1 standard deviation around the mean. The horizontal black line indicates the mean island-mainland F_ST_ measured across disjoint intergenic windows. Patterns of variation depicted in **B**-**F** correspond to a representative island-mainland pair of *P. maniculatus* in the Gulf Islands (Saturna Island and mainland Maple Ridge). Observed patterns of variation in other island and mainland populations are depicted in Supplementary Figure S1.

### Contrasting signatures of purifying selection across genic regions

Patterns of variation across genic elements reflect the joint action of natural selection and demographic processes. Signatures of distinct demographic histories are evident across element types in the overall enrichment of intermediate frequency variants in the site frequency spectrum (SFS) of island populations and lower levels of nucleotide diversity compared to their mainland counterparts (Figure 2B-E; Supplementary Figure S1). Differences in the strength of selection acting on mutations within each class of genic elements manifest as element-specific distortions in these summaries of variation. Across island and mainland populations, exons and nonsynonymous mutations display the most extreme departures from patterns of variation observed in intergenic regions, exhibiting the largest reductions in element-wise nucleotide diversity and the greatest skew towards rare alleles in the SFS (Figure 2B-E; Supplementary Figure S1). Differentiation between island and mainland populations, as measured by F_ST_, is lowest within exons and highest within introns (Figure 2F; Supplementary Figure S1). Collectively, these observations are consistent with stronger purifying selection acting on variation in exons compared to other genic elements.

### Purifying selection structures shared variation between island and mainland populations

Given the timescale of island-mainland divergence in these two systems, many of the mutations segregating among populations likely reflect retained ancestral variation rather than *de novo* mutations that arose post-split. Among intergenic SNPs, most segregating variation is either shared between island and mainland or is private to the mainland (Figure 3). This pattern is consistent with larger effective population sizes (*N_e_*) retaining a greater reservoir of ancestral variation on the mainland. Island populations, by contrast, harbor little private variation, which reflects the minority of ancestral variants that were lost or fixed in the mainland plus a smaller contribution of mutations that occurred on the island (Figure 3).

**Figure 3.**
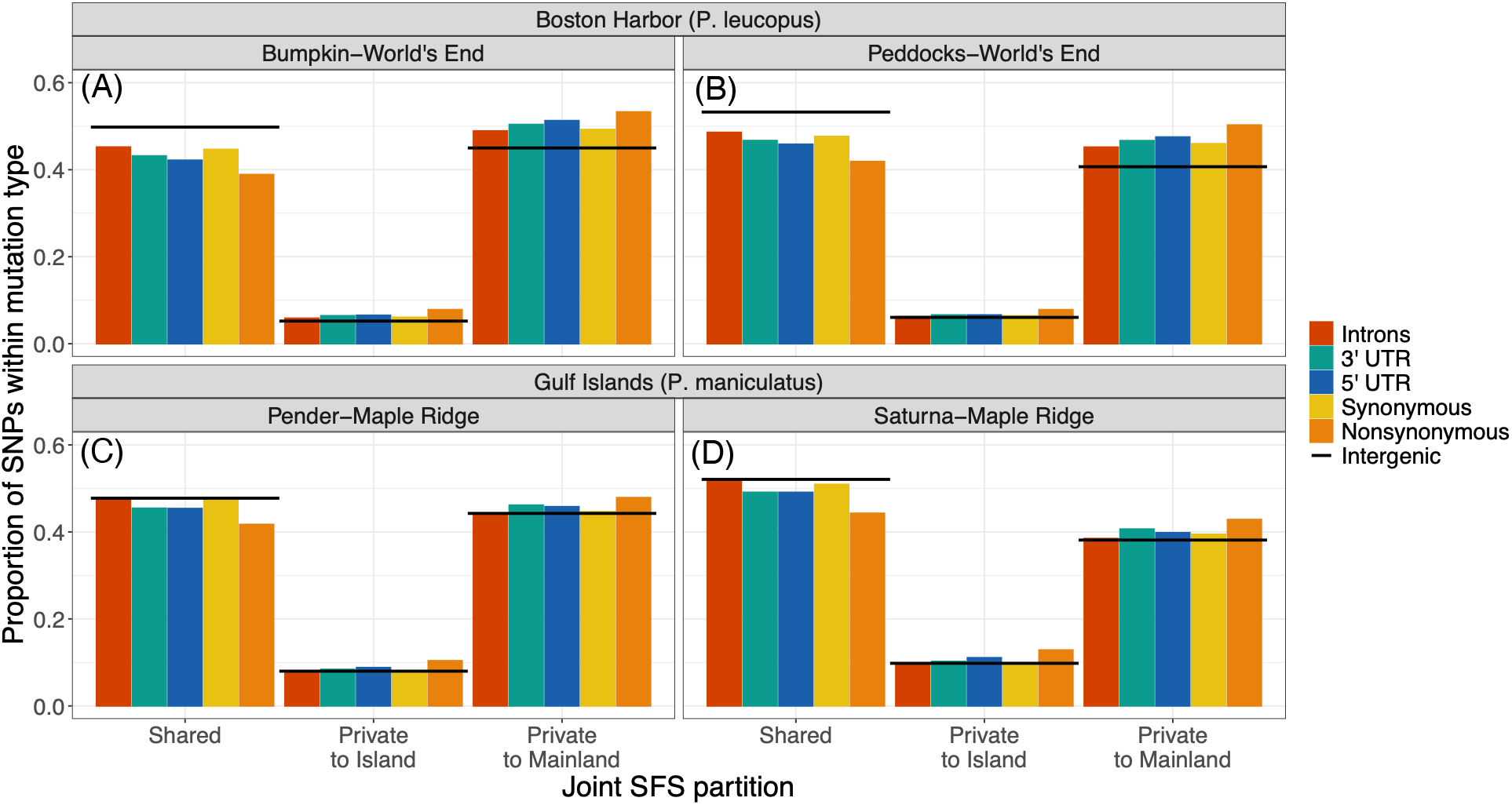
Purifying selection affects how ancestral variation is partitioned between island and mainland populations. Panels **A**-**D** distinguish each of the four island-mainland comparisons that comprise the studied systems. **A** and **B** reflect comparisons between island (Bumpkin and Peddocks) and mainland (World’s End) populations of *P. leucopus* in the Boston Harbor. **C** and **D** reflect comparisons between island (Pender and Saturna) and mainland (Maple Ridge) populations of *P. maniculatus* in the Gulf Islands. X-axis categories represent three partitions of variation segregating between each island-mainland pair: (left) SNPs that are found in both populations, (middle) SNPs that are only found in the island population, and (right) SNPs that are only found in the mainland population. Colors correspond to five genic mutation types: introns (red), 3’ UTRs (green), 5’ UTRs (blue), synonymous sites (yellow), and nonsynonymous sites (orange). Bar heights reflect the proportion of SNPs within each mutation type that fall within each partition. Black horizontal lines denote the size of each partition observed for intergenic SNPs.

Differences between intergenic and genic elements in their relative proportions of shared versus private variation highlight the impact of purifying selection. Compared to intergenic mutations, nonsynonymous mutations are comprised of a higher proportion of island-private or mainland-private SNPs and a lower proportion of shared SNPs (Figure 3). This pattern is consistent with strong purifying selection in the island-mainland ancestor, where selection against nonsynonymous changes reduces their frequencies in the ancestral population such that island and mainland populations are less likely to “sample” overlapping subsets of these variants following their split. There is a larger discrepancy between the level of shared variation among intergenic and genic SNPs in Boston Harbor *P. leucopus* (Figure 3A-B) compared to Gulf Islands *P. maniculatus* (Figure 3C-D). This difference, coupled with mutation-type specific dissimilarities in the SFS between the two species (Supplementary Figure S1), suggests that the impacts of linked selection extend over greater genomic distances in *P. leucopus*, possibly due to differences in recombination rate between the species.

### Reconstruction of the joint DFE reveals differences in the strength of selection across mutation types

As illustrated in the previous section, patterns of variation within and between populations yield qualitative insights about the relative strength of purifying selection across genic mutation types. The distribution of fitness effects (DFE), by contrast, provides a quantitative, model-based portrait of how selection shapes the dynamics of fitness-affecting mutations. Drawing inferences about the DFE in island populations is complicated by two major characteristics of island evolution. First, divergence between island and mainland populations involves asymmetric reductions in effective population size that are most extreme on islands, making it difficult to disentangle the effects of increased genetic drift (captured by *N_e_*) from the effects of selection (captured by *s*). Second, most population genetic frameworks for DFE inference assume that selection has remained constant over the lifespan of a mutation. This assumption is likely violated on islands, where island-mainland divergence coincides with major environmental shifts that may alter the fitness effects of shared mutations. In other words, mutations that carry a given selective effect in the ancestral mainland environment (*s*_mainland_) may produce distinct effects on fitness when “tested” on islands (*s*_island_).

To reconstruct mutational fitness effects while accommodating these features of island populations, we employed a “joint DFE” framework (Huang et al., 2021). We fit bivariate models of selection to observed allele frequencies of fitness-affecting mutations segregating between island and mainland populations. Joint DFE models are characterized by a marginal distribution (which is either distinct or shared between populations) that represents the distribution from which *s* is drawn, as well as a correlation parameter that measures the similarity of *s*_island_ and *s*_mainland_ and for shared mutations (Supplementary Figure S2). We assume that deleterious selection coefficients follow a bivariate lognormal distribution with means μ_1_ and μ_2_, standard deviations σ_1_ and σ_2_, and correlation ρ. To account for the possibility that DFEs for certain mutation types are enriched for neutral or beneficial mutations, we additionally fit joint DFE models comprised of the aforementioned continuous deleterious distribution (*s* < 0), plus a discrete point mass of neutral (*s* = 0) or weakly beneficial (*s* > 0) mutations with proportions *P*_neu_ and *P*_pos_, respectively (see Supplementary Figure S2 and Materials and Methods for details). For each system and each genic mutation type, we separately fit these joint DFE models to variation segregating among island and mainland populations.

Joint DFE models involving distinct marginal distributions, where the “shape” of the DFE differs between island and mainland (μ_1_ ≠ μ_2_ and σ_1_ ≠ σ_2_), failed to achieve parameter convergence for any mutation type (Supplementary Figure S3H-N; Supplementary Figure S4H-N). As a result, we proceeded with joint DFE parameterizations where mutational fitness effects on the island and mainland are drawn from identical marginal distributions (μ_1_ = μ_2_ and σ_1_ = σ_2_). Most mutation types were best fit by continuous DFE models, with the exception of nonsynonymous, 3’ UTR, and 5’ UTR mutations in Gulf Islands *P. maniculatus*, which were best fit by DFE models that include a discrete neutral point mass. Likelihood ratio tests did not favor DFE models that included a point mass of beneficial mutations, suggesting that the genic mutation types analyzed here are not enriched for segregating beneficial mutations (Supplementary Table S1). Model fits and parameter estimates obtained under best-fitting joint DFE models for each mutation type and island-mainland comparison are presented in Supplementary Table S2 and Supplementary Figures S5-S8.

Across species and island-mainland comparisons, nonsynonymous mutations show the greatest selective constraint, with inferred median |*s*| (weighted by *P*_neu_ for Gulf Islands *P. maniculatus*) ranging from 4.35 x 10^-5^ to 6.13 x 10^-5^, followed by 5’ UTR mutations (6.63 x 10^-6^-2.24 x 10^-5^), and 3’ UTR mutations (3.42 x 10^-7^-5.73 x 10^-6^) (Figure 4; Supplementary Figure S9; Supplementary Table S2). Synonymous and intronic sites are the most enriched for neutral mutations, with median |*s*| ranges of 1.89 x 10^-10^-7.59 x 10^-9^ and 1.24 x 10^-11^-9.72 x 10^-10^, respectively (Figure 4; Supplementary Figure S9; Supplementary Table S2). Across genic mutation types, median |*s*| is negatively correlated with nucleotide diversity (π), Tajima’s *D*, and island-mainland F_ST_ measured across the corresponding genic elements (Supplementary Figures S10-S11).

**Figure 4.**
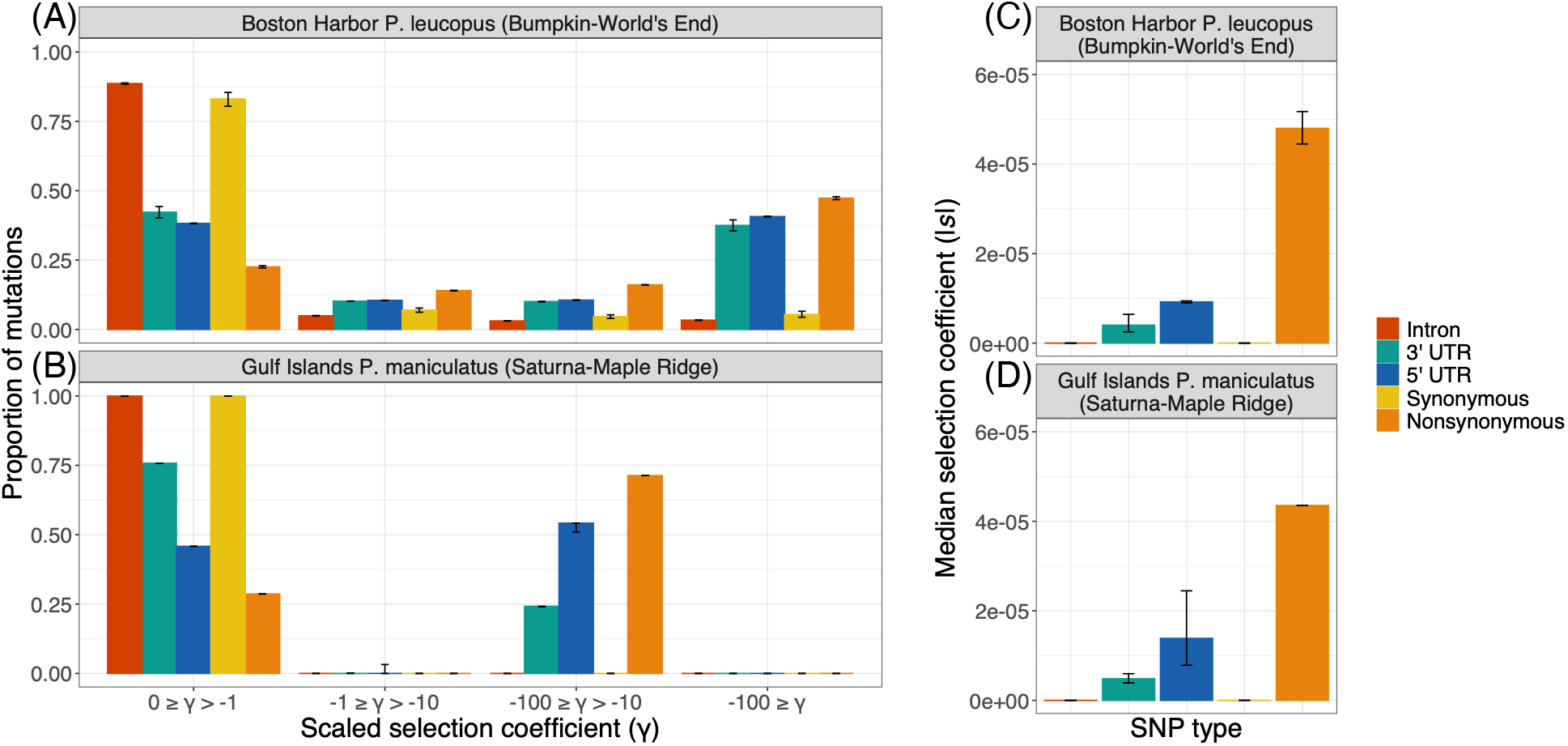
Marginal DFEs capture differences in selective constraint across genic mutation types. Panels **A** and **B** depict discretized versions of the shared marginal DFE inferred from representative island-mainland comparisons of Boston Harbor *P. leucopus* (Bumpkin Island versus mainland World’s End; **A**) and Gulf Islands *P. maniculatus* (Saturna Island versus mainland Maple Ridge; **B**). For mutation types best-fit by a DFE with a neutral point mass, discretized plots are weighted to include a proportion *P*_neu_ of mutations with *s* = 0 and a proportion 1 - *P*_neu_ of mutations with *s* < 0. X-axes denote binned population scaled selection coefficients (γ=2*N_Anc_s*) ranging from neutral/weakly deleterious (left-most bins) to more strongly deleterious (right-most bins). Bar colors reflect different mutation types. Bar height indicates the proportion of mutations falling into a given bin of γ based on the inferred DFE parameters. For a given mutation type, error bars reflect uncertainty in the estimated μ of the lognormal distribution (see Materials and Methods for details). Panels **C** and **D** summarize the central tendency of the distributions in **A** and **B** by the (absolute) median selection coefficient, *s*. Corresponding marginal DFEs for other island-mainland comparisons of Boston Harbor *P. leucopus* and Gulf Islands *P. maniculatus* are presented in Supplementary Figure S9.

### Divergence in mutational fitness effects mirrors environmental divergence

The joint DFE framework provides a rare opportunity to measure the similarity of fitness effects of shared mutations between island and mainland populations following their divergence in distinct environments. For nonsynonymous mutations, joint DFE correlations across all island-mainland comparisons are significantly less than 1 (Figure 5A), indicating that fitness effects of this constrained class of mutations have diverged between island and mainland populations since their split from a shared ancestor. The divergence in fitness effects is especially strong for *P. leucopus* in Boston Harbor, where point estimates of the joint DFE correlation range from ρ = 0.11 to ρ = 0.21. Fitness effects are more strongly correlated, but still divergent, between island and mainland populations of *P. maniculatus* in the Gulf Islands, where point estimates of the joint DFE correlation range from ρ = 0.61 to ρ = 0.65.

**Figure 5.**
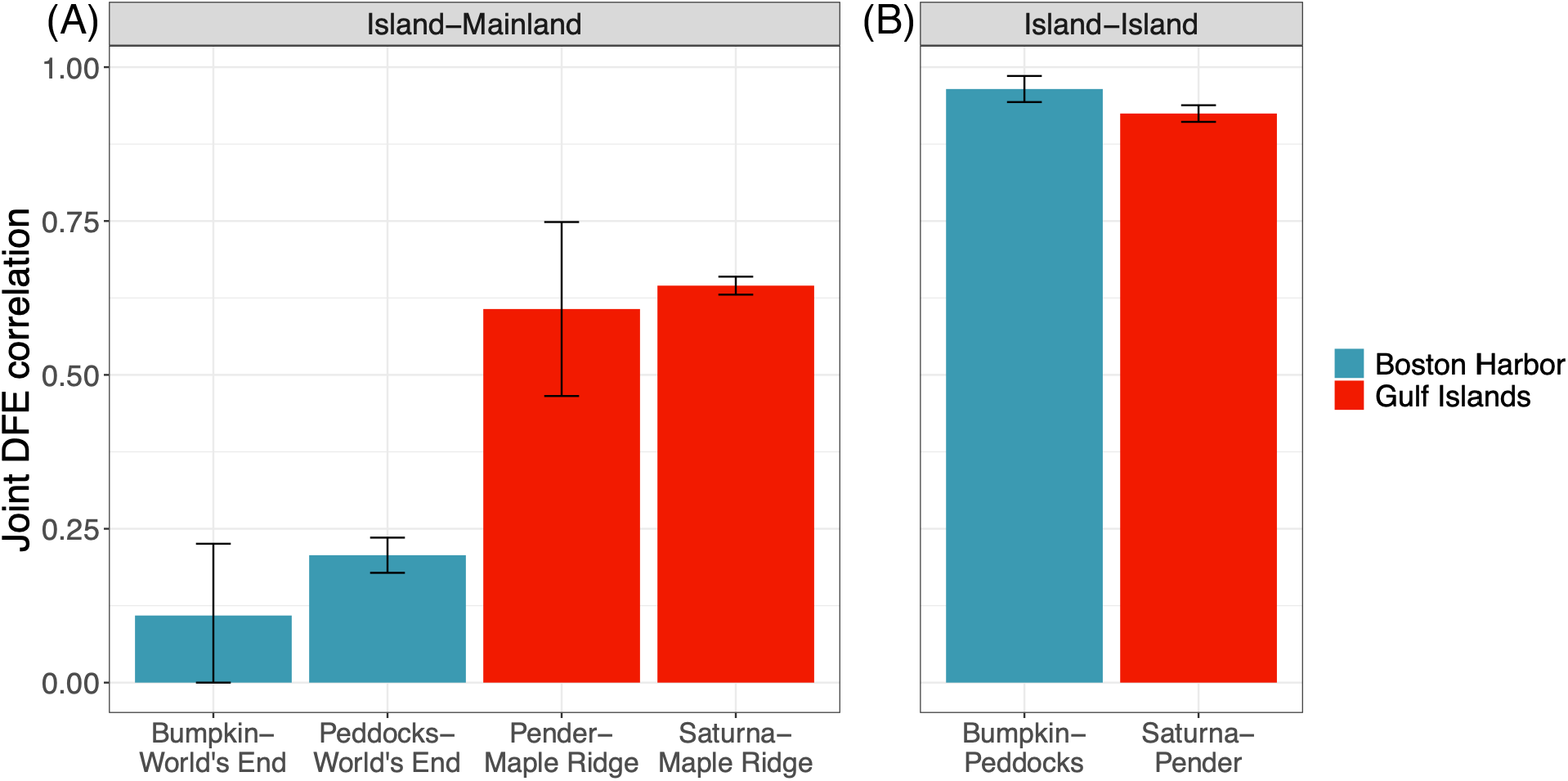
Fitness effects at nonsynonymous mutations are more strongly correlated between island populations. Panel **A** compares the strength of the joint DFE correlation measured across island-mainland contrasts in Boston Harbor *P. leucopus* (blue) and Gulf Islands *P. maniculatus* (red) for nonsynonymous mutations. Y-axis measures the magnitude of the correlation. X-axis denotes the population pair to which the joint DFE was fit. Panel **B** depicts the joint DFE correlation for models fit to nonsynonymous mutations segregating in island-island population pairs. Error bars depict +/-1 standard deviation around each point estimate of ρ.

To better understand the relationship between the joint DFE correlation and environmental divergence, we leveraged additional *within*-species comparisons of island populations, which we expect to experience more similar environments to each other than to their mainland counterparts. To draw this contrast between island-mainland and island-island comparisons, we additionally fit joint DFE models to nonsynonymous variants segregating among each pair of island populations within the Boston Harbor and Gulf Islands systems. Strikingly, there are major differences in the magnitude of the joint DFE correlation between island-island and island-mainland comparisons (Figure 5; Supplementary Table S2). Across both systems, the joint DFE correlation between islands (ranging from ρ = 0.92 to ρ = 0.96) is significantly higher than those inferred for corresponding island-mainland comparisons (Figure 5B). Moreover, median |*s*| of the marginal distributions for island-island comparisons are also higher than those inferred from island-mainland comparisons (Supplementary Figure S12). Collectively, these results indicate that differences in the selective regime conferred by island versus mainland environments are visible in genome-wide patterns of variation.

## DISCUSSION

### Divergence between island and mainland populations in the fitness consequences of mutations

The experimental measurement of mutant fitness across distinct environments has been employed for diverse purposes, from testing predictions of Fisher’s geometric model of adaptation (Hietpas et al., 2013) to predicting collateral sensitivity and resistance in human pathogens (Ardell & Kryazhimskiy, 2021). Deep mutational scanning (Fowler et al., 2010; Hietpas et al., 2012) and other emerging high-throughput techniques for quantifying mutant fitness enable fresh examination of the relationship between genotype, environment, phenotype, and fitness at high resolution. Fewer studies have investigated the consequences of mutations across distinct environments in a broader ecological context, where the variety of selective pressures that differ between environments and the specific genetic changes that contribute to differential fitness are unknown. Theoretical examination of the joint DFE in the context of this “divergent optima” model suggests that mutational fitness correlations between environments depend on both the distance of the “wild type” from each environment’s optimum and how closely aligned these distinct optima are in phenotypic space (Martin & Lenormand, 2015).

Under this interpretive framework, the high island-island fitness correlations we infer within each system are consistent with close alignment between islands in the position of their optima (Martin & Lenormand, 2015). In other words, segregating mutations elicit fewer trade-offs in fitness between island environments than between island and mainland. Though ecological idiosyncrasies vary across islands, there is abundant evidence to suggest that islands are more similar to each other than to their mainland counterparts. In both systems, islands deviate from the mainland in their composition of predator and interspecific competitor species, each of which are thought to be important drivers of patterns of phenotypic evolution on islands (Foster, 1964; Case, 1978; Lomolino, 1985; Adler & Levins, 1994; Palkovacs, 2003; Lomolino, 2005). Detailed field surveys of islands in the Boston Harbor support ephemeral incidences of large mammalian predators, like coyotes, and variable populations of other predator and competitor species, such as foxes, mink, voles, and squirrels (Nolfo-Clements, 2018). Though no systematic species survey has been conducted on the Gulf Islands, citizen scientists report a lower predator richness on islands (Berg & Nietlisbach, 2025), and multiple field observations describe an absence of mainland competitors, such as chipmunks, squirrels, and voles, on specific islands (Halpin & Sullivan, 1978; Berg & Nietlisbach, 2025).

This theoretical model of the joint DFE also predicts that the correlation in mutational fitness effects will be low when the wild type is well-adapted to one environment and poorly adapted to the other (Martin & Lenormand, 2015). Assuming that mice are well-adapted to their ancestral mainland environments, one interpretation for the lower island-mainland fitness correlation we estimated in Boston Harbor *P. leucopus* is that the “mainland phenotype” is more deleterious on islands in this system compared to Gulf Islands *P. maniculatus*. Though we lack a framework for comparing the degree and mode of island-mainland environmental divergence between systems, one prominent difference concerns island size: the Gulf Islands are over an order of magnitude larger in area than islands in the Boston Harbor. Biogeographical principles predict that larger islands may retain more of their mainland character than smaller, more isolated islands (MacArthur & Wilson, 1967a; Adler et al., 1986; Adler & Levins, 1994). In addition, the larger sizes of the Gulf Islands have likely buffered against human impacts in these urban-proximal systems. In the Boston Harbor, these impacts of human exploitation have had severe and lasting consequences. Dramatic alteration of the natural landscape for agricultural, industrial, and military usage have reshaped present-day vegetation, with over 50% of island flora comprised of non-native plant species (Elliman, 2005; Richburg & Patterson, 2005). Together, this more pervasive and extreme environmental restructuring may contribute to greater mutational trade-offs between island and mainland in the Boston Harbor.

Although our findings provide strong evidence for divergence in the fitness effects of nonsynonymous mutations across environments, support for distinct “marginal” DFEs between island and mainland populations is equivocal. The poorer fit of models featuring population-specific DFEs may indicate that the overall “shape” of the DFE is conserved across populations or reflect the challenges of fitting increasingly complex models to real data. Conservation of the DFE has been observed at multiple levels of biological organization, including across populations of the same species in forest trees (James et al., 2023), and among closely related species in primates (Castellano et al., 2019; Sendrowski et al., 2026). Studies of the DFE across a much broader phylogenetic scope suggest that there is strong phylogenetic signal in DFE parameters (Lin et al., 2025), and that variation in the DFE is likely driven by higher order properties such as long-term effective population size and organismal complexity (Huber et al., 2017; Lin et al., 2025).

### Distinct levels of selective constraint across mutation types

Estimated strengths of selection acting on each genic mutation type are similar in *P. leucopus* and *P. maniculatus*, a finding that supports the robustness of our inferences. Among the mutation types considered here, nonsynonymous variants carry the strongest fitness consequences, consistent with the negative impacts that nonsense, missense, and frameshift mutations have on protein sequence and folding. 5’ UTR mutations in both species exert stronger selective effects than those in 3’ UTRs. Both untranslated regions play key roles in the post-transcriptional regulation of gene expression. As the site of translation initiation, 5’ UTRs exert translational control and modulate translational efficiency, whereas 3’ UTRs affect mRNA stability, degradation, and localization (Mignone et al., 2002). Comparisons of substitution rates across mammalian genes have found greater sequence conservation at 5’ UTRs compared to 3’ UTRs (Pollard et al., 2010). In addition, 5’ UTRs exhibit greater uniformity in their average length across diverse eukaryotic taxa (Mignone et al., 2002). Together, these observations support stronger selective constraint on 5’ UTRs compared to their downstream counterparts.

The DFEs we reconstructed also support the notion that synonymous and intron variants are largely neutral. Despite their common use as a neutral proxy in population genetic inferences based on exome capture sequencing (Ragsdale et al., 2018; Martinez i Zurita et al., 2025), experimental measurement of deleterious fitness effects of synonymous mutations (Shen et al., 2022; Zhang & Qian, 2025), paired with evidence of codon usage bias across diverse taxa (reviewed in Hershberg & Petrov, 2008), indicate that a non-trivial proportion of these mutations may experience selection (Zhang & Qian, 2025). Sequence variation in introns may also affect fitness by altering or disrupting RNA splicing and through modifications to the regulatory elements that reside within them (Majewski & Ott, 2002). Despite this potential for direct selection on intronic and synonymous variation (as well as linked selection driven by their proximity to highly constrained nonsynonymous sites), we find little evidence for widespread purifying selection on these two classes as a whole. Further partitioning of these two variant classes by their likely functional consequence could reveal selection acting on specific subsets of mutations. For example, stratifying synonymous changes by effective codon number (Wright, 1990) or by gene length may enable estimation of the strength of selection on translational efficiency (Comeron et al., 1999). In addition, separating intronic changes into those affecting the first intron or those overlapping splice control regions could provide additional insights into the nature of purifying selection acting on these gene elements (Keightley & Gaffney, 2003; Comeron, 2004; Halligan et al., 2004).

### Limitations of the joint DFE

Because the joint DFE correlation shapes the amount of shared polymorphism between populations, inference of this parameter depends on the timescale of divergence between populations. Huang et al. (2021) reported that the variance in the estimated joint DFE correlation can be high both for populations that diverged very recently, where there is insufficient time for selection to act differently, and for populations that diverged long ago, where there is insufficient shared variation for inference. It is less well-known how the temporal scope of the joint DFE correlation may be affected by the selective character of variation shared between populations. For example, inferences about the DFE may be biased towards the weaker-effect mutations that comprise the bulk of segregating variation, as highly detrimental mutations are unlikely to be observed in a sample of polymorphism (Eyre-Walker & Keightley, 2007). In an analogous manner, inferences about the joint DFE (and the joint DFE correlation) are circumscribed to the variation shared between populations, which will change as populations diverge. In the Gulf Islands, where island and mainland lineages diverged longer ago, mutations with highly divergent effects on fitness have likely been removed from shared polymorphism, enriching shared mutations for those with more similar effects on fitness. Such temporal “bias” could provide an alternative explanation for the stronger joint DFE correlation inferred for nonsynonymous mutations in this system compared to more recently diverged island and mainland populations in the Boston Harbor.

Our approach to modeling selective divergence between island and mainland populations also assumes that mutations act in an additive manner. In reality, we expect dominance coefficients to be both dynamic (e.g., influenced by genetic background or environment) and highly variable across mutations (reviewed in Di & Lohmueller, 2024). Under departures from demographic equilibrium, dominance coefficients play an important role in the dynamics of deleterious variation (Balick et al., 2022). For populations segregating a large proportion of recessive deleterious mutations, extreme bottlenecks increase the proportion of homozygous genotypes, leading to more effective “purging” of these mutations (Hedrick & Garcia-Dorado, 2016; Kyriazis & Lohmueller, 2024). Given the large asymmetries in the *N_e_* reductions experienced by island and mainland populations in both systems (Howell et al., 2025a; Howell et al. 2025b), it is possible that the observed disparity in the strength of selection we infer for nonsynonymous mutations in island-island versus island-mainland contrasts may reflect a greater deficit of nonsynonymous mutations on islands due to more effective purging of recessive deleterious mutations.

### Prospects for empirical study of the joint DFE

Although the joint DFE framework employed here enabled robust estimation of the distribution of *s* across mutation types, the joint DFE correlation was more difficult to infer. That the ability to estimate this correlation was restricted to nonsynonymous variants (the most constrained class of mutations) suggests challenges with identifiability for weakly-selected mutation types. In the context of demographic inference, distinct population size histories have been found to give rise to identical SFS (Myers et al., 2008; Baharian & Gravel, 2018), revealing issues with identifiability for frameworks that rely on the SFS. Similar principles may apply to DFE inference–especially in the context of multiple populations–where information about natural selection is contained in both marginal and joint allele frequencies. Though this lack of identifiability precludes a comparison of how fitness effect correlations vary across mutation types, it is possible that prioritizing mutations in these weakly-selected classes that are most likely to affect fitness (e.g., intronic mutations within splice control regions or synonymous mutations with low effective codon number) could enable robust estimation and comparison of this parameter across functional classifications.

Laboratory comparisons in house mice have yielded rich insights into the specific genomic intervals, pathways, and genes that contribute to morphological, physiological, and behavioral differences between island and mainland populations (Gray et al., 2015; Parmenter et al., 2016; Nolte et al., 2020; Wilches et al., 2021; Parmenter et al., 2022; Payseur et al., 2023; Stratton et al., 2023). These comparisons offer additional biologically relevant partitions across which the joint DFE can be estimated and contrasted, providing opportunities to query fitness effect correlations among variants located in genes suspected to be connected to phenotypic evolution on islands. In addition, exploration of the joint DFE in systems where differences between island and mainland environments have been surveyed in greater detail present possibilities for identifying the ecological factors that drive divergence in fitness effects. Recent advances in scalable DFE inference (Sendrowski & Bataillon, 2024) provide an efficient framework for identifying covariates of DFE parameters. Future integration of such software with the underlying joint DFE model proposed by Huang et al. (2021) could empower statistical testing of how such ecological variables (e.g., predator and competitor abundance) modulate fitness effect correlations between island and mainland populations.

## CONCLUSION

Our results highlight the capacity of the joint DFE framework to quantify divergence in fitness effects between populations adapting to distinct environments and undergoing demographic changes. By extending this framework to multiple island radiations, we provide a rare empirical demonstration that divergence in selective regimes in nature is recorded in genome-wide patterns of variation. Extension of this framework to other island-mainland systems presents exciting opportunities to connect divergence in mutational fitness effects to the specific ecological variables and genetic changes that distinguish island and mainland populations.

## MATERIALS AND METHODS

### Population genomic dataset construction

We analyzed the whole genome, short-read population genomic callsets generated by Howell et al. (2025a) and Howell et al. (2025b) for *P. leucopus* sampled from Massachusetts’ Boston Harbor and *P. maniculatus* sampled from the Gulf Islands of British Columbia. This encompasses the subset of unrelated (≥ 3^rd^ degree) individuals sampled from each island and mainland locale. For Boston Harbor *P. leucopus*, unrelated sample sizes are n=13 for Bumpkin Island, n=13 for Peddocks Island, and n=17 for mainland World’s End. For Gulf Islands *P. maniculatus*, unrelated sample sizes are n=20 for Saturna Island, n=10 for Pender Island, and n=17 for mainland Maple Ridge. All samples were sequenced to a target coverage of 30X. Details regarding population sampling, whole genome sequencing, alignment, and variant calling are described in Howell et al. (2025a) and Howell et al. (2025b) for *P. leucopus* and *P. maniculatus* datasets, respectively. Sample identifiers and corresponding NCBI SRA BioSample accessions for genomes analyzed in this study are provided in Supplementary Table S3.

To standardize variant callsets between the two species, we refiltered the raw “joint” variant calls produced by running GATK (v4.2.0.0) GenotypeGVCFs within each population (i.e., separately for Bumpkin, Peddocks, World’s End, Saturna, Pender, and Maple Ridge cohorts). Starting from these unfiltered, single-population callsets, we filtered variant calls with GATK VariantFiltration using site-level annotation thresholds based on GATK Best Practices recommendations for hard filtering (“FisherStrand” > 60.0, “StrandOddsRatio” > 3.0, “RMSMappingQuality” < 40.0, “MappingQualityRankSumTest” < -12.5, and “ReadPosRankSumTest” < -8.0; McKenna et al., 2010; Auwera & O’Connor, 2020) with modifications (“QualByDepth” < 5.0). Repetitive DNA can manifest as regions of high heterozygosity when repeat-derived reads map to the same position. To mitigate this issue, we additionally used GATK VariantFiltration to remove sites with “ExcessHet” > 15.0, corresponding to a p-value less than 0.05 for the Hardy-Weinberg exact test of excess heterozygosity (Wigginton et al., 2005). We then used GATK SelectVariants to remove filtered sites and restrict each callset to autosomal, biallelic, single-nucleotide variants.

Jointly calling variants within (rather than among) populations of a species can yield incongruent callsets when there is sufficient differentiation between them, as variants private to one cohort will not be represented among the variant calls of other cohorts. This creates ambiguity with respect to variants that are “missing” between callsets, which reflect sites that are either monomorphic for the reference allele within a given cohort or are low-confidence genotype calls that were not emitted. To ensure that joint allele frequencies within each species dataset were properly calibrated for downstream analyses, we “rescued” genotypes at these missing sites separately for Boston Harbor *P. leucopus* and Gulf Islands *P. maniculatus*. Within each species, we used the bcftools (v1.8) (Danecek et al., 2011) “isec” function to extract the locations of private variants observed in each population-specific call set. For each collection of private variants, we used the “-all-sites” mode of GATK GenotypeGVCFs with the “-stand-call-conf” threshold set to 0 to emit monomorphic reference calls at these private variant sites for each of the other cohorts in which they were not observed. We used GATK VariantFiltration and SelectVariants to apply the same site-level filtering thresholds to these “rescued” calls (excluding “QualByDepth”). The resulting high-quality “invariant” calls were then combined with the polymorphic calls of their respective cohort and the resulting monomorphic + polymorphic callsets were merged across cohorts. This procedure yielded a single, multi-population callset for each species that represents the union of variant sites observed across the three populations that comprise each species dataset. We retained 80,504,816 SNPs in the Boston Harbor *P. leucopus* dataset and 125,459,727 SNPs in the Gulf Islands *P. maniculatus* dataset following site-level variant filtering and callset merging.

To obtain high-confidence SNP callsets for downstream population genomic analyses, we conducted additional filtering of each multi-population callset using two individual-level annotations: coverage depth (DP), which measures individual read count at a given SNP, and genotype quality score (GQ), which measures the difference between the second lowest and the lowest phred-scaled genotype likelihood. Since both low coverage (indicating poor support for an individual’s genotype call) and inflated coverage (which may reflect copy number variation that is poorly resolved in the reference assembly) can lead to erroneous variant calls (Li, 2014; Pfeifer, 2017), we set upper and lower bounds on coverage depth at each site. Based on thresholds established in Howell et al. (2025b), we use GATK VariantFiltration and SelectVariants to remove sites in each multi-population callset for which any individual failed the following filters: “DP” > 8, “DP” < 52, and “GQ” > 25. To ensure that joint allele frequencies could be properly measured at each SNP, we additionally removed sites in each multi-population callset that contained missing genotype information for one or more individuals. We retained 98,430,235 SNPs in the Gulf Islands *P. maniculatus* callset and 60,790,545 SNPs in the Boston Harbor *P. leucopus* callset following these rigorous individual-level filtering procedures.

We also restricted downstream analyses to variants falling outside of annotated repetitive elements and low-complexity DNA, which can cause errors in read mapping and variant calling (Pfeifer 2017). For each species, we removed SNPs falling within regions annotated by the RepeatMasker (https://repeatmasker.org), WindowMasker (Morgulis et al., 2006), and Simple Repeats (Benson, 1999) tracks of the UCSC Genome Browser (http://genome.ucsc.edu; Karolchik, 2004). The corresponding names of these tracks are hub_6477253_repeatMasker, hub_6477253_windowMasker, and hub_6477253_simpleRepeat in *P. leucopus* and hub_6502785_repeatMasker, hub_6502785_windowMasker, and hub_6502785_simpleRepeat in *P. maniculatus. P. maniculatus* are also known to harbor an abundance of large inversion polymorphisms (Hager et al., 2022; Harringmeyer & Hoekstra, 2022) that cause major distortions in local patterns of variation (Howell et al. 2025b). Because inversions can experience distinct recombination and selective landscapes, SNPs falling within these intervals may also violate the assumptions of the SFS-based inference frameworks we employ here. To mitigate these issues, we excluded *P. maniculatus* SNPs falling within the breakpoints of the 21 inversion polymorphisms defined by Harringmeyer and Hoekstra (2022). After excluding sites falling within annotated repeats and inversions (for *P. maniculatus*), we retained 18,377,786 SNPs in our Boston Harbor *P. leucopus* dataset and 8,971,514 SNPs in our Gulf Islands *P. maniculatus* dataset.

### SNP polarization

To gain increased resolution for our SFS-based analyses, we polarized SNPs in both our *P. leucopus* and *P. maniculatus* callsets, which enables us to distinguish between high and low-frequency derived variation. To determine ancestral states at polymorphic sites, we employed the approaches of Keightley et al. (2016) and Keightley & Jackson (2018) as implemented in the *est-sfs* program. In contrast to maximum parsimony-based methods, which use comparisons of single representative genomes for both focal and outgroup species, *est-sfs* leverages up to three outgroups and uses polymorphism information (i.e., nucleotide counts at each SNP) in the focal species to estimate per-site ancestral state probabilities.

This method assumes that observed variation coalesces within the focal species and is not share with outgroups. To guide the selection of appropriate outgroup species, we conducted whole-genome alignments of all *Peromyscus* NCBI RefSeq assemblies available at the time of this analysis: *P. maniculatus* (HU_Pman_2.1.3; GCF_003704035.1), *P. polionotus* (HU_Ppol_1.3.3; GCA_003704135.2), *P. leucopus* (UCI_PerLeu_2.1; GCF_004664715.2), *P. californicus* (ASM782708v3; GCF_007827085.1), *P. eremicus* (PerEre_H2_v1; GCF_949786415.1), *P. nudipes* (Pnud_10x_v1; GCA_902168325.1), *P. melanophrys* (Pmel_10x_v1; GCA_902168415.1), *P. attwateri* (Patt_10x_v1; GCA_902168425.1), and *P. aztecus* (Pazt_10x_v1; GCA_902168405.1). Prior to aligning these nine *Peromyscus* species, we “soft-masked” repetitive elements in each assembly with RepeatMasker (v4.1.7) (http://www.repeatmasker.org) using the Dfam (v3.8) database of transposable elements (Hubley et al., 2016) and the RMBlast engine. Across all species, RepeatMasker masked between 33.06% and 40.38% of the genome. We conducted whole-genome alignments of these soft-masked assemblies using progressive Cactus (v2.9.3) (Armstrong et al., 2020). This approach makes use of a bifurcating “guide tree” that determines the order in which the input sequences and reconstructed ancestral sequences should be aligned. To construct this guide tree, we used mashtree (v1.4.6) (Katz et al., 2019) to estimate pairwise distances between soft-masked *Peromyscus* assemblies. The resulting distance-based tree (Supplementary Figure S13) recovers previously characterized phylogenetic relationships by Bradley et al. (2007) and Platt et al. (2015). Using this guide tree and the corresponding soft-masked *Peromyscus* assemblies as input, we ran progressive Cactus on its default settings to generate a whole-genome, multiple sequence alignment in the Hierarchical Alignment (HAL) format (Hickey et al., 2013).

To mitigate the impact of incomplete lineage sorting on SNP polarization in *P. leucopus* and *P. maniculatus*, we constructed a set of three outgroups for each of these focal species by selecting a single representative species from each of the more distantly related *P. californicus-P. eremicus*, *P. melanophrys-P. nudipes*, and *P. aztecus-P. attwateri* clades (Supplementary Figure S13). We used the halSnps utility of the HAL toolkit (v2.2) (Hickey et al. 2013) to output nucleotide states of each potential outgroup species at each polymorphic site in our *P. leucopus* and *P. maniculatus* callsets. To identify optimal outgroup combinations, we compared their “coverage” of each focal species based on the fraction of sites in each callset for which outgroup states could be identified. The results of this coverage analysis (depicted in Supplementary Figures S14-S15) suggested that *P. californicus*, *P. melanophrys*, and *P. attwateri* comprise the optimal three-outgroup set for both *P. leucopus* and *P. maniculatus*. Collectively, this outgroup combination covered 79% and 76% of sites in our *P. leucopus* and *P. maniculatus* callsets.

We then used *est-sfs* to estimate ancestral state probabilities separately for our *P. leucopus* and *P. maniculatus* datasets using nucleotide states from each of the selected outgroup species and combined allele counts at each SNP in our multi-population callset. We ran *est-sfs* (v2.04) with the following specifications in the configuration file: “n_outgroup 3”, “model 1” and “nrandom 10”, which instruct the program to use three outgroup species, the Kimura two-parameter model of nucleotide substitution, and ten random starting value runs for maximum likelihood estimation. For each SNP in the focal species, *est-sfs* returns a per-site probability that the major allele is ancestral. We assigned ancestral states in a probabilistic manner for each SNP by modeling a single draw from a binomial distribution with probability of success equal to the ancestral state probability output by *est-sfs* (following Beichman et al., 2023). For “successes”, we assigned the major allele as the ancestral state; otherwise, we assigned to major allele as the derived state.

### Classification of mutation types and functional elements

Genetic variation analyzed in this study can be partitioned into “intergenic” SNPs– which are assumed to behave in a neutral manner– and “genic” SNPs– which we assume could be experiencing direct or linked selection. Our genic category is comprised of SNPs falling within NCBI RefSeq protein-coding genes. We further classify genic SNPs into five mutation types: those falling in 5’ UTRs, 3’ UTRS, introns, synonymous exon variants, and nonsynonymous exon variants. To construct these genic partitions, we used the Ensembl Variant Effect Predictor (VEP) program (McLaren et al., 2016) to generate variant predictions for SNPs observed in each multi-population callset. For each species, we ran VEP (v114.0) using the corresponding multi-population callset, NCBI RefSeq assembly (UCI_PerLeu_2.1/GCF_004664715.2 for *P. leucopus*; HU_Pman_2.1.3/GCF_003704035.1 for *P. maniculatus*) and associated GFF annotation file as inputs. Supplementary Table S4 summarizes variant consequences predicted by VEP for Boston Harbor *P. leucopus* and Gulf Islands *P. maniculatus* SNPs.

We classify these variant effect predictions into five genic mutation types (introns, 5’ UTR, 3’ UTR, synonymous, and nonsynonymous) based on the Sequence Ontology hierarchy (Eilbeck et al., 2005; Cunningham et al., 2015). Intron variants include SNPs annotated as “intron_variant” (SO:0001627), “splice_donor_variant” (SO:0001575), or “splice_acceptor_variant” (SO:0001574). 5’ UTR variants are those annotated as “5_prime_UTR_variant” (SO:0001623). 3’ UTR variants are those annotated as “3_prime_UTR_variant” (SO:0001624). Synonymous variants include SNPs annotated as “synonymous_variant” (SO:0001819), “stop_retained_variant” (SO:0001567), or “start_retained_variant” (SO:0002019). Nonsynonymous variants include SNPs annotated as “nonsynonymous_variant” (SO:0001992), “stop_lost” (SO:0001578), “start_lost” (SO:0002012), “missense_variant” (SO:0001583), or “stop_gained” (SO:0001587). Within each of these categories, we retain the subset of “high-confidence” SNPs that passed the variant filtering, masking, and polarization procedures described in the *Population genomic dataset construction* and *SNP polarization* sections. To account for instances where individual SNPs carried multiple predicted effects (e.g., due to transcript isoforms), we excluded SNPs that fell into multiple of the five genic mutation type classification described above. The resulting disjoint set of genic mutations comprised 7,358,310 intronic, 25,973 5’ UTR, 155,697 3’ UTR, 136,068 synonymous, and 62,425 nonsynonymous SNPs in the Boston Harbor *P. leucopus* callset and 4,394,533 intronic, 17,004 5’ UTR, 132,108 3’ UTR, 99,159 synonymous, and 37,786 nonsynonymous SNPs in the Gulf Islands *P. maniculatus* callset. Our intergenic category is comprised of the subset of SNPs located at least 1 kbp from annotated genes that passed the additional variant filtering, masking, and polarization procedures described in the *Population genomic dataset construction* and *SNP polarization* sections. This intergenic category amounted to 7,751,753 SNPs in our Boston Harbor *P. leucopus* callset and 3,304,122 SNPs in our Gulf Islands *P. maniculatus* callset.

Downstream DFE inference requires information about the effective sequence length (L) for each class of mutations, which represents the total length of “callable” sequence for a given category. To approximate L for each genic mutation type, we first used the UCSC Genome Browser Table Browser tool (http://genome.ucsc.edu; Karolchik et al 2004) to output BED-style records for introns, 5’ UTRs, 3’ UTRs, and coding exons in each species using the NCBI RefSeq gene annotation tracks (hub_6477253_ncbiRefSeq for *P. leucopus* and hub_6502785_ncbiRefSeq for *P. maniculatus*). We then restricted records to autosomal genes and used bedtools (v2.26.0) (Quinlan & Hall, 2010) to remove intervals of repetitive DNA (and inversions in *P. maniculatus*) that were masked from our SNP callset (described in the *Population genomic dataset construction* section). Effective sequence lengths for nonsynonymous and synonymous classes depend on the proportion of exon mutations that give rise to nonsynonymous versus synonymous changes. Following Huber et al. (2017), we assume a mammalian-like ratio of L_nonsyn_ = 2.31 x L_syn_, which they compute from estimates of the transition/transversion ratio and CpG mutational bias in humans. The resulting total sequence length for each mutation type (introns, 5’ UTRs, 3’ UTRs, nonsynonymous, and synonymous) was then multiplied by (1) the percent of SNPs retained after variant filtering (described in the *Population genomic dataset construction* section) and (2) the percent of SNPs that were retained after excluding variants with overlapping mutation type annotations (described above). Collectively, these operations capture the effect of our variant detection and annotation procedures on the amount of “callable” sequence corresponding to each genic mutation type. Similarly, we approximated L for the intergenic class by subtracting intervals representing annotated genes (plus 1 kbp of flanking sequence to mirror our ascertainment procedure), repetitive elements, and inversions (for *P. maniculatus*) from the total length of the autosomal genome in each species. As with genic mutation types, we then multiplied the resulting sequence length by the percent of SNPs retained after variant filtering to obtain the amount of callable sequence for intergenic mutations. Effective sequence lengths used for each mutation type in downstream analyses are summarized in Supplementary Table S5.

We characterized patterns of variation across each element type using the repeat-masked and (for *P. maniculatus*) inversion-masked intervals constructed above for autosomal introns, 5’ UTR, 3’ UTR, exons, and intergenic regions. Using our multi-population SNP callsets, we computed nucleotide diversity π (Tajima, 1983) and Tajima’s *D* (Tajima, 1989) within each *P. leucopus* and *P. maniculatus* population as well as F_ST_ (Hudson et al., 1992) between each island-mainland population pair across each element with scikit-allel v1.3.6 (https://github.com/cggh/scikit-allel).

### Re-fitting demographic models to the unfolded SFS

To ensure our demographic models were well calibrated to capture the effects of population history on allele frequency dynamics at genic SNPs, we re-fit the models inferred from the folded site frequency spectrum by Howell et al. (2025a) for Boston Harbor *P. leucopus* and Howell et al. (2025b) for Gulf Islands *P. maniculatus* to the polarized allele frequencies we obtained for intergenic variants. Using the high-confidence, intergenic callsets described above, we constructed 2D unfolded joint SFS for each island-mainland and island-island pair of *P. leucopus* and *P. maniculatus* populations. To estimate demographic parameters from these unfolded joint SFS, we used the maximum likelihood framework implemented in ∂a∂i (v2.4.3) (Gutenkunst et al., 2009). For each population pair, we re-fit the corresponding best-fitting 2D demographic models from Howell et al. (2025a) and Howell et al. (2025b) (described in Supplementary Figure S16). To account for potential errors in our SNP polarization, we augmented these models with an additional parameter, *p_misid_*, which represents the proportion of variants for which the ancestral state was misidentified. We used the BFGS optimization routine in ∂a∂i to obtain maximum likelihood estimates of model parameters. To ensure thorough exploration of the parameter space, starting parameter values were permuted across 10,000 independent searches. The resulting best-fit parameter values, which reflect the increased resolution provided by the polarized allele frequencies, are tabulated in Supplementary Table S6.

### Modeling the joint effects of demographic and selection on allele frequencies

The diffusion approximation employed by ∂a∂i is well-suited to accommodate a single selection coefficient, allowing one to model the joint effects of demography and selection on allele frequencies. Inference of the DFE, however, requires modeling the effect of demographic history and an arbitrary *distribution* of selection coefficients on the SFS. To overcome this challenge, the fit∂a∂i framework formulated by Kim et al. (2017) breaks the problem of DFE inference into multiple, computationally tractable steps. First, expected SFS are computed under a parameterized demographic model for a range of individual γ (where γ = 2*N_Anc_s*). Using these “cached” single-γ SFS, the expected SFS for a distribution of γ is computed by integrating over these cached spectra, weighting by the continuous distribution used to model the DFE. Then, using the same Poisson Random Field framework employed for demographic inference (Sawyer & Hartl, 1992; Gutenkunst et al., 2009), comparisons between this expected SFS (computed under a given DFE parameterization) and the observed SFS form the basis for maximum-likelihood estimation of DFE parameters (Kim et al., 2017).

Huang et al. (2021) extend this DFE framework to the case of two, recently diverged populations. Here, allele frequency dynamics at fitness-affecting variants reflect not only the joint demographic history of the two populations, but also the cumulative effects of selection acting in a shared ancestral population and (potentially) divergent selective regimes acting in contemporary populations following their split. This is achieved by allowing mutations with selective effect *s*_1_ in the ancestral population to assume a new selective effect *s*_2_ in the diverging population following its split. Modeling the impact of this joint distribution of selective effects (*s*_1_, *s*_2_) on joint allele frequencies requires “caching” 2D expected SFS under a 2D model of demographic history across a range of γ pairs (γ_1_, γ_2_).

Following the ∂a∂i/fit∂a∂i documentation (Gutenkunst et al., 2009; Kim et al., 2017; https://dadi.readthedocs.io/), to compute the expected joint SFS for a given γ_1_, γ_2_ pair, we modified the demographic functions corresponding to the models illustrated in Supplementary Figure S16 to include a “gamma=γ_1_” argument in the “dadi.PhiManip.phi_1D” function. This instantiates the population allele frequency spectrum, φ, with a fixed selective effect of 2*N_Anc_s*. Here, *s* is the selective advantage and genotypic fitnesses in the homozygous ancestral, heterozygous, and homozygous derived classes are given as 1, 1+2*sh*, and 1+2*s*. We assume that *h*=0.5 for all mutations, such that they act in a semi-dominant manner. For all island-mainland demographic models, we assume that selective effects may diverge immediately following the instantaneous split between populations. Following this split, subsequent manipulations of ϕ assign a “gamma=γ_1_” to the mainland population (i.e., selective effects remain constant) and a “gamma=γ_2_” to the island population. For island-island demographic models, we randomly choose which island population is assigned a “gamma=γ_1_” versus “gamma=γ_2_” following their split from a shared ancestor. We used the “dadi.DFE.Cache2D” function to generate expected joint SFS under these demographic + selection models for 100 logarithmically-spaced deleterious γ_1_, γ_2_ pairs within the range [-2000, -1×10^-4^]. In order to fit joint DFE models with point masses of neutral and beneficial mutations (see *Modeling the joint DFE* section below), we used the “additional_gammas” argument to cache expected joint SFS for an additional set of neutral and positive selection coefficients: γ_1_, γ_2_ ∈ [0, 1, 2, 3, 4, 5].

### Modeling the joint DFE

To draw inferences about the joint distribution of γ_1_, γ_2_, we assume that the joint DFE follows a bivariate lognormal distribution. We focus on this particular bivariate probability distribution because its marginal distribution has few parameters (μ and σ), it has an easily interpretable correlation parameter (ρ), and it permits the modeling of either shared (μ_1_ = μ_2_ and σ_1_ = σ_2_) or distinct (μ_1_ ≠ μ_2_ and σ_1_ ≠ σ_2_) marginal distributions. This allows us not only to estimate how strongly correlated selective effects of shared mutations are between island and mainland populations (captured by the magnitude of ρ), but also to test whether there is evidence for divergence between populations in the shape of the underlying distribution from which new selection coefficients are drawn (captured by the fit of distinct versus shared marginal parameters, μ and σ). To model the joint DFE as a bivariate lognormal distribution, we use the “dadi.DFE.PDFs.biv_lognormal” selection distribution function.

Tested joint DFE models include bivariate lognormal DFEs with distinct marginal distributions (μ_1_ ≠ μ_2_ and σ_1_ ≠ σ_2_) or shared marginal distributions (μ_1_ = μ_2_ and σ_1_ = σ_2_). Within these two different joint DFE parameterizations, we consider three different DFE types: (1) “deleterious-only” DFEs, where selection coefficients are continuously distributed according to a lognormal distribution with all γ < 0, (2) “deleterious + neutral” DFEs, where a proportion *P*_neu_ of mutations have a fixed γ = 0 and a proportion 1 – *P*_neu_ of mutations have a continuous, lognormal distribution of deleterious selection coefficients (γ < 0), or (3) “deleterious + beneficial” DFEs, where a proportion *P*_pos_ of mutations have a fixed γ ∈ [1, 2, 3, 4, 5] and a proportion 1 – *P*_pos_ of mutations have a continuous, lognormal distribution of deleterious selection coefficients (γ < 0). For the “deleterious + neutral” and “deleterious + beneficial” DFE types, we assume that the discrete point mass (*P*_neu_ or *P*_pos_) is symmetric between populations, and model them using the “dadi.DFE.Cache2D.integrate_symmetric_point_pos” function. For models involving a neutral point mass, the joint DFE is comprised of three different components: (1) variants that are neutral in both populations (with weight *p*_00_), (2) variants that are neutral in one population and deleterious in the other (with weight *p*_–0_), and (3) variants that are deleterious in both populations (with weight *p*_– –_). Building off the triallelic selection models employed by Ragsdale et al. (2016), these weights are computed as follows: *p*_00_ = *P*_neu_^2^ + ρ(*P*_neu_ – *P*_neu_^2^), *p*_–0_ = (1 – ρ) × *P*_neu_ × (1 – *P*_neu_), and *p*_– –_ = (1 – *P*_neu_)^2^ + ρ[1 – *P*_neu_ – (1 – *P*_neu_)^2^]. This ensures that when ρ = 0, these weights correspond to independent sampling of selective effects between the two populations. At the other extreme, when ρ = 0, these formulae ensure that no variant has divergent effects between the two populations. This same logic was applied for models involving a beneficial point mass (by replacing *P*_neu_ with *P*_pos_). Taking into account the five different fixed beneficial γ values used to model the “deleterious + beneficial” DFEs, we tested a total of 14 different joint DFE models across DFE parameterizations and DFE types. These distinct joint DFE models are graphically depicted in Supplementary Figure S2.

### Joint DFE model fitting and parameter estimation

Using the cached spectra described above, we fit each joint DFE model to unfolded joint SFS constructed from our high-confidence genic SNP callsets. For each island-mainland population pair in our Boston Harbor *P. leucopus* (Bumpkin-World’s End and Peddocks-World’s End) and Gulf Islands *P. maniculatus* (Saturna-Maple Ridge and Pender-Maple Ridge) datasets, we fit joint DFEs separately for each genic mutation type: introns, 5’ UTR, 3’ UTR, nonsynonymous, and synonymous. We additionally fit joint DFEs to nonsynonymous mutations for each island-island populations pair (Bumpkin-Peddocks for Boston Harbor *P. leucopus* and Saturna-Pender for Gulf Islands *P. maniculatus*). For each of the joint DFE models described above, we include an additional free parameter, *p_misid_*, which allows the proportion of variants for which the ancestral state was misidentified to vary across mutation types and population pairs.

In the two-step DFE inference framework employed here, θ_selected_, the expected level of polymorphism for each class of fitness-affecting polymorphisms, is treated as a fixed parameter of each joint DFE model. Assuming that mutation rates (μ) between neutral and selected classes are the same, θ_selected_ can be determined by multiplying θ_neutral_ obtained from the demographic model by the ratio of effective sequence lengths (L) between selected and neutral categories. To compute θ_selected_ for each genic mutation types, we multiply θ_neutral_ inferred from each population pair’s demographic model by the ratio of effective sequence lengths, L_selected_/L_neutral_. Effective sequence lengths used to compute θ_selected_ are tabulated in Supplementary Table S5 (see section *Classification of mutation types and functional elements* for details).

To obtain maximum likelihood estimates of each free parameter in tested joint DFE models, we used the BFGS optimization routine in ∂a∂i. Starting parameter values were permuted across 50,000 independent searches for each joint DFE model fit to each unfolded joint SFS. For each free parameter, we enforced the following upper and lower bounds during optimization: μ ∈ [-15, 10], σ ∈ [0, 10], *P*_neu_ ∈ [0, 1], *P*_pos_ ∈ [0, 1], ρ ∈ [0, 1], and *p_misid_* ∈ [0, 0.15]. After fitting each joint DFE model to each genic mutation type, we inspected parameter convergence and model likelihoods to select among contrasting models. Parameter convergence was evaluated based on the variance in estimated parameter values across the top 0.1% highest likelihood estimates for each model. Distributions of the top 0.1% highest likelihood parameter estimates for each model and mutation type are depicted in Supplementary Figures S3-S4 for island-mainland comparisons and Supplementary Figure S17 for island-island comparisons. Within each of the two joint DFE parameterization (bivariate lognormal with shared marginal parameters vs. bivariate lognormal with distinct marginal parameters), we selected among the three different DFE types (“deleterious-only” vs. “deleterious + neutral” vs. “deleterious + beneficial”) through either direct comparisons of model likelihoods (e.g., comparing “deleterious + neutral” vs. “deleterious + beneficial” DFE types with equivalent numbers of parameters), or through adjusted likelihood ratio tests (e.g., comparing nested “deleterious-only” DFEs to those with a discrete point mass of neutral or beneficial mutations) conducted using the Godambe Information Matrix (GIM) methods described in Coffman et al. (2016). Parameter estimates obtained under all tested joint DFE models are tabulated in Supplementary Table S7 for each mutation type and population comparison.

Due to either their increased complexity or poorer fit to the data, joint DFE models parameterized by a bivariate lognormal distribution with distinct marginal parameters exhibited poor convergence across model parameters in each island-mainland comparison (Supplementary Figure S3H-N; Supplementary Figure S4H-N). As a result, we excluded these parameterizations from downstream model comparisons and did not explore these more complex models for subsequent island-island joint DFEs. In addition, although we were able to robustly infer marginal DFE parameters (μ and σ) across mutation types under the bivariate lognormal distribution with shared marginal parameters, we found that the joint DFE correlation, ρ, could only be confidently estimated for nonsynonymous mutations (Supplementary Figure S3A; Supplementary Figure S4B). For all other mutation types, we observed poor convergence of this parameter across maximum-likelihood optimizations (Supplementary Figures S3B-G; Supplementary Figure S4A,C-G), which may suggest issues in the identifiability of the joint DFE correlation for certain variant classes. Supplementary Table S1 summarizes the results of adjusted likelihood ratio tests conducted within this simpler DFE parameterization, which we used to identify best-fit joint DFE models.

The resulting fit of these candidate models to the joint and marginal SFS are depicted in Supplementary Figures S5-S8 for island-mainland comparisons and Supplementary Figure S18 for island-island comparisons. To represent best-fitting joint DFEs (which specify the distribution of γ) in terms of *s*, we divide by 2*N_Anc_*, where *N_Anc_* represents the ancestral effective population size obtained from the corresponding demographic model (tabulated in Supplementary Table S6). Because of the extreme left-skew and long right tails of the lognormal distribution, we summarize the central tendencies of inferred marginal DFEs by their median value, compute as *e*^μ^. For best-fitting joint DFE models, we estimate standard deviations of each free parameter using the GIM methods formulated in Coffman et al. (2016) (tabulated in Supplementary Table S2). Marginal DFE error bars depicted in Figure 4, Supplementary Figure S9, and Supplementary Figure S12 reflect uncertainty (i.e., +/-1 standard deviation) in the inferred μ of each distribution.

## DATA AVAILABILITY

Whole genome sequences used in this study are available from the Sequence Read Archive (SRA) under BioProject accessions PRJNA1337437 and PRJNA1231036. See Supplementary Table S3 for associated BioSample IDs. All code used in this study is available on GitHub (https://github.com/PayseurLabUWMadison/island_mainland_dfe).

## ACKNOWLEDGEMENTS

We thank members of the Payseur lab for their helpful input on this work. We thank Hopi Hoekstra for the generous contribution of Gulf Islands *P. maniculatus* samples used in the study. Computational analyses were conducted using resources provided by the University of Wisconsin-Madison’s Center for High Throughput Computing.

## AUTHOR CONTRIBUTIONS

EKH and BAP designed the study. LEN and FB acquired the samples used for the study. EKH conducted the research with supervision from BAP. EKH and BAP wrote the manuscript with input from LEN and FB.

