## Supplementary Figure for "Evolution of mutational fitness effects in island populations"

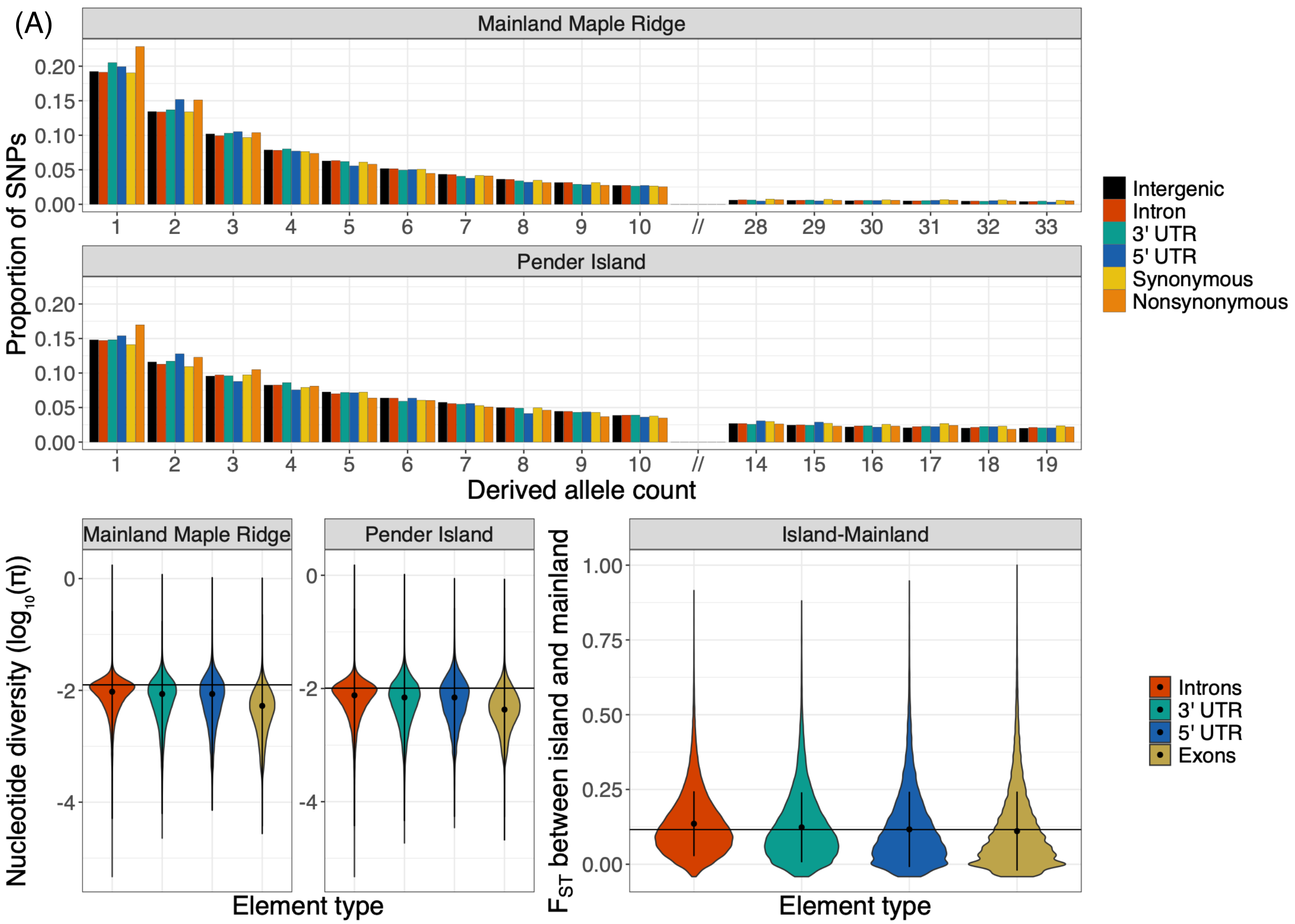


See figure legend below.

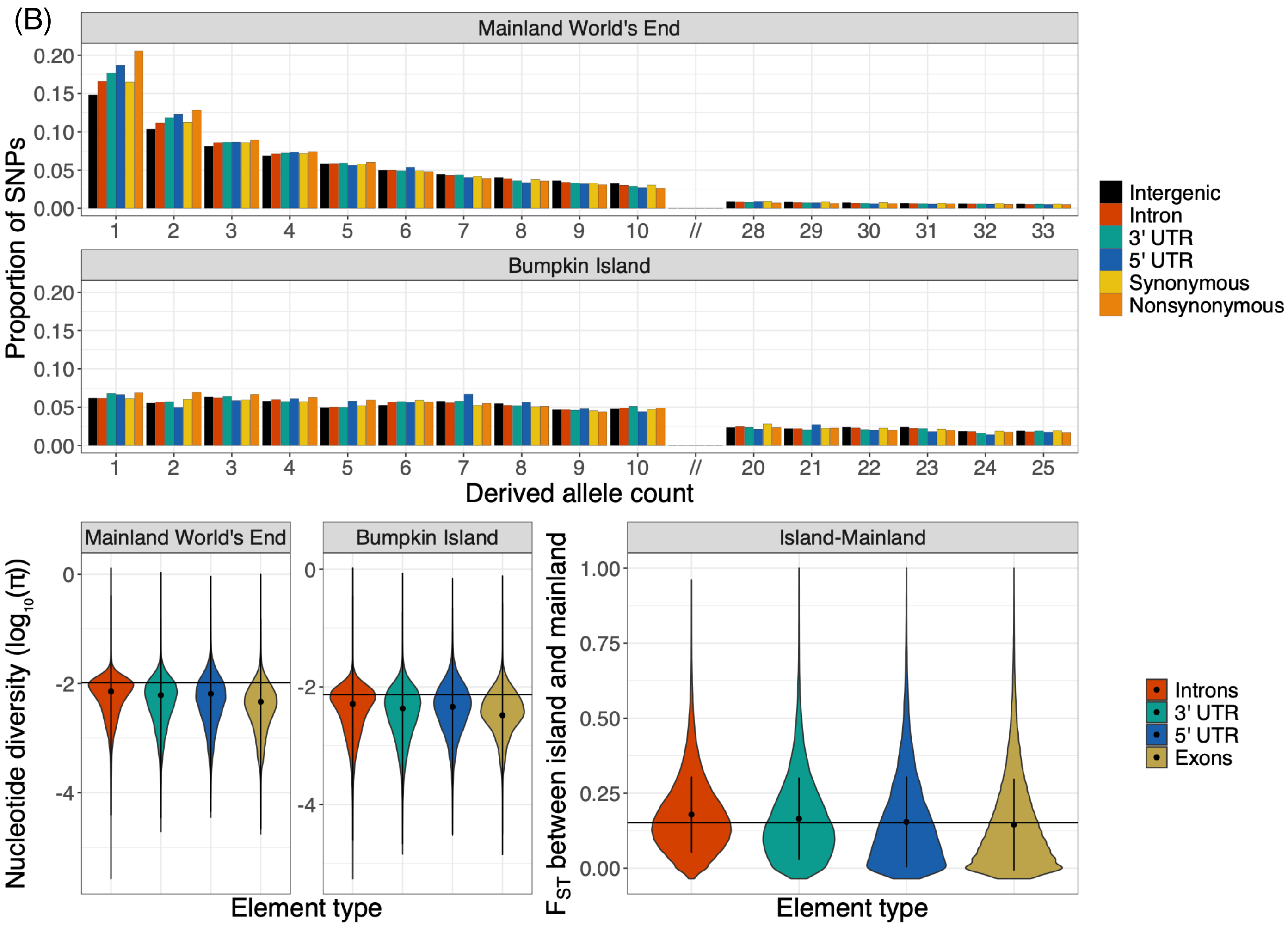


See figure legend below.

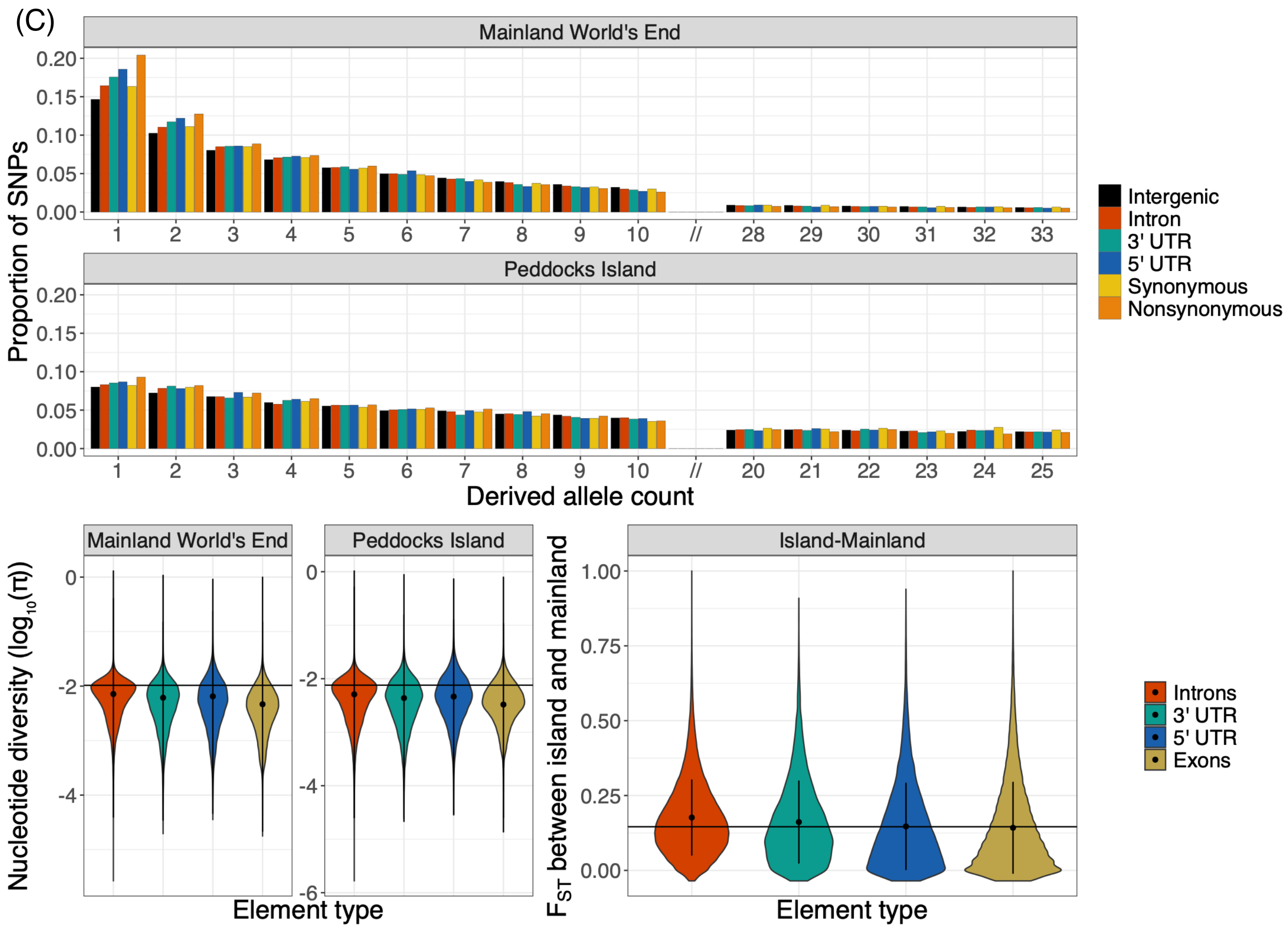


**Figure S1. Distinct signatures of purifying selection across genic elements.** Comparisons of the site frequency spectrum (SFS; top panel), nucleotide diversity (bottom left two panels), and island-mainland $F_{ST}$ (bottom right panel) across genic element/mutation types for the island-mainland contrasts excluded from Figure 2. Plots are arranged as in Figure 2 for the Pender-Maple Ridge **(A)**, Bumpkin-World's End **(B)**, and Peddocks-World's End **(C)** comparisons. See main text for details.

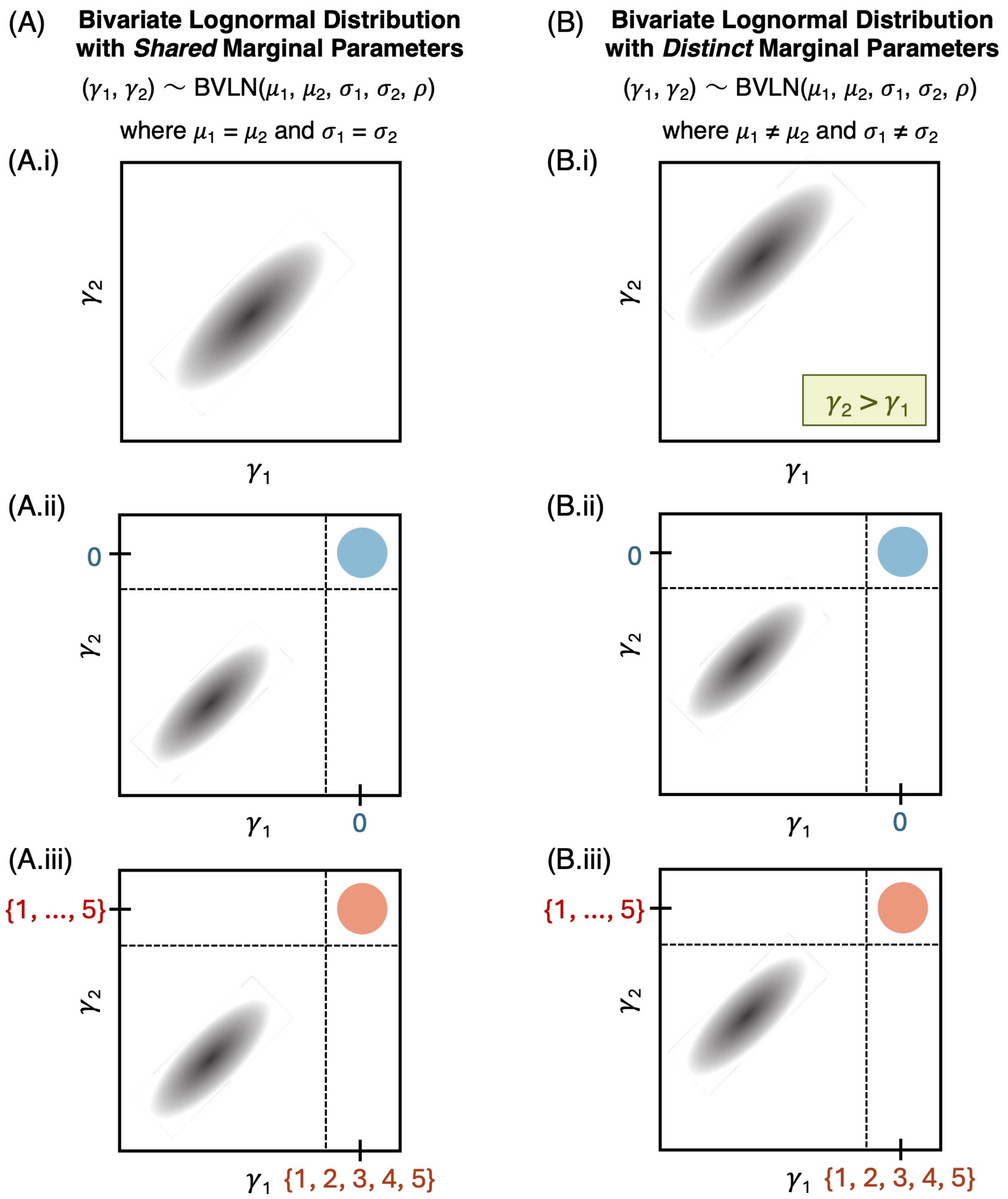


**Figure S2. Graphical representations of 2D probability distributions used to model the joint DFE.** We tested two different parameterizations of the joint DFE: **(A)** a bivariate lognormal distribution with *shared* marginal parameters ($\mu_1 = \mu_2$ and $\sigma_1 = \sigma_2$) and **(B)** a bivariate lognormal distribution with *distinct* marginal parameters ($\mu_1 \neq \mu_2$ and $\sigma_1 \neq \sigma_2$). On top of these basic parameterizations, we considered three different DFE types: **(i)** "deleterious-only" DFEs, where selection coefficients are continuously distributed according to a lognormal distribution with all $\gamma < 0$, **(ii)** "deleterious + neutral" DFEs, where a proportion $P_{neu}$ of mutations have a fixed $\gamma = 0$ and a proportion $1 - P_{neu}$ of mutations have a continuous, lognormal distribution of deleterious selection coefficients ($\gamma < 0$), and **(iii)** "deleterious + beneficial" DFEs, where a proportion $P_{pos}$ of mutations have a fixed $\gamma \in [1, 2, 3, 4, 5]$ and a proportion $1 - P_{pos}$ of mutations have a continuous, lognormal distribution of deleterious selection coefficients ($\gamma < 0$). For types **(ii)** and **(iii)**, we assume that the discrete point mass ($P_{neu}$ or $P_{pos}$) is symmetric between populations. In addition to estimating the values of these shared (or distinct) $\mu$ and $\sigma$, all joint DFE models include a parameter, $\rho$, which represents the correlation in mutational fitness effects between populations, and a $p_{misid}$ parameter which represents the rate of ancestral state misidentification. Models of type **(ii)** and **(iii)** include additional parameters for $P_{neu}$ and $P_{pos}$, respectively.

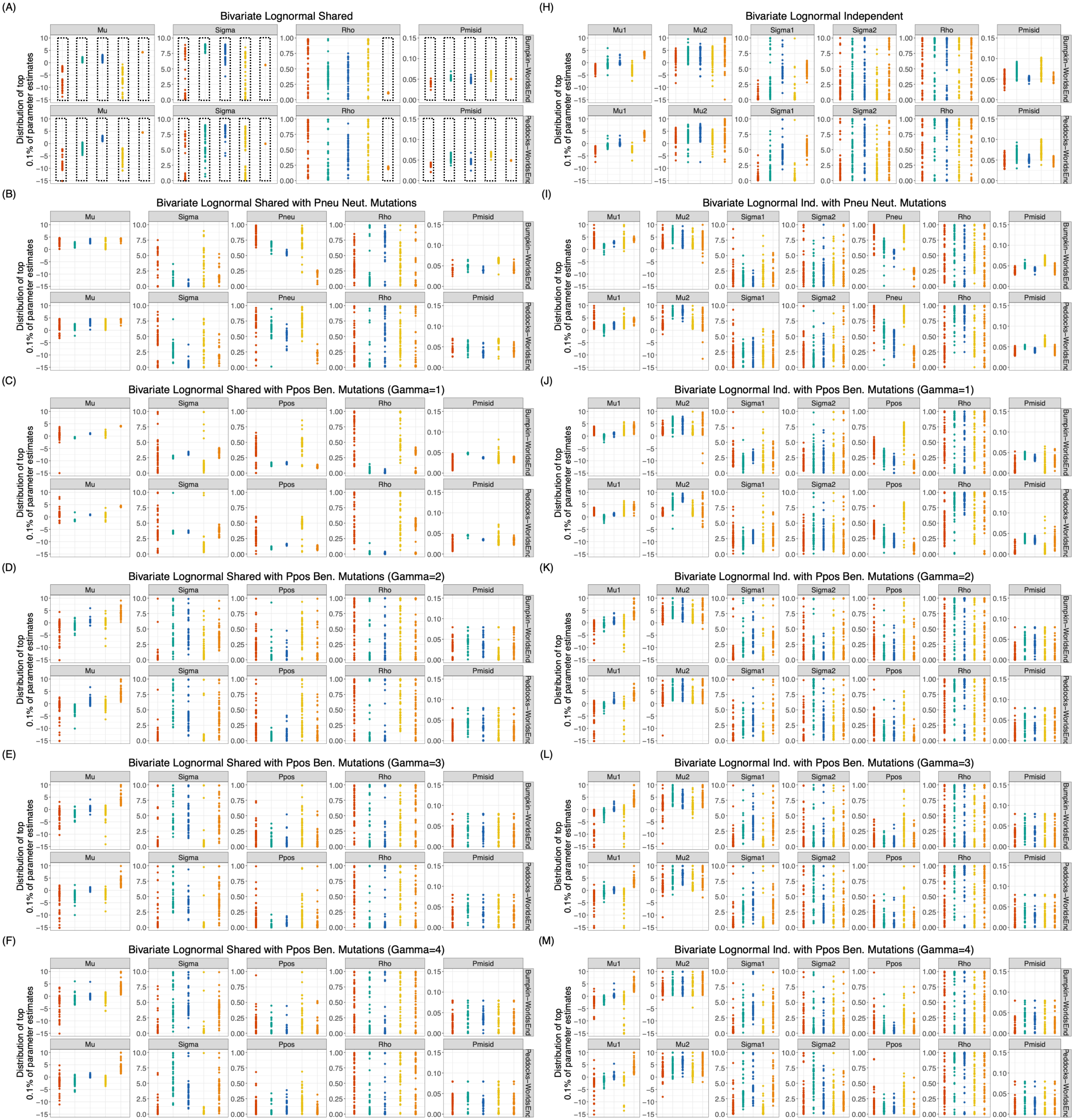
(A)
Bivariate Lognormal Shared
(B)
Bivariate Lognormal Shared with Pneu Neut. Mutations
(C)
Bivariate Lognormal Shared with Ppos Ben. Mutations (Gamma=1)
(D)
Bivariate Lognormal Shared with Ppos Ben. Mutations (Gamma=2)
(E)
Bivariate Lognormal Shared with Ppos Ben. Mutations (Gamma=3)
(F)
Bivariate Lognormal Shared with Ppos Ben. Mutations (Gamma=4)
(H)
Bivariate Lognormal Independent
(I)
Bivariate Lognormal Ind. with Pneu Neut. Mutations
(J)
Bivariate Lognormal Ind. with Ppos Ben. Mutations (Gamma=1)
(K)
Bivariate Lognormal Ind. with Ppos Ben. Mutations (Gamma=2)
(L)
Bivariate Lognormal Ind. with Ppos Ben. Mutations (Gamma=3)
(M)
Bivariate Lognormal Ind. with Ppos Ben. Mutations (Gamma=4)
Mu
Sigma
Pneu
Ppos
Rho
Pmisid
Mu1
Mu2
Sigma1
Sigma2
Bumpkin–WorldsEnd
Peddocks–WorldsEnd
Distribution of top
0.1% of parameter estimates

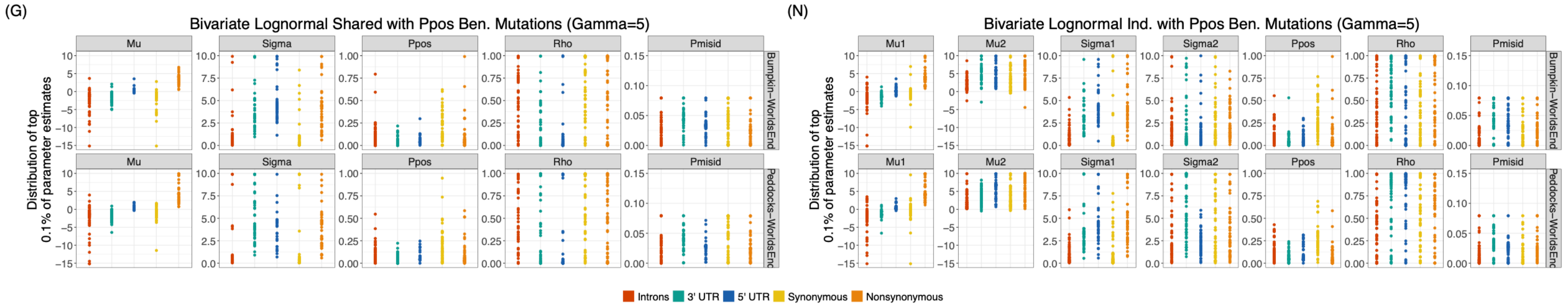


**Figure S3. Parameter convergence across tested joint DFE models for Boston Harbor *P. leucopus*.** Each panel depicts parameter convergence for a specific joint DFE model, indicated by panel title. Panels **A-G** on the left reflect joint DFEs modeled by a bivariate lognormal distribution with shared marginal parameters, μ and σ. Panels **H-N** on the right reflect joint DFEs modeled by a bivariate lognormal distribution with *distinct* marginal parameters ($\mu_1 \neq \mu_2$ and $\sigma_1 \neq \sigma_2$). The first row (panels **A** and **N**) represents "deleterious-only" DFEs, where selection coefficients are continuously distributed according to a lognormal distribution with all $\gamma < 0$. The second row (panels **B** and **I**) represents "deleterious + neutral" DFEs, where a proportion $P_{neu}$ of mutations have a fixed $\gamma = 0$ and a proportion $1 - P_{neu}$ of mutations have a continuous, lognormal distribution of deleterious selection coefficients ($\gamma < 0$). Remaining rows (panels **C-G** and **J-N**) represent "deleterious + beneficial" DFEs, where a proportion $P_{pos}$ of mutations have a fixed $\gamma \in [1, 2, 3, 4, 5]$ and a proportion $1 - P_{pos}$ of mutations have a continuous, lognormal distribution of deleterious selection coefficients ($\gamma < 0$). Within each panel, convergence results are stratified by model parameter (column names) and island-mainland comparison (row names). Y-axes span the upper and lower bounds enforced for each parameter value. X-axis groupings reflect different mutation types. Points indicate the distribution of parameter estimates observed across the top 0.1% highest likelihood parameter combinations obtained from ∂a∂i's BFGS optimization routine for a given mutation type. Dotted boxes in panel **A** illustrate the degree of convergence of each parameter in the best-fit models presented in the main text. Because the joint DFE correlation, ρ, exhibited poor convergence for all but nonsynonymous mutations, we exclude discussion of this parameter for intronic, synonymous, 3' UTR, and 5' UTR mutations.

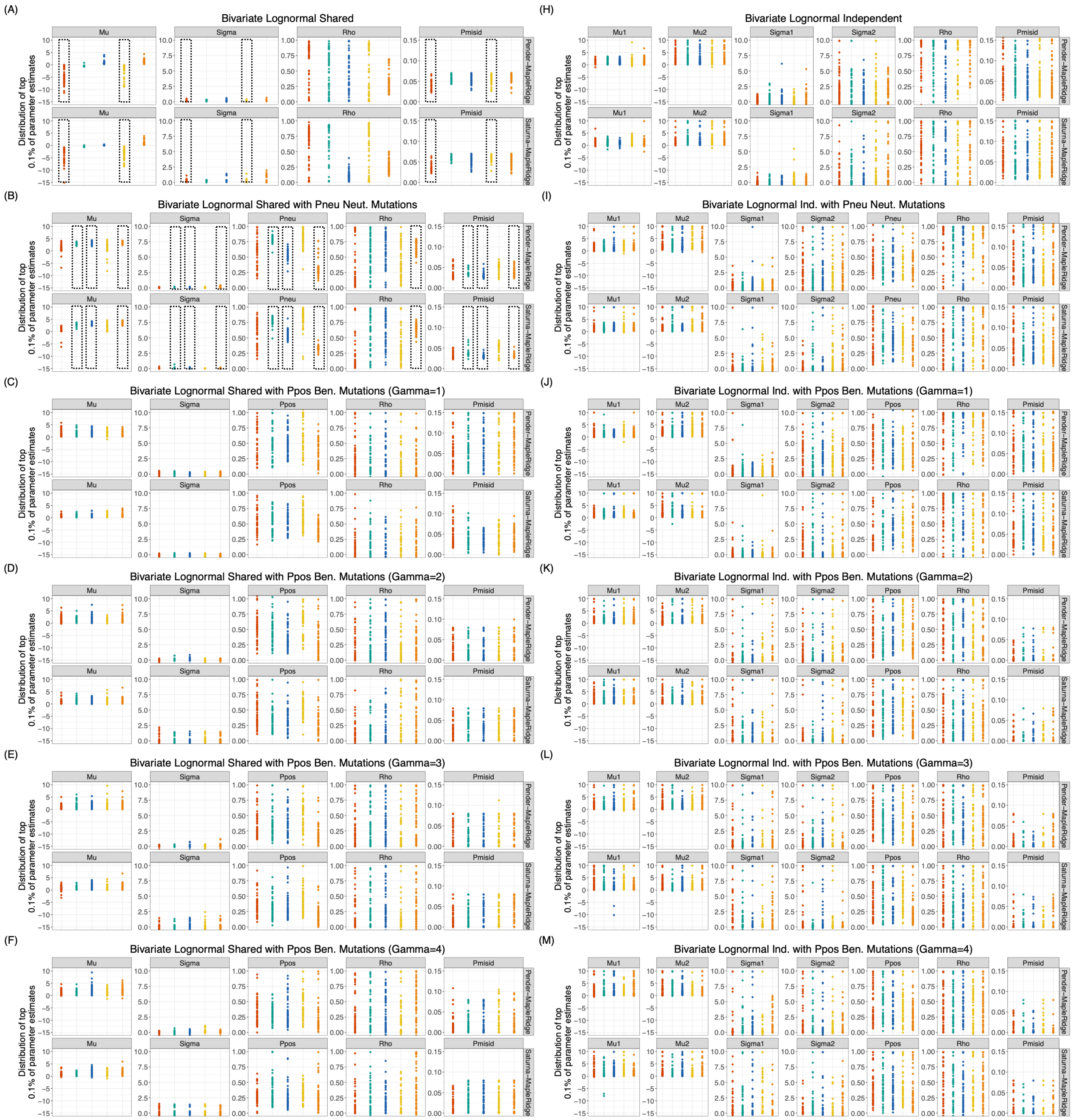
(A)
Bivariate Lognormal Shared
Mu
Sigma
Rho
Pmisid
Pender−MapleRidge
Saturna−MapleRidge
Distribution of top 0.1% of parameter estimates
(H)
Bivariate Lognormal Independent
Mu1
Mu2
Sigma1
Sigma2
(B)
Bivariate Lognormal Shared with Pneu Neut. Mutations
Pneu
(I)
Bivariate Lognormal Ind. with Pneu Neut. Mutations
(C)
Bivariate Lognormal Shared with Ppos Ben. Mutations (Gamma=1)
Ppos
(J)
Bivariate Lognormal Ind. with Ppos Ben. Mutations (Gamma=1)
(D)
Bivariate Lognormal Shared with Ppos Ben. Mutations (Gamma=2)
(K)
Bivariate Lognormal Ind. with Ppos Ben. Mutations (Gamma=2)
(E)
Bivariate Lognormal Shared with Ppos Ben. Mutations (Gamma=3)
(L)
Bivariate Lognormal Ind. with Ppos Ben. Mutations (Gamma=3)
(F)
Bivariate Lognormal Shared with Ppos Ben. Mutations (Gamma=4)
(M)
Bivariate Lognormal Ind. with Ppos Ben. Mutations (Gamma=4)

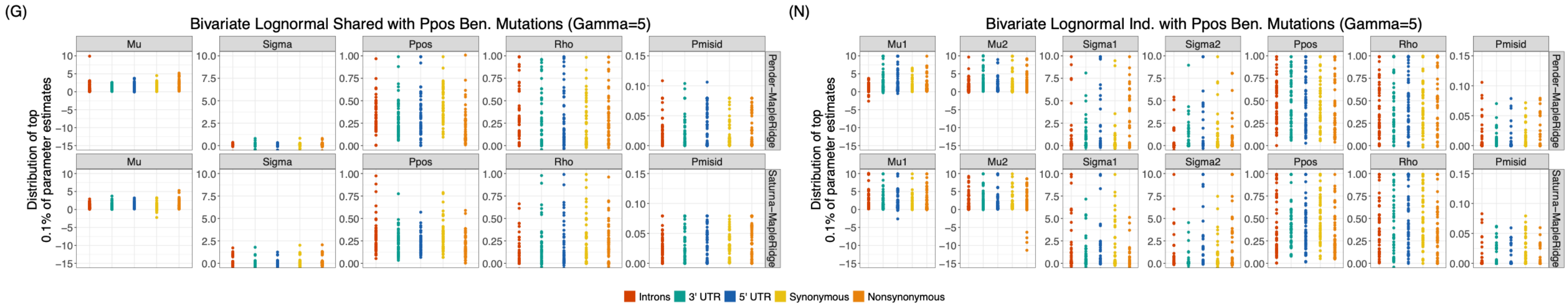


**Figure S4. Parameter convergence across tested joint DFE models for Gulf Islands *P. maniculatus*.** Panels are arranged as in Supplementary Figure S3. Dotted boxes in panels **A** and **B** illustrate the degree of convergence of each parameter in the best-fit models presented in the main text. Because the joint DFE correlation, ρ, exhibited poor convergence for all but nonsynonymous mutations, we exclude discussion of this parameter for intronic, synonymous, 3’ UTR, and 5’ UTR mutations. See Supplementary Figure S3 legend for additional details.

(A) Peddocks-World's End residual fit; synonymous mutations

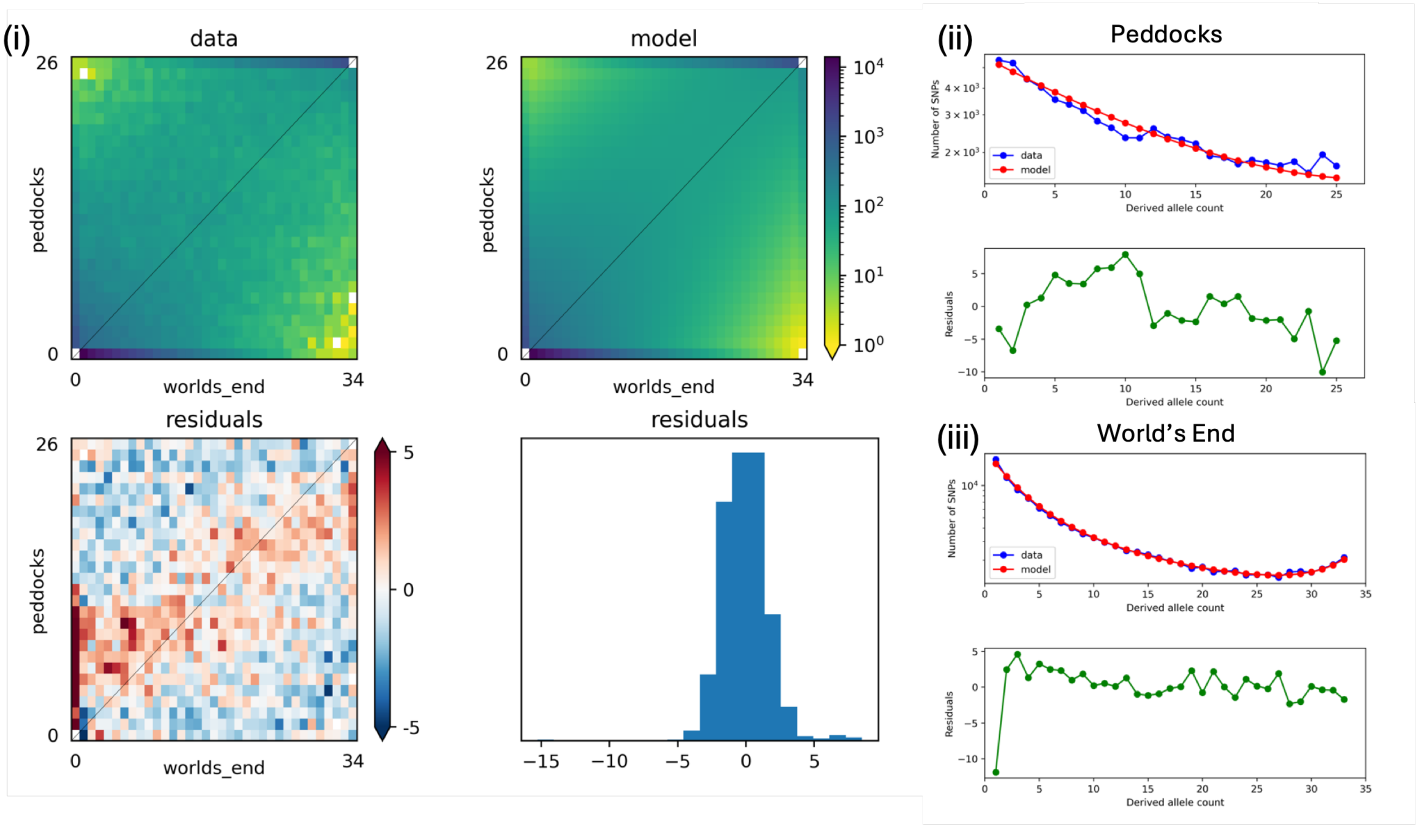


(B) Peddocks-World's End residual fit; nonsynonymous mutations

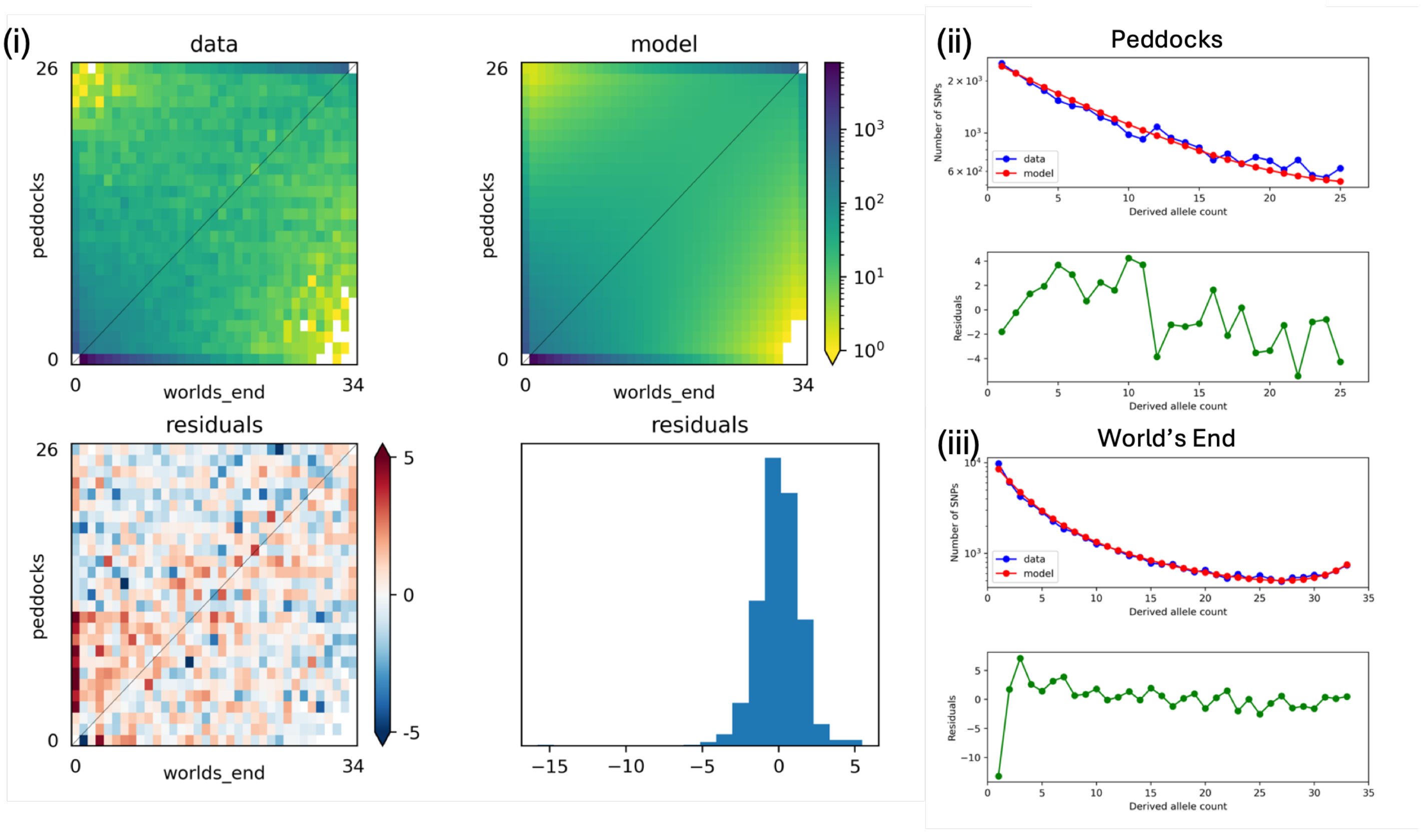

### (C) Peddocks-World's End residual fit; 3' UTR mutations

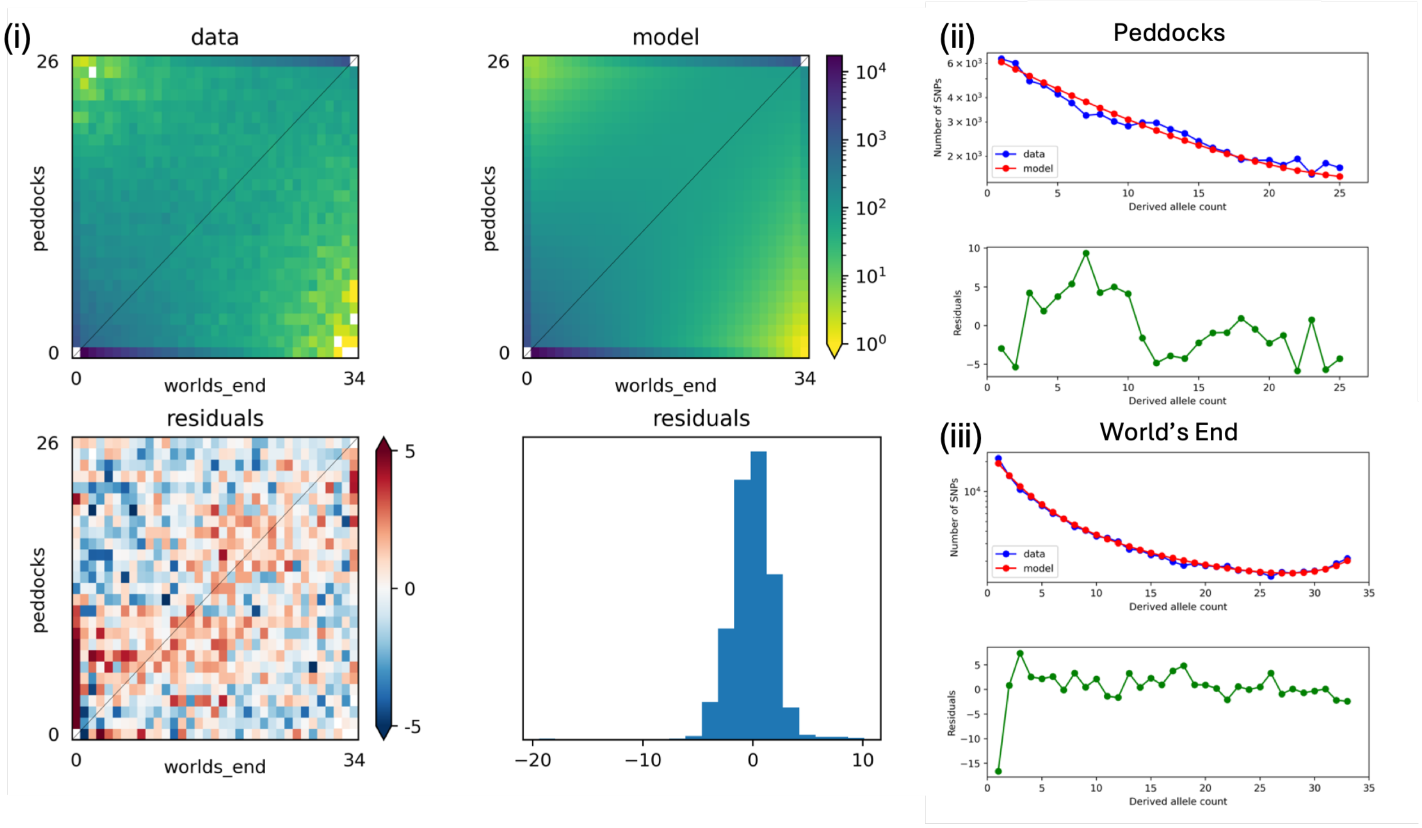


### (D) Peddocks-World's End residual fit; 5' UTR mutations

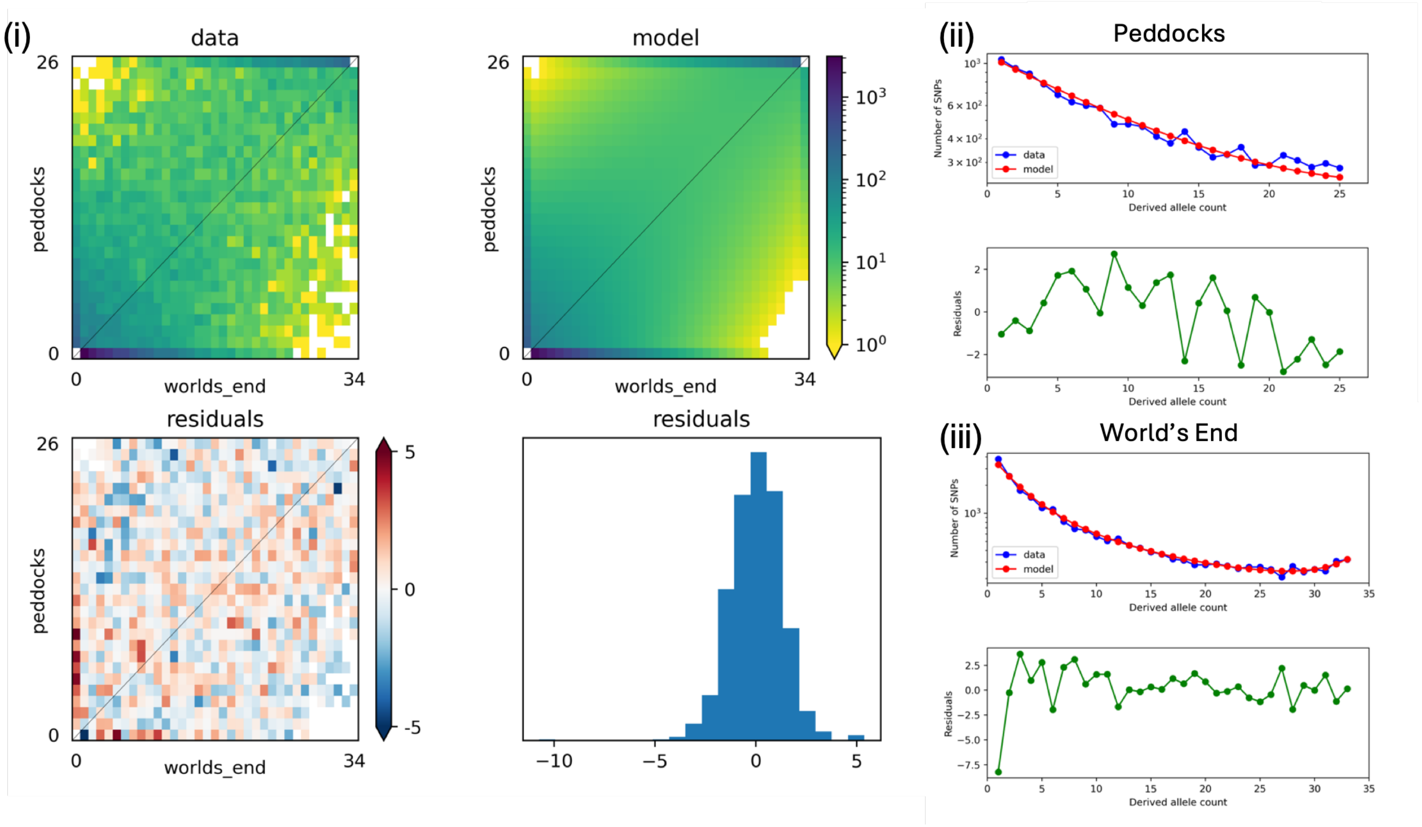

(E) Peddocks-World's End residual fit; intron mutations

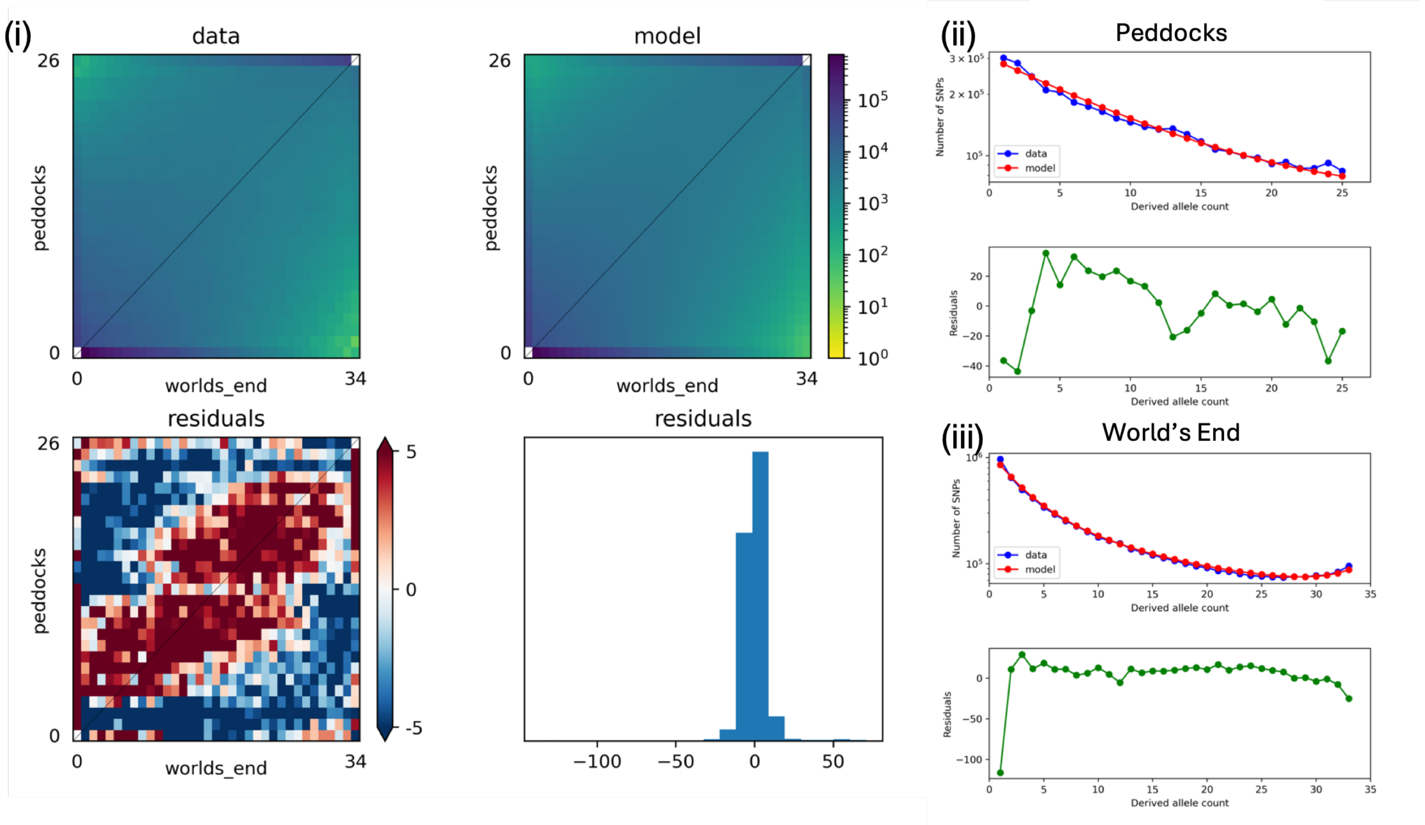


**Figure S5. Candidate joint DFE model fits to genic mutations segregating among Peddocks Island and mainland World's End *P. leucopus*.** Each panel depicts the fit of the inferred joint DFE model to observed allele frequencies for each mutation type: synonymous mutations **(A)**, nonsynonymous mutations **(B)**, 3' UTR mutations **(C)**, 5' UTR mutations **(D)**, and intronic mutations **(E)**. Within each panel, **(i)** depicts differences between the joint site frequency spectrum (jSFS) predicted by the model (top right) and the observed jSFS (top left). Each entry (*x*, *y*) in the jSFS represents the proportion of SNPs observed in *x* copies in the mainland World's End sample (x-axis) and *y* copies in the Peddocks Island sample (y-axis). The bottom left plot measures differences between these jSFS. Here, the color of each cell represents the Anscombe residual difference, where cooler colors indicate that the model underpredicts the observed number of SNPs and warmer colors indicate an overprediction by the model. The accompanying histogram (bottom right) plots the distribution of these residual values. **(ii-iii)** plot comparisons between model and data for the marginal SFS in each population. Top plots depict the expected (red) and observed (blue) SFS. Bottom plots depict the residual value for each SFS bin in green.

(A) Bumpkin-World's End residual fit; synonymous mutations

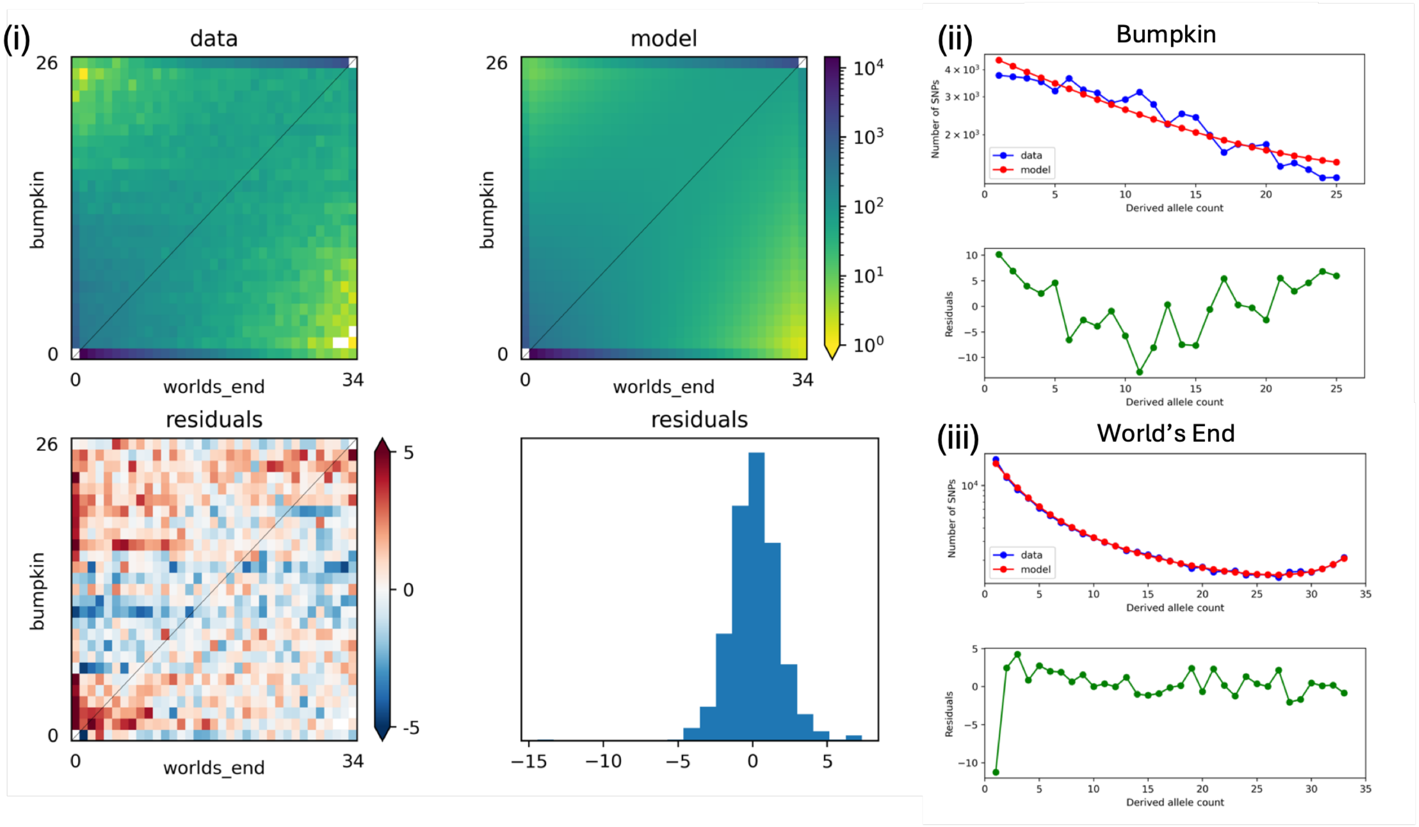


(B) Bumpkin-World's End residual fit; nonsynonymous mutations

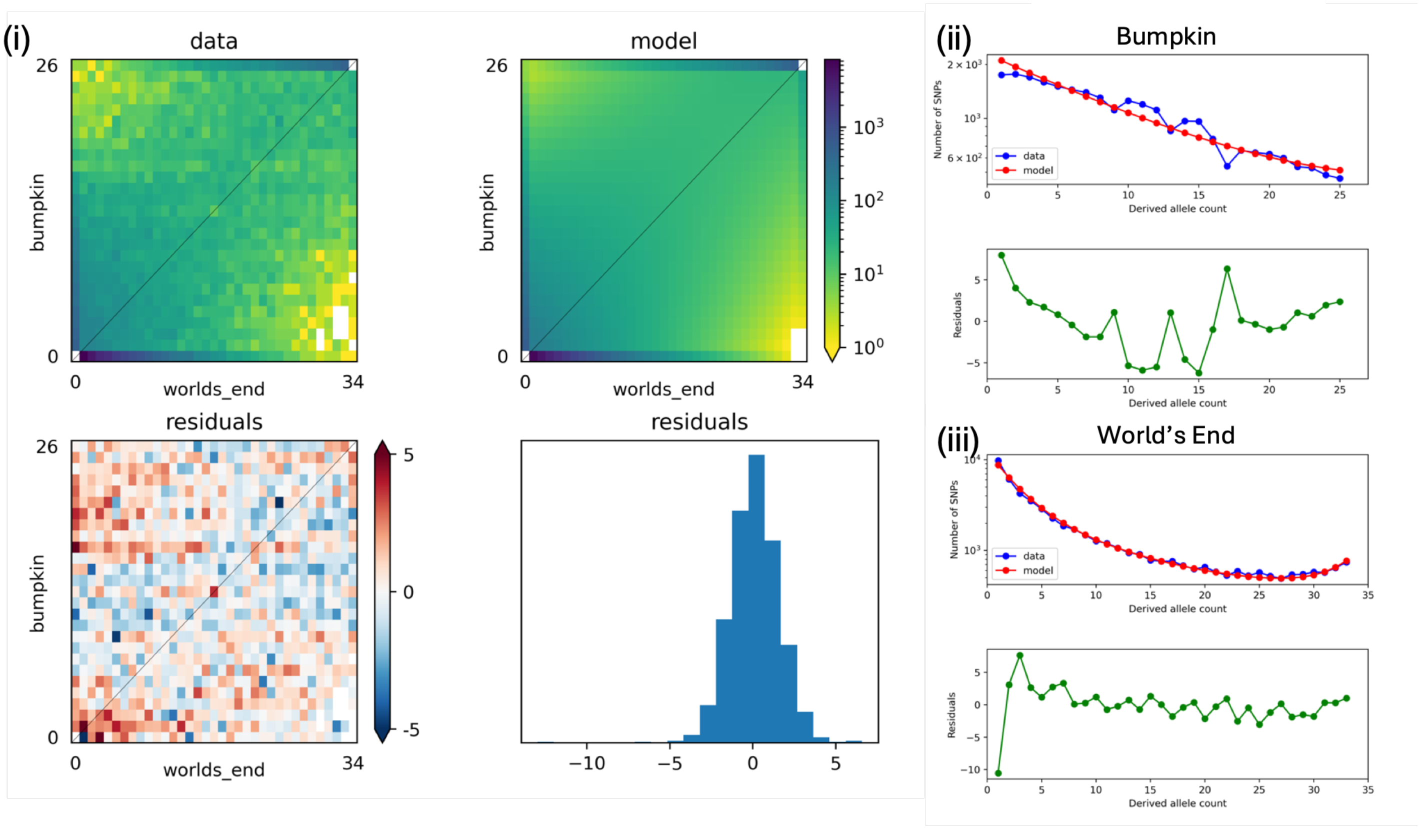

### (C) Bumpkin-World's End residual fit; 3' UTR mutations

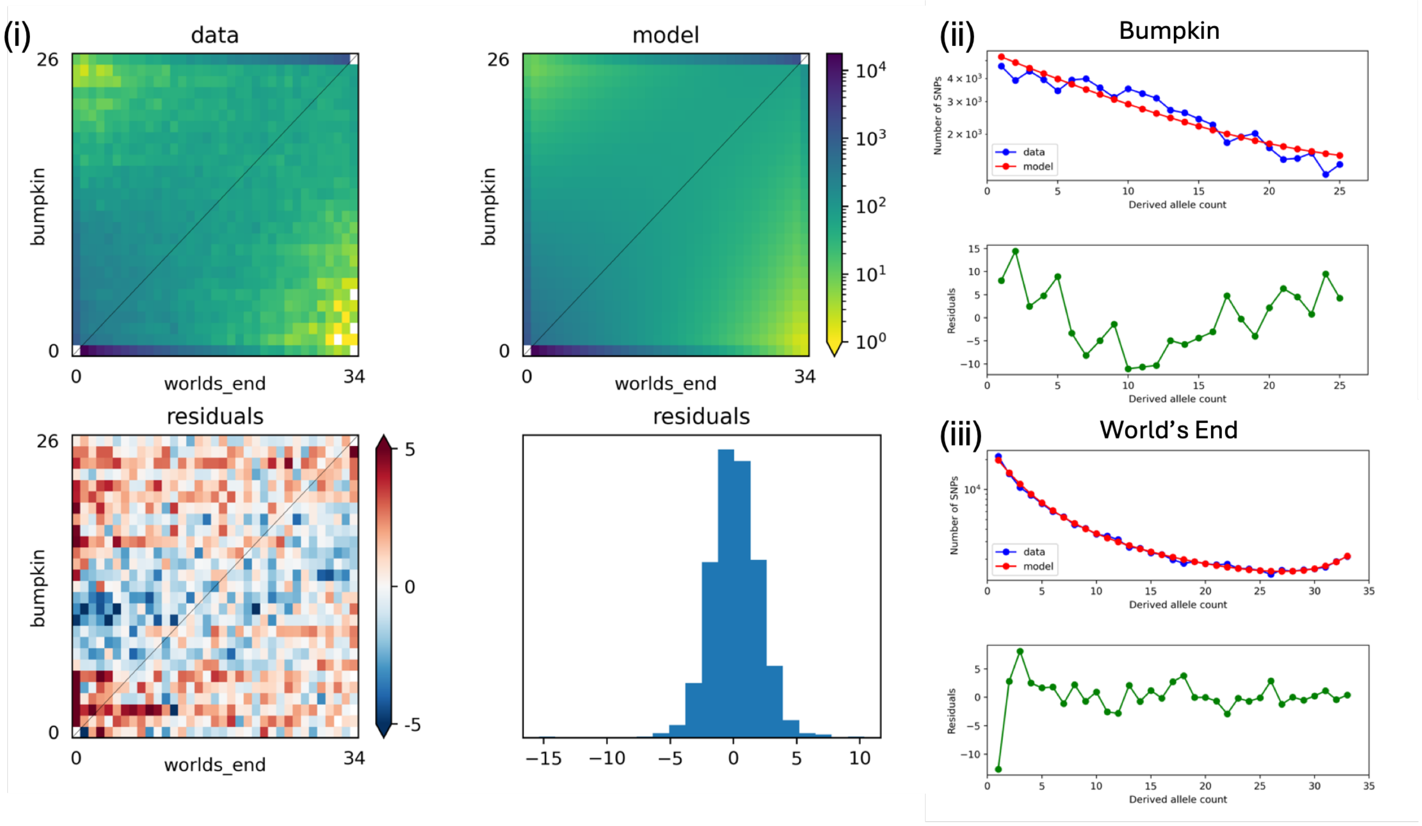


### (D) Bumpkin-World's End residual fit; 5' UTR mutations

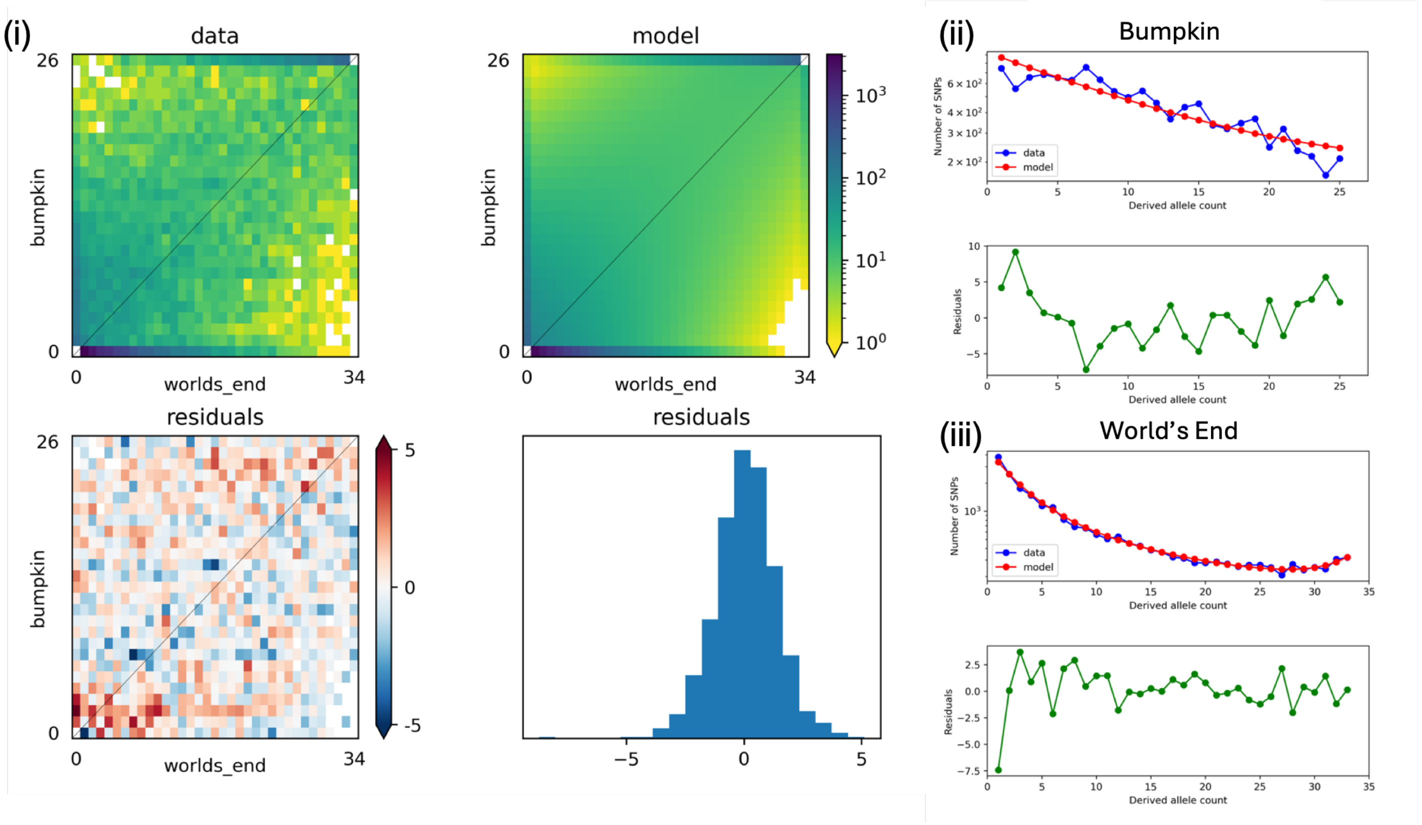

(E) Bumpkin-World's End residual fit; intron mutations

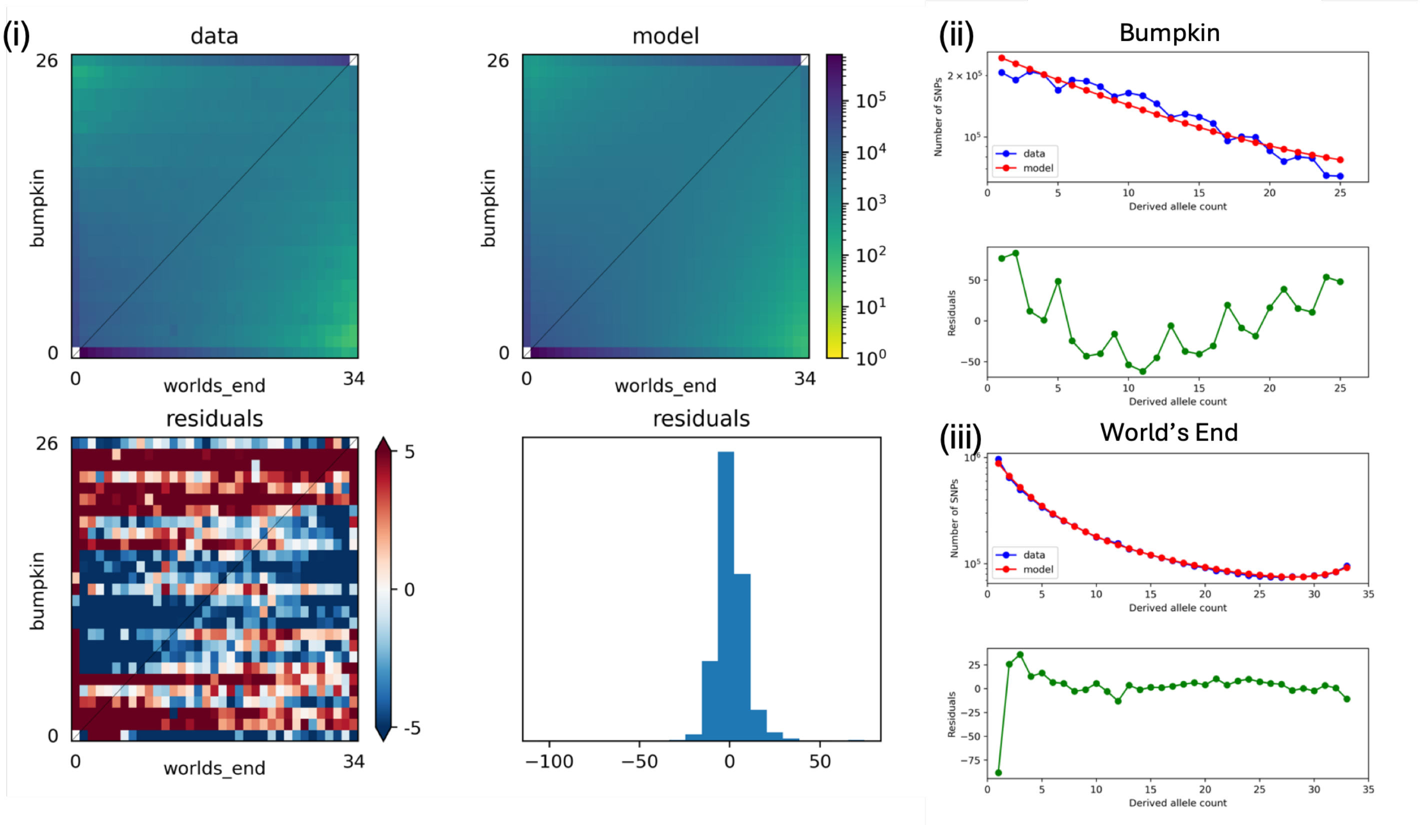


**Figure S6. Candidate joint DFE model fits to genic mutations segregating among Bumpkin Island and mainland World's End *P. leucopus*.** Plots are arranged as in Supplementary Figure S5 for the Bumpkin-World's End comparison. See Supplementary Figure S5 legend for details.

### (A) Pender-Maple Ridge residual fit; synonymous mutations

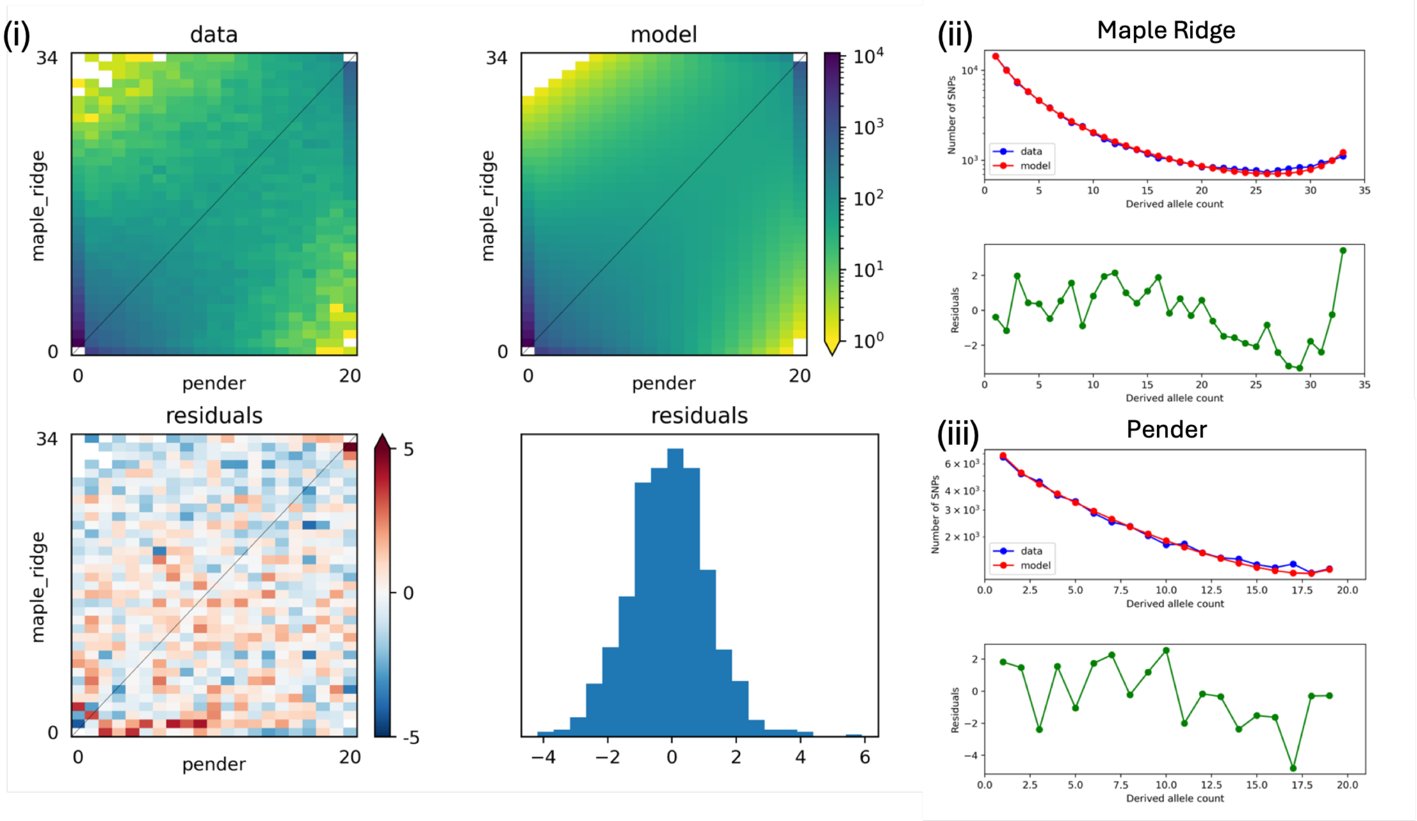


### (B) Pender-Maple Ridge residual fit; nonsynonymous mutations

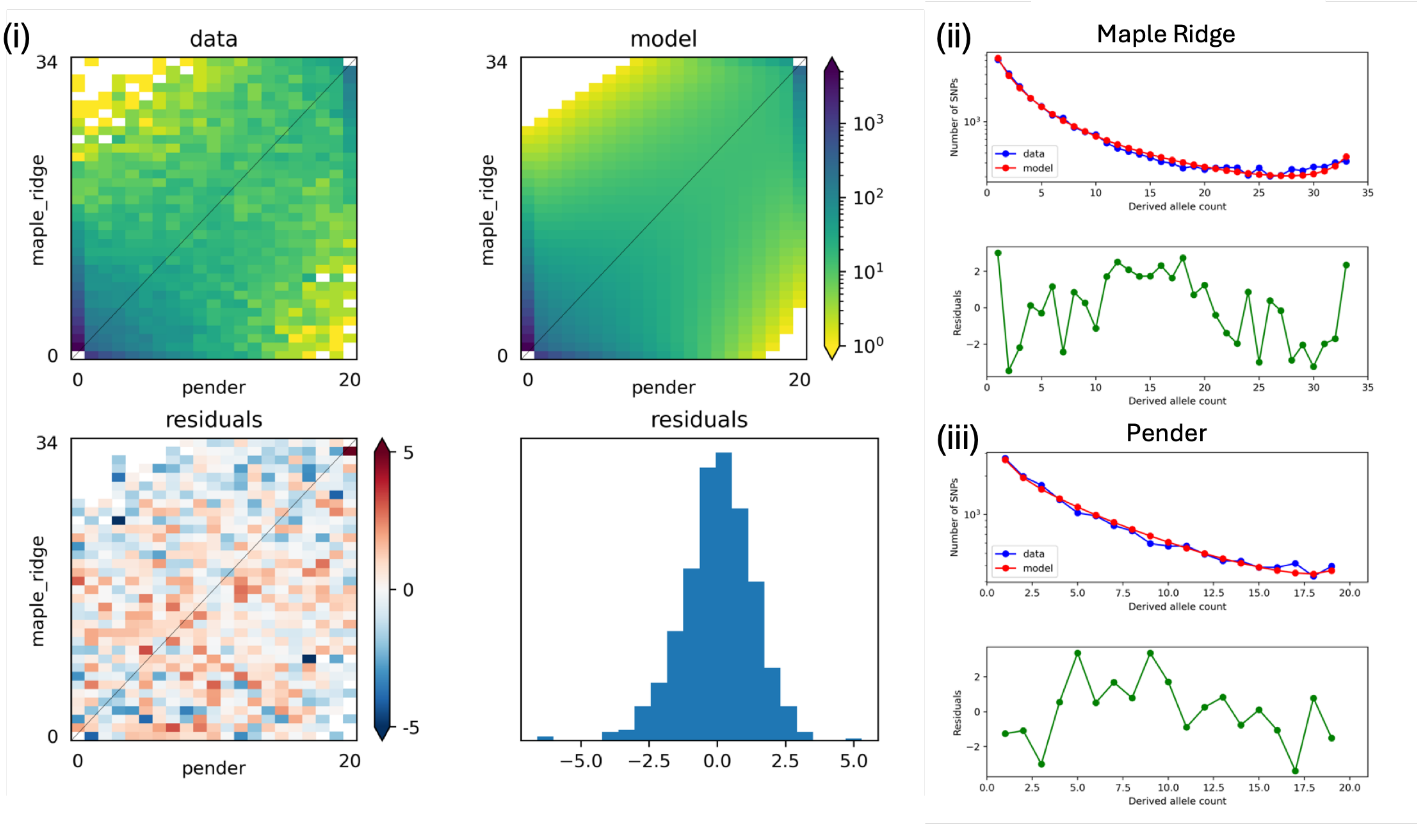

### (C) Pender-Maple Ridge residual fit; 3' UTR mutations

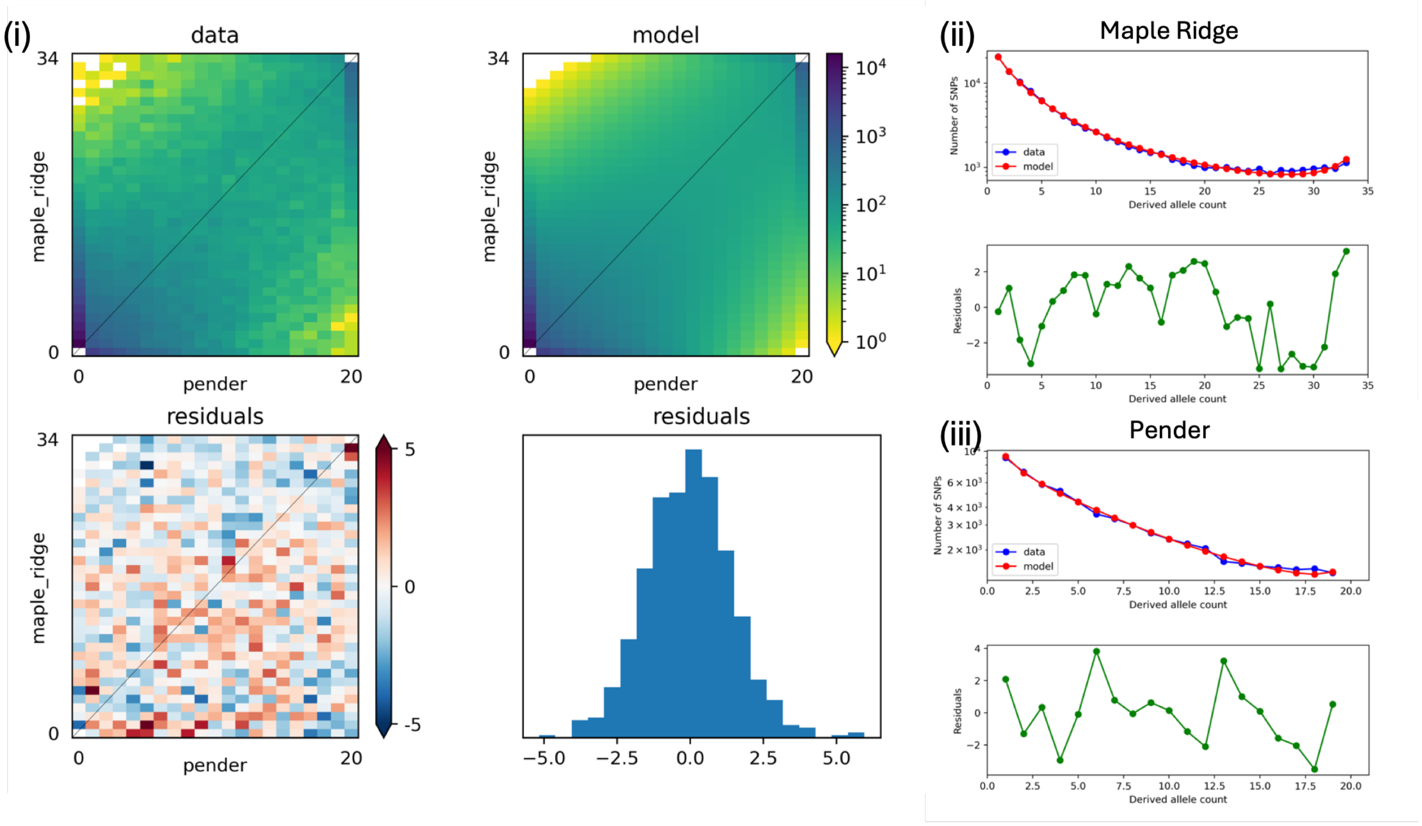


### (D) Pender-Maple Ridge residual fit; 5' UTR mutations

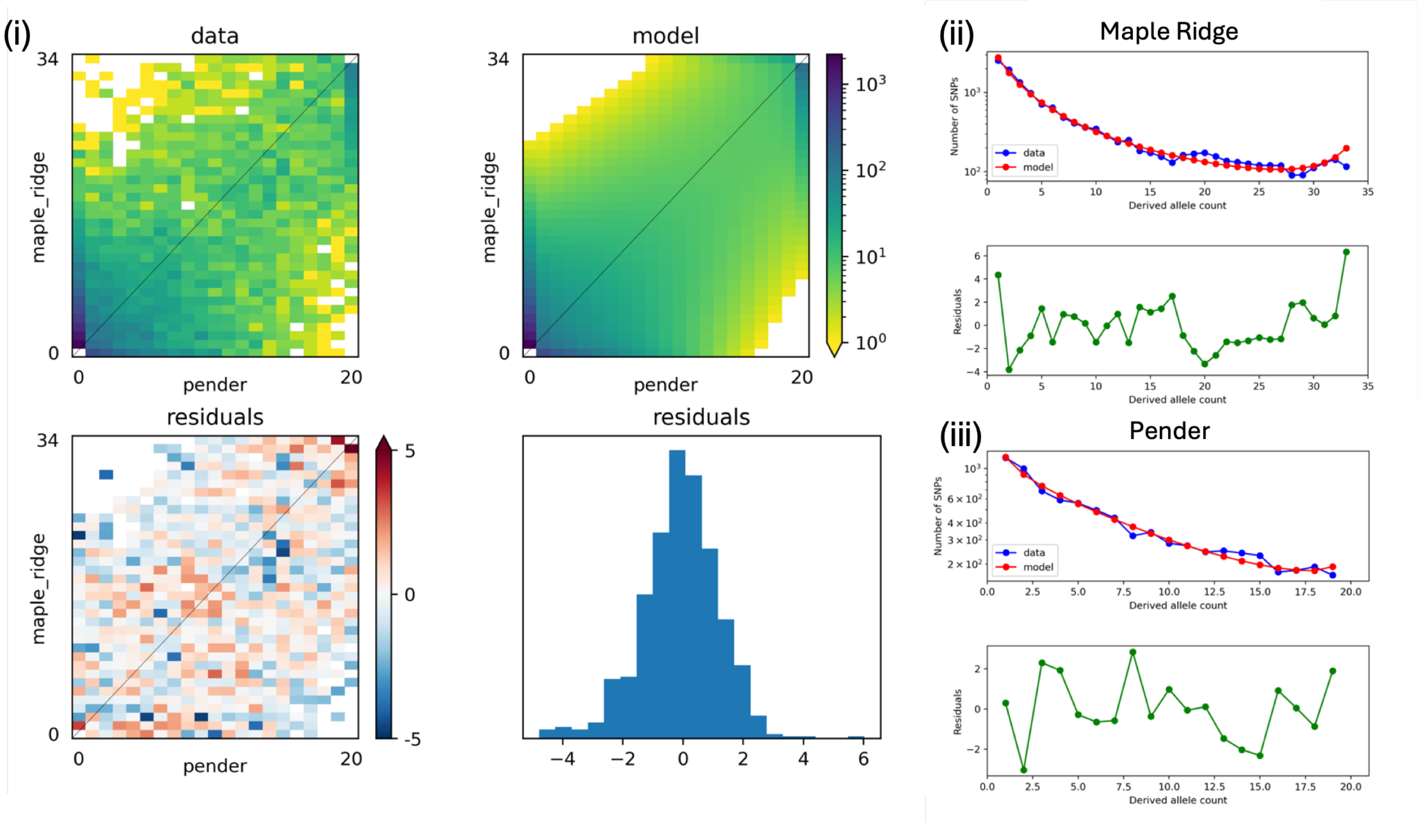

(E) Pender-Maple Ridge residual fit; intron mutations

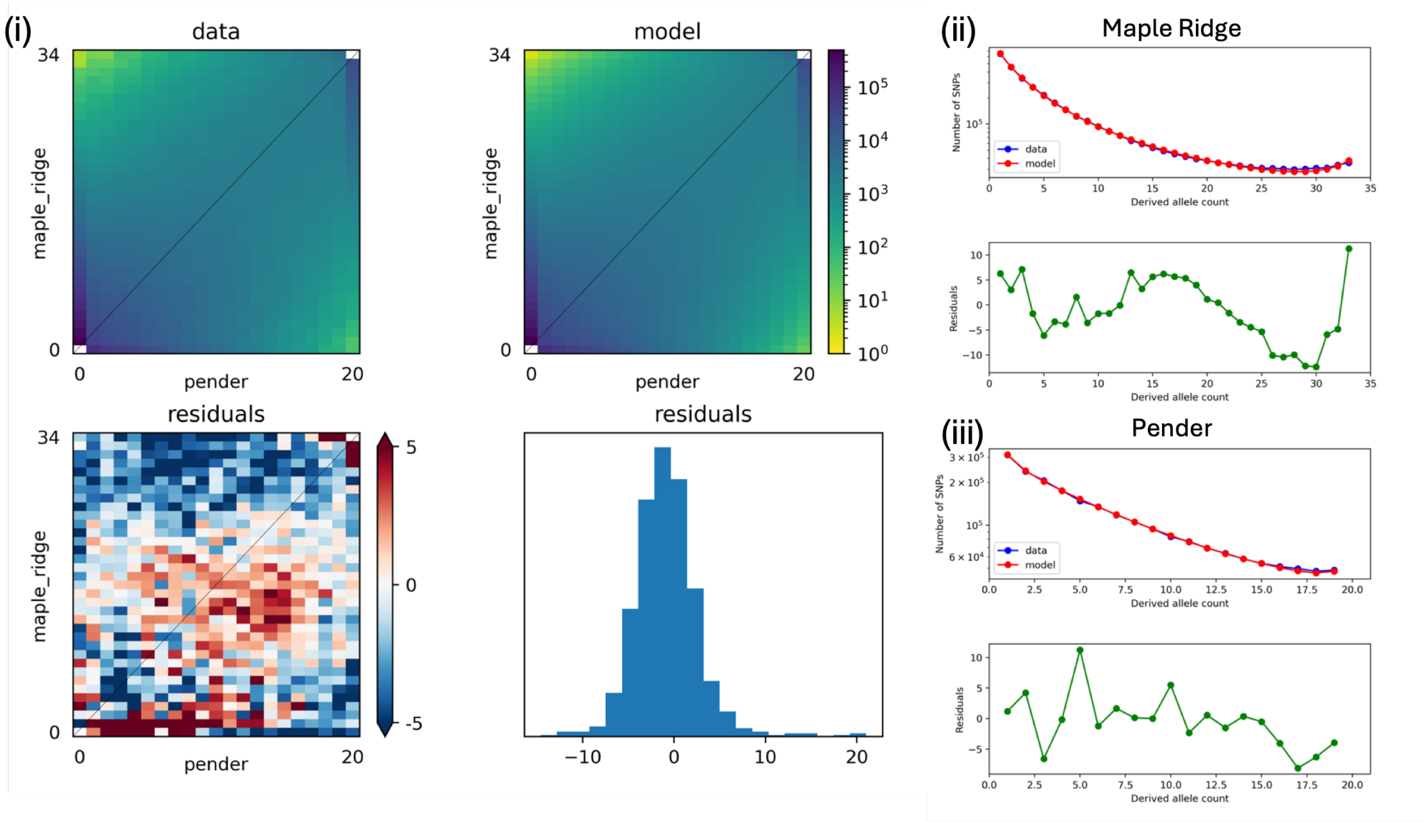


**Figure S7. Candidate joint DFE model fits to genic mutations segregating among Pender Island and mainland Maple Ridge *P. maniculatus*.** Plots are arranged as in Supplementary Figure S5 for the Pender-Maple Ridge comparison. See Supplementary Figure S5 legend for details.

(A) Saturna-Maple Ridge residual fit; synonymous mutations

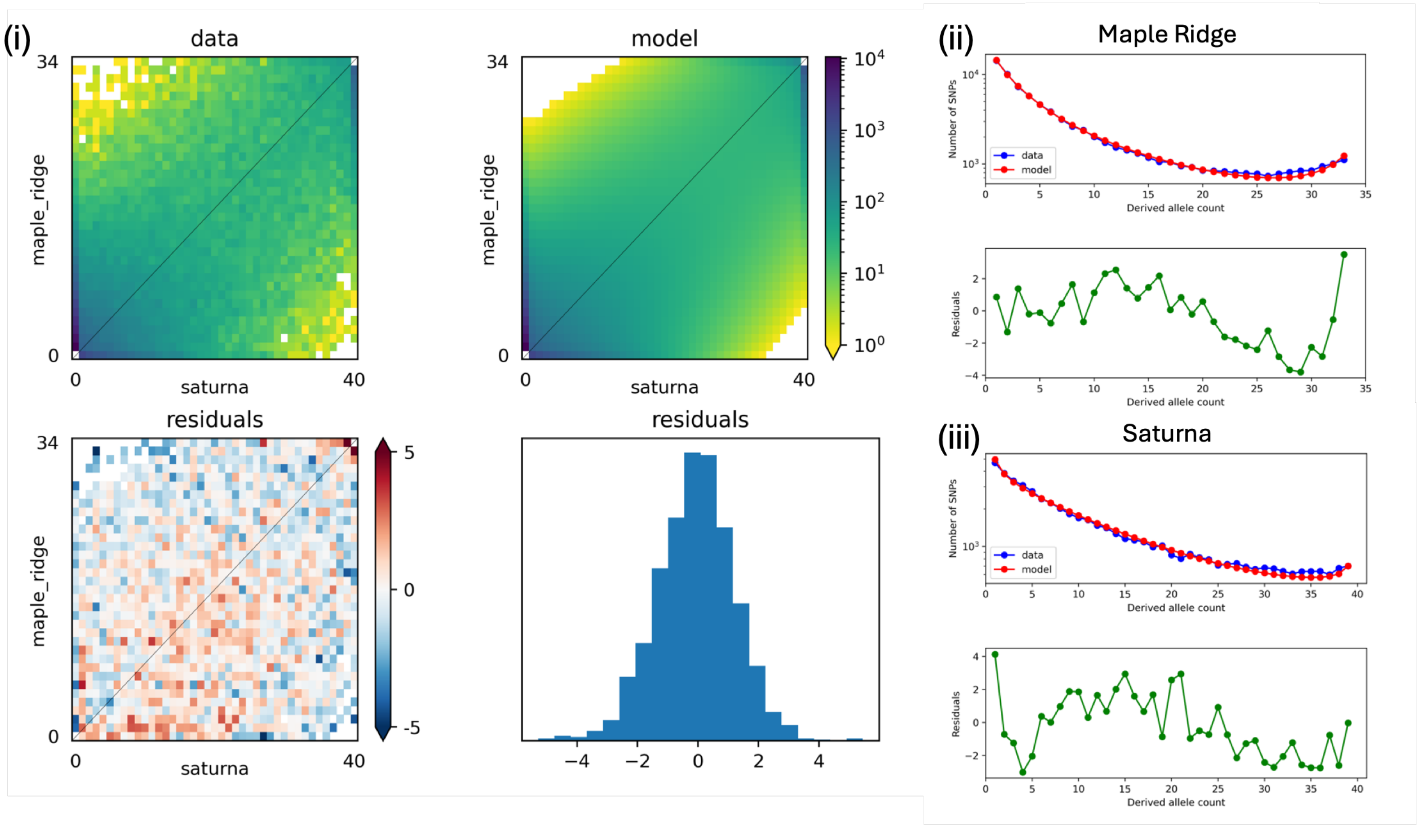


(B) Saturna-Maple Ridge residual fit; nonsynonymous mutations

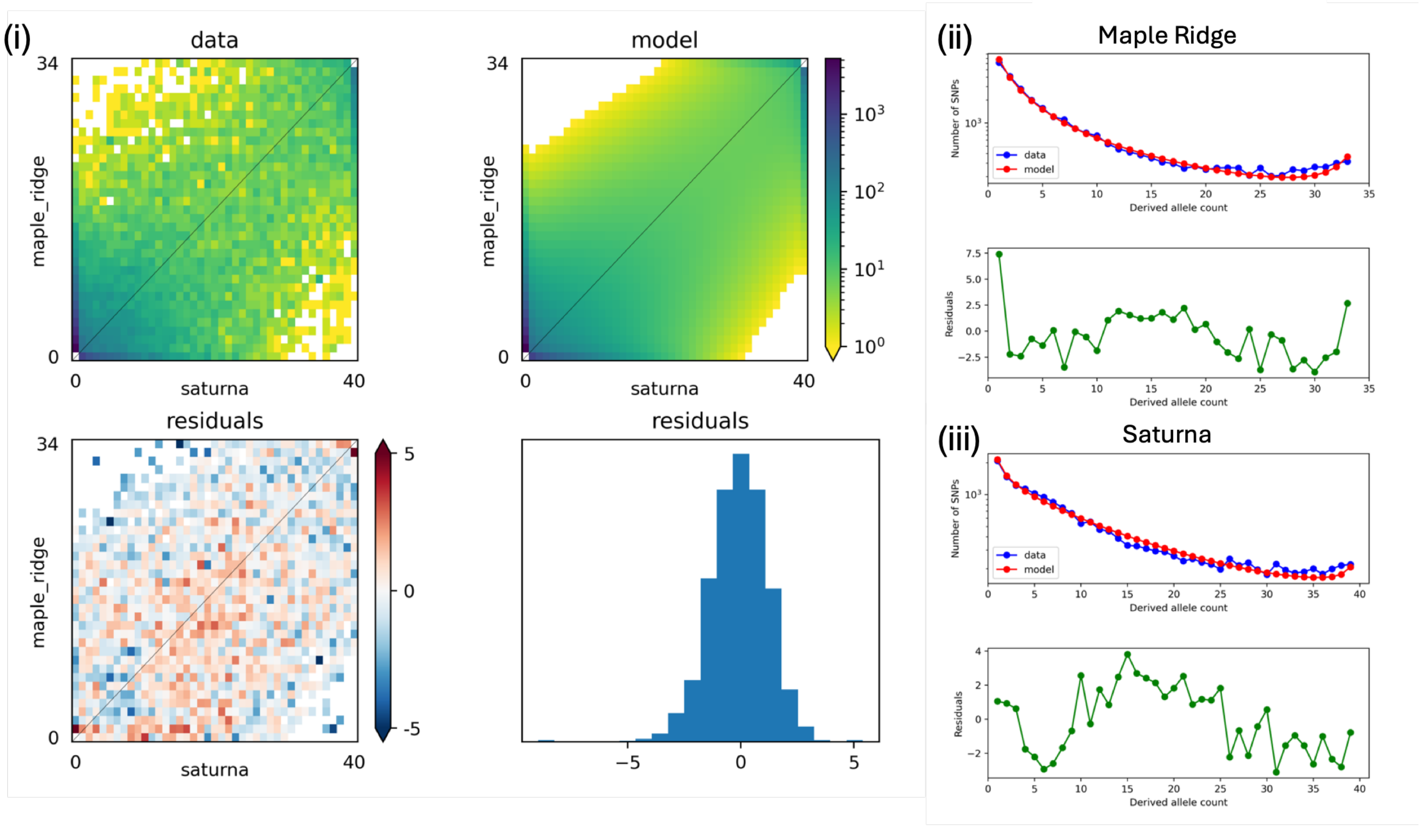

(C) Saturna-Maple Ridge residual fit; 3’ UTR mutations

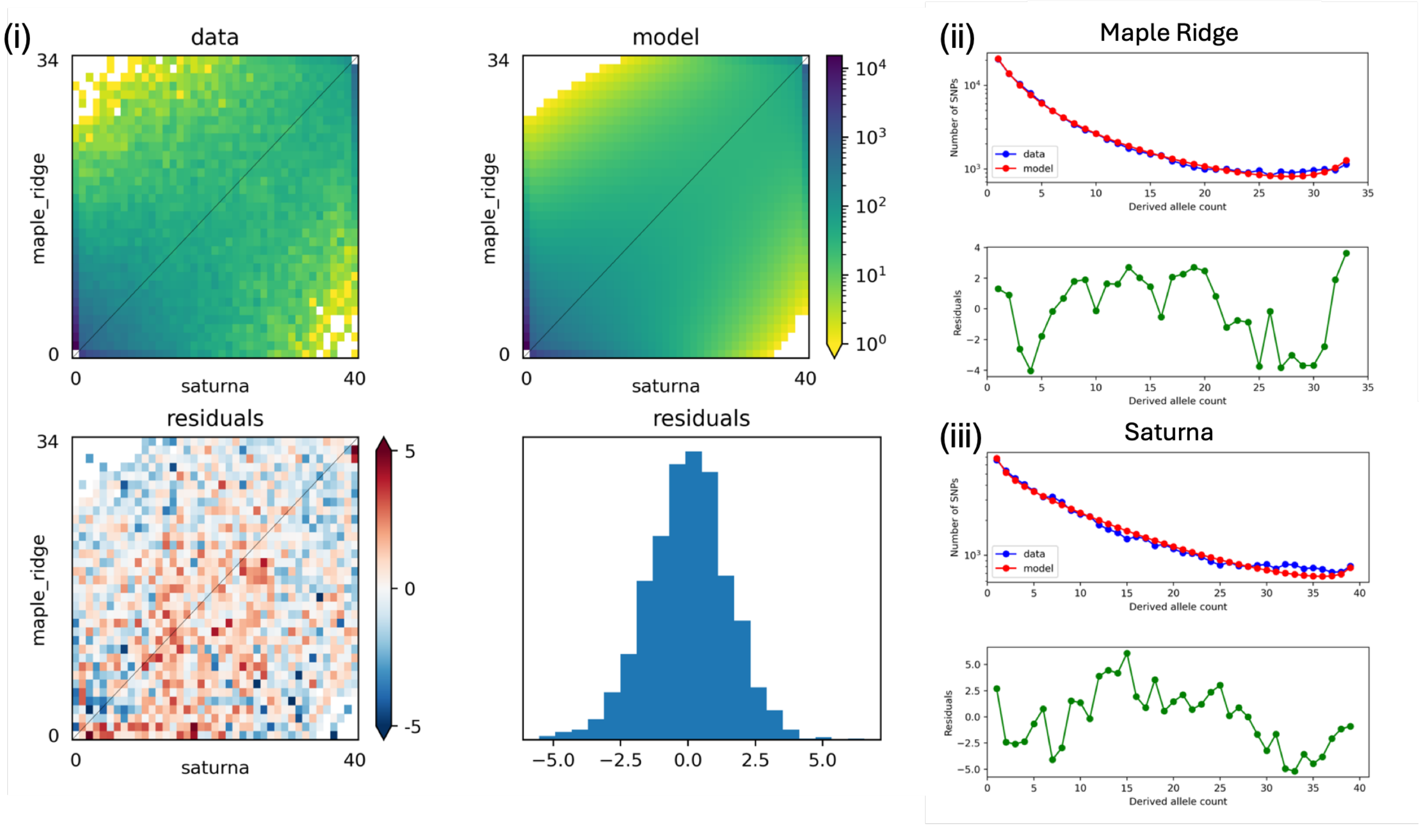


(D) Saturna-Maple Ridge residual fit; 5’ UTR mutations

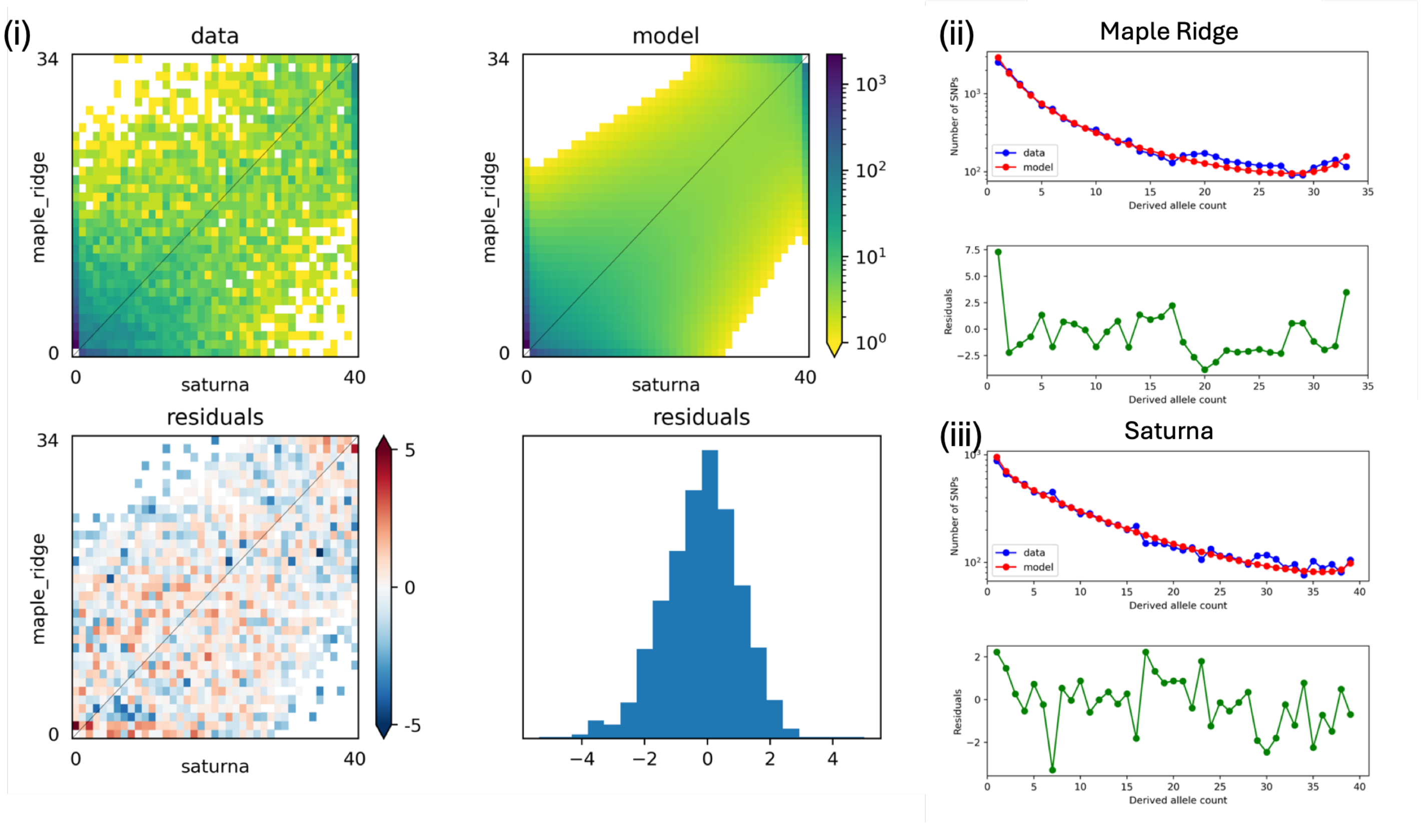

(E) Saturna-Maple Ridge residual fit; intron mutations

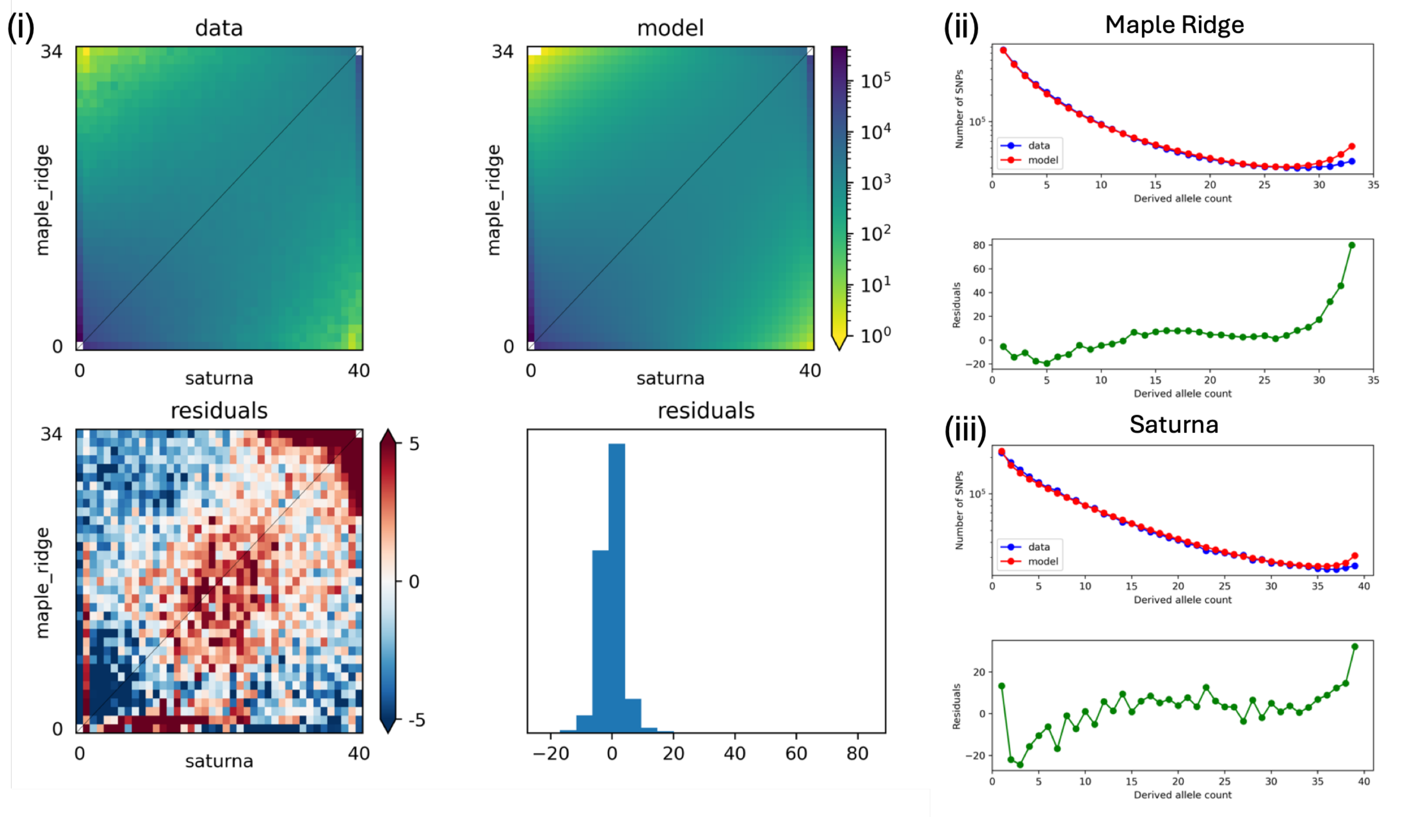


**Figure S8. Candidate joint DFE model fits to genic mutations segregating among Saturna Island and mainland Maple Ridge *P. maniculatus*.** Plots are arranged as in Supplementary Figure S5 for the Saturna-Maple Ridge comparison. See Supplementary Figure S5 legend for details.

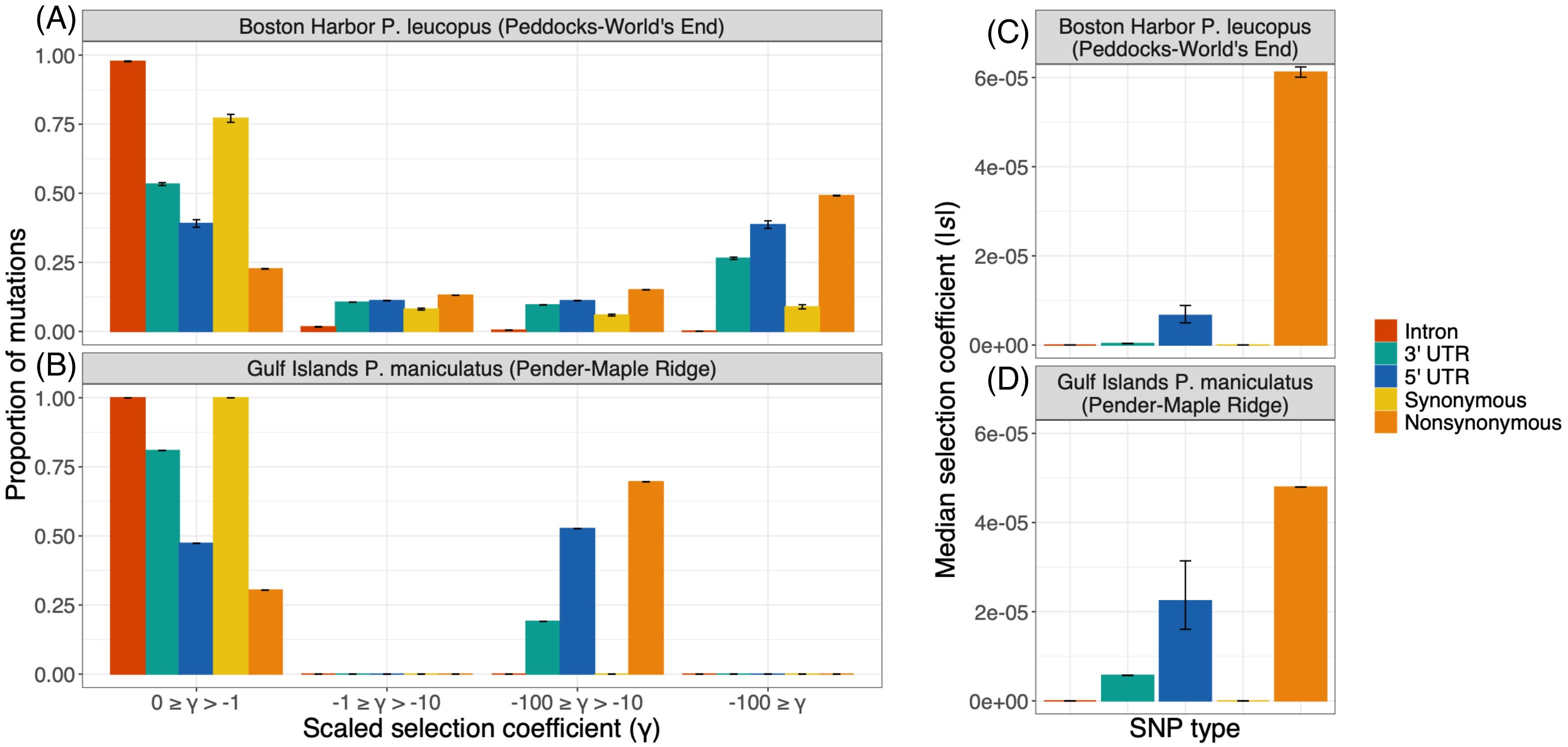


**Figure S9. Marginal DFEs capture differences in selective constraint across genic mutation types.** Panels **A** and **B** depict discretized versions of the shared marginal DFE inferred from island-mainland comparisons of Boston Harbor *P. leucopus* (Peddocks Island versus mainland World's End; **A**) and Gulf Islands *P. maniculatus* (Pender Island versus mainland Maple Ridge; **B**). Panels **C** and **D** summarize the central tendency of the distributions in **A** and **B** by the (absolute) median selection coefficient, *s*. Plots are arranged as in Figure 4. See main text for details.

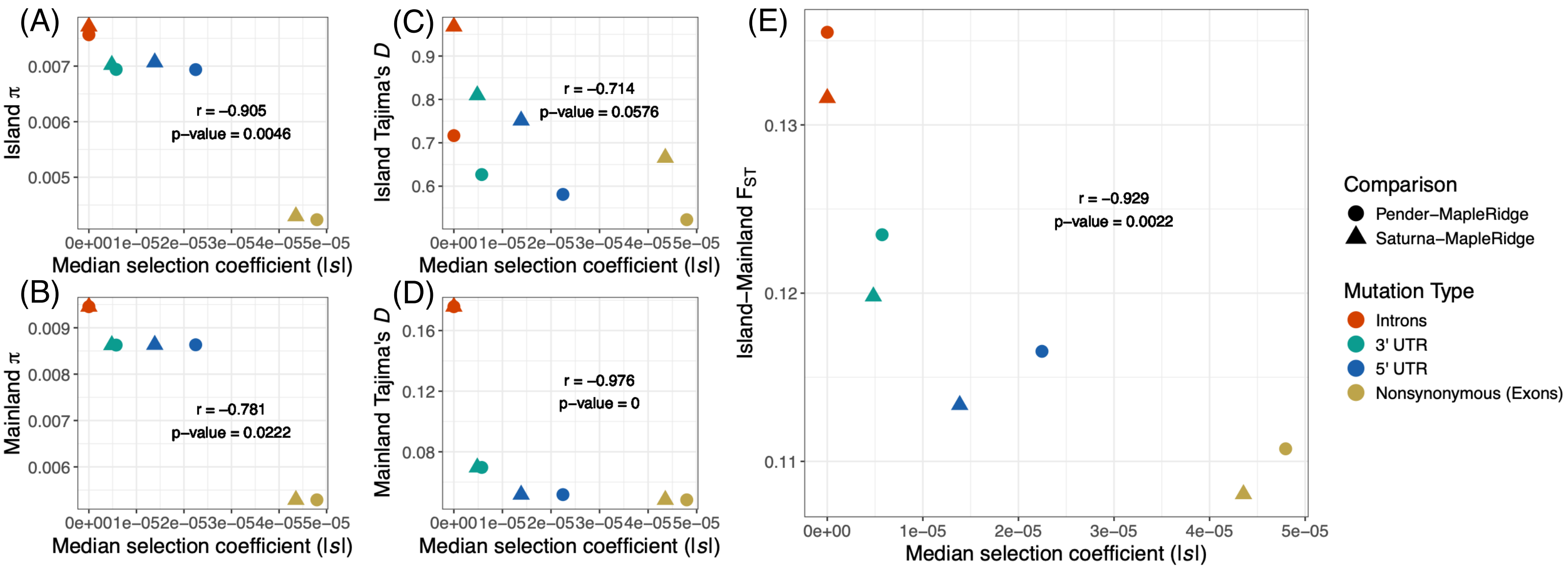


**Figure S10. Relationship between inferred |*s*| and observed summaries of variation in Gulf Islands *P. maniculatus*.** Panels compare mean nucleotide diversity (π) **(A-B)**, Tajima's D **(C-D)**, and island-mainland $F_{ST}$ **(E)** measured across genic element types (y-axes) to the median absolute selection coefficient (|*s*|) of the marginal DFE inferred for the corresponding genic mutation type (x-axes). Each point represents a specific island-mainland comparison (distinguished by point shape) and a specific element/mutation type (distinguished by point color). For exons, we compare observed summaries of variation to DFE estimates obtained for nonsynonymous mutations. Labels within each panel denote the correlation coefficient and corresponding p-value obtained using Spearman's rank correlation.

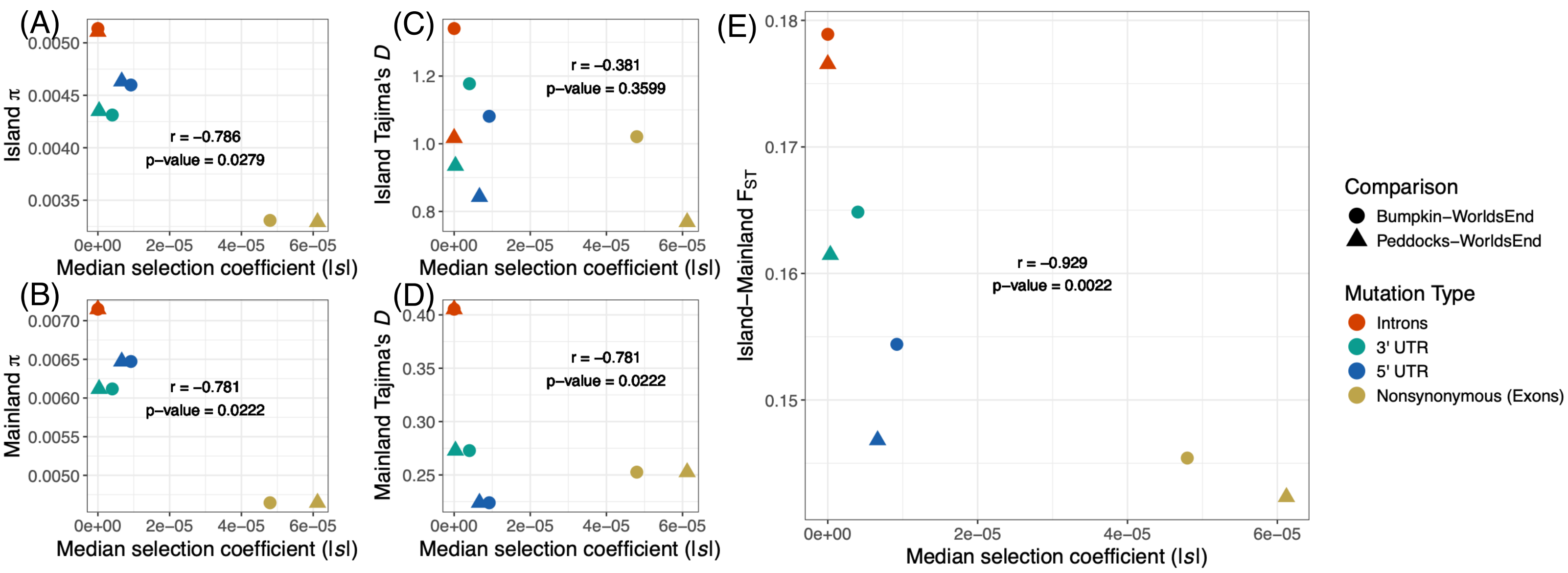


**Figure S11. Relationship between inferred |*s*| and observed summaries of variation in Boston Harbor *P. leucopus*.** Panels are arranged as in Supplementary Figure S10. See Supplementary Figure S10 legend for details.

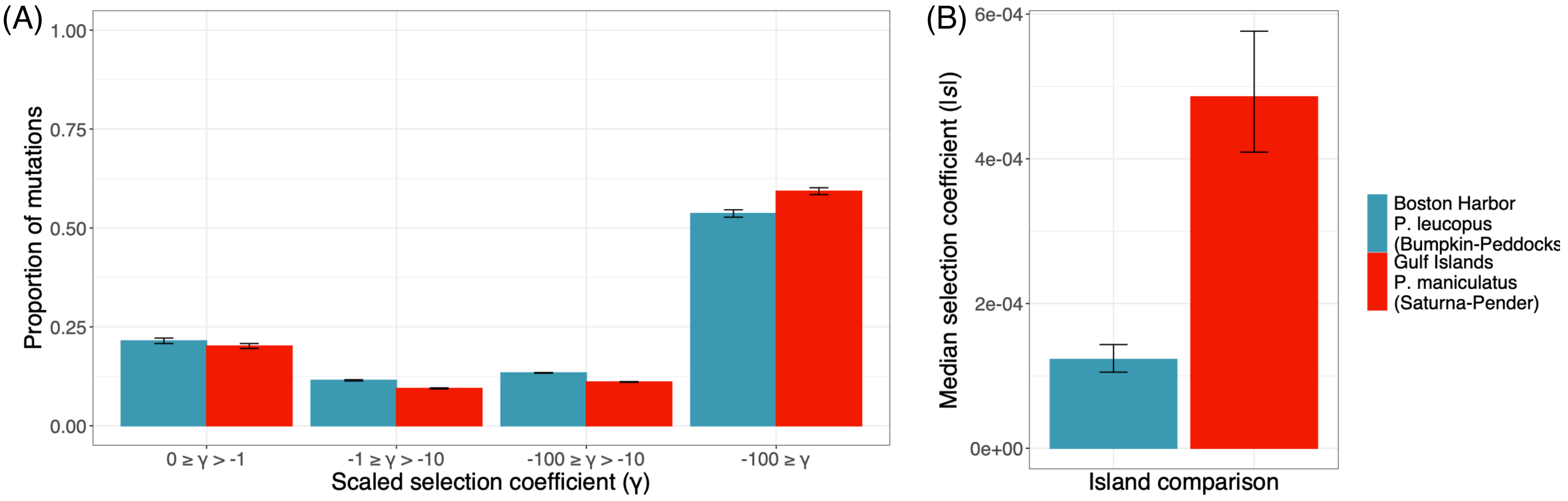


**Figure S12. Marginal DFEs inferred for nonsynonymous mutations segregating among island populations**. Panel **A** depicts discretized versions of the shared marginal DFEs inferred from nonsynonymous mutations in island-island comparisons of Boston Harbor *P. leucopus* (Bumpkin Island versus Peddocks Island; blue) and Gulf Islands *P. maniculatus* (Saturna Island versus Pender Island; red). X-axis denotes binned population scaled selection coefficients ($\gamma=2N_{Anc}s$) ranging from neutral/weakly deleterious (left-most bins) to more strongly deleterious (right-most bins). Bar height indicates the proportion of mutations falling into a given bin of γ based on the inferred DFE parameters. For each island-island comparison, error bars reflect uncertainty in the estimated μ of the lognormal distribution (see Materials and Methods for details). Panel **B** summarizes the central tendencies of the distributions in **A** by the (absolute) median selection coefficient, *s*.

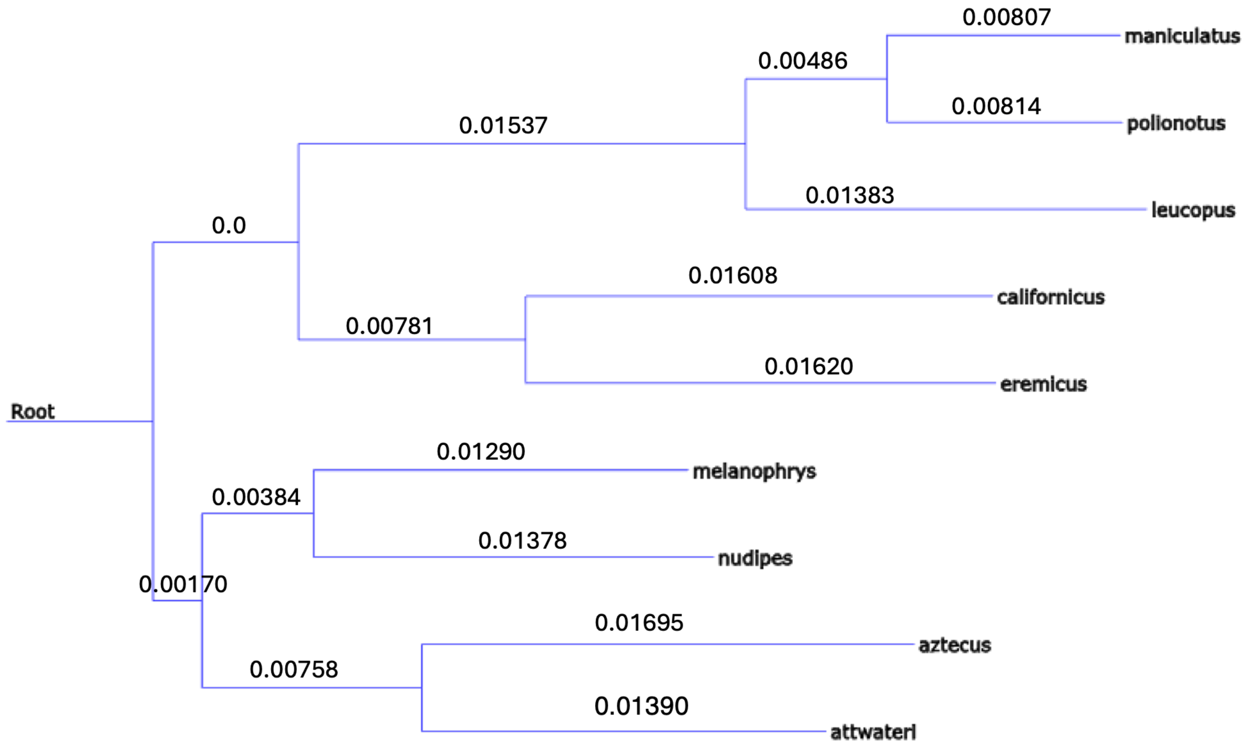


**Figure S13. Distance-based guide tree constructed from *Peromyscus* reference assemblies.** Graphical representation of the Newick-formatted output from Mashtree (Katz et al. 2019) run on the FASTA files of nine NCBI RefSeq assemblies from diverse *Peromyscus* species, visualized with T-REX (Boc et al. 2012). Branch lengths are drawn proportional to the distance-based metric computed by Mashtree (indicated by branch labels). Labels of leaves reflect species names.

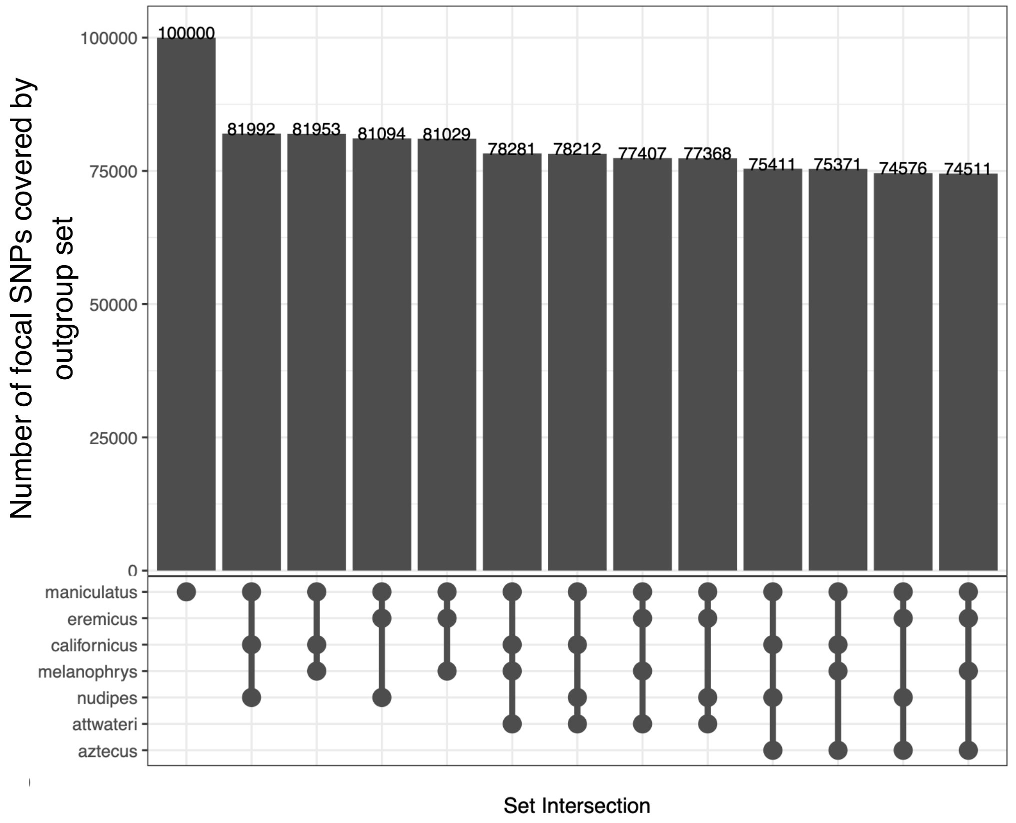


**Figure S14. Coverage of focal *P. maniculatus* SNPs by outgroup species.** Upset plot illustrates the number of polymorphic sites in our multi-population Gulf Islands *P. maniculatus* callset for which outgroup nucleotide states could be identified. This plot illustrates such "outgroup coverage" for a random sample of 100,000 genome-wide SNPs. Bottom panel depicts outgroup set. Bar height reflects the size of the intersection of polymorphic sites that are recoverable in each species in the given set (out of 100,000 SNPs).

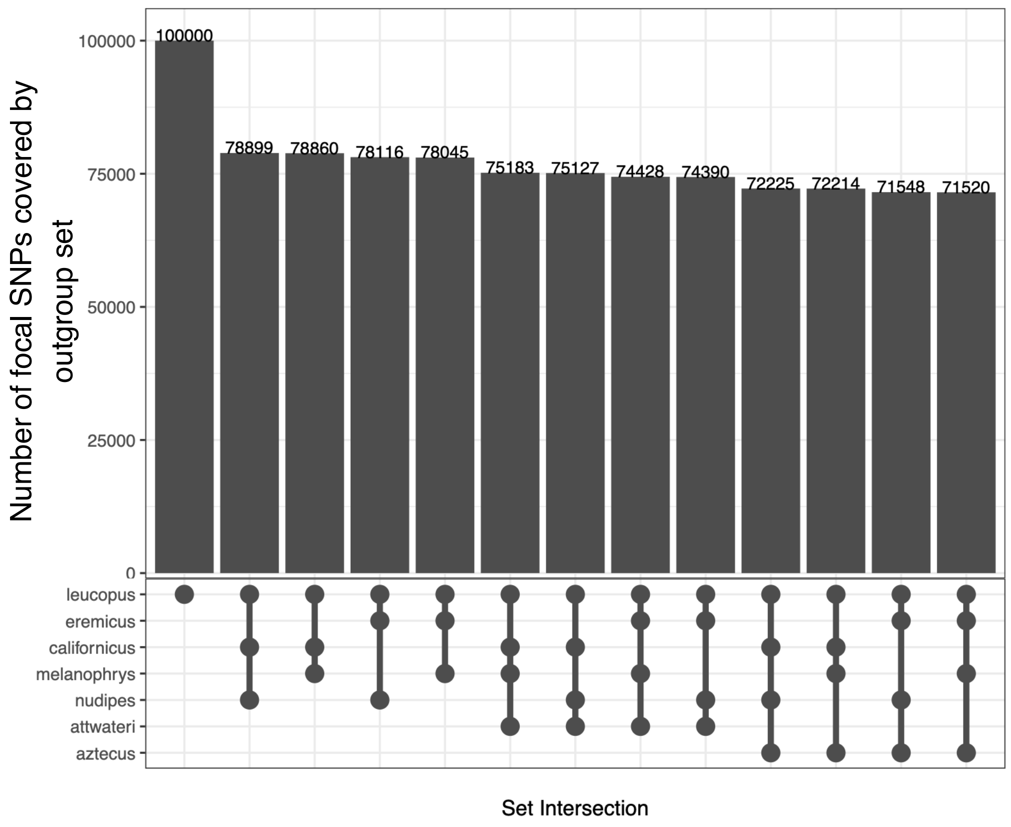


**Figure S15. Coverage of focal *P. leucopus* SNPs by outgroup species.** Upset plot is arranged as in Supplementary Figure S14 using our multi-population Boston Harbor *P. leucopus* callset as the set of focal SNPs. See Supplementary Figure S14 legend for details.

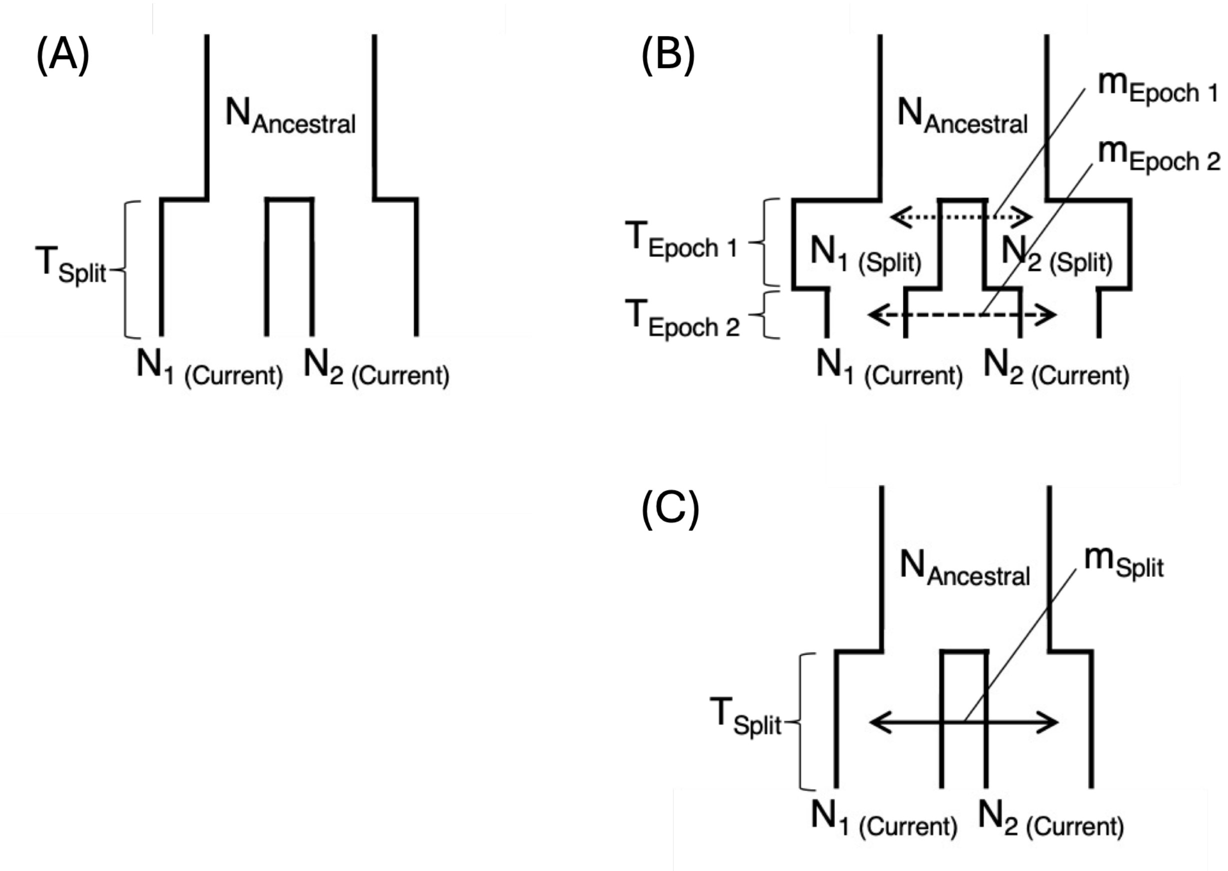


**Figure S16. Structure of 2D demographic models re-fit to unfolded site frequency spectra.** Panels depict the structure and corresponding parameters of the best-fit 2D demographic models identified in Howell et al. (2025a) for Boston Harbor *P. leucopus* **(A)** and in Howell et al. (2025b) for Gulf Islands *P. maniculatus* **(B-C)**. For Boston Harbor *P. leucopus*, the same demographic model was used for all population pairs **(A)**. For Gulf Islands *P. maniculatus*, model **B** was used for island-mainland comparisons while model **C** was used for island-island comparisons. Demographic parameters, including population sizes, migration rates, and time intervals are labeled. All migration rates are assumed to be symmetric between populations. Diagrams are not drawn to scale, and the relative magnitude of population sizes and time intervals are not constrained in one direction (e.g., populations may increase or decrease in size). In addition to depicted parameters, all models include an additional $p_{misid}$ parameter, which specifies the rate of ancestral state misidentification among polarized allele frequencies.

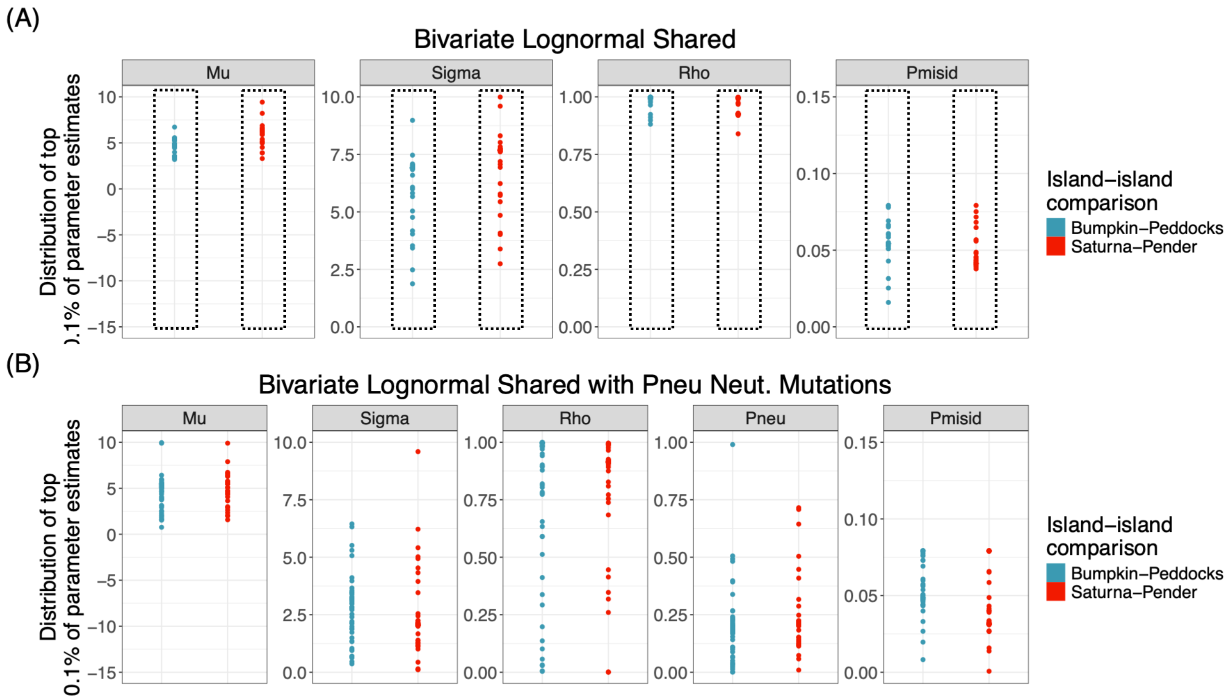


**Figure S17. Parameter convergence across tested joint DFE models for island-island comparisons.** Panels depict parameter convergence for island-island joint DFE models fit to nonsynonymous mutations. Due to the poor convergence of bivariate lognormal DFEs with distinct marginal parameters ($\mu_1 \neq \mu_2$ and $\sigma_1 \neq \sigma_2$) observed in island-mainland comparisons (Supplementary Figures S3-S4), we restricted tested island-island models to bivariate lognormal DFEs with shared marginal parameters ($\mu_1 = \mu_2$ and $\sigma_1 = \sigma_2$). Panel **A** represents a "deleterious-only" DFE, where selection coefficients are continuously distributed according to a lognormal distribution with all $\gamma < 0$. Panel **B** represents a "deleterious + neutral" DFE, where a proportion $P_{neu}$ of mutations have a fixed $\gamma = 0$ and a proportion $1 - P_{neu}$ of mutations have a continuous, lognormal distribution of deleterious selection coefficients ($\gamma < 0$). Facets within each panel reflect model parameters. Y-axes span the upper and lower bounds enforced for each parameter value. X-axis groupings distinguish the two island-island comparisons for which nonsynonymous DFEs were inferred: Bumpkin Island-Peddocks Island *P. leucopus* (blue) and Saturna Island-Pender Island *P. maniculatus* (red). Points indicate the distribution of parameter estimates observed across the top 0.1% highest likelihood parameter combinations obtained from ∂a∂i's BFGS optimization routine. Dotted boxes in panel **A** illustrate the degree of convergence of each parameter in the best-fit models presented in the main text.

(A) Bumpkin-Peddocks residual fit; nonsynonymous mutations

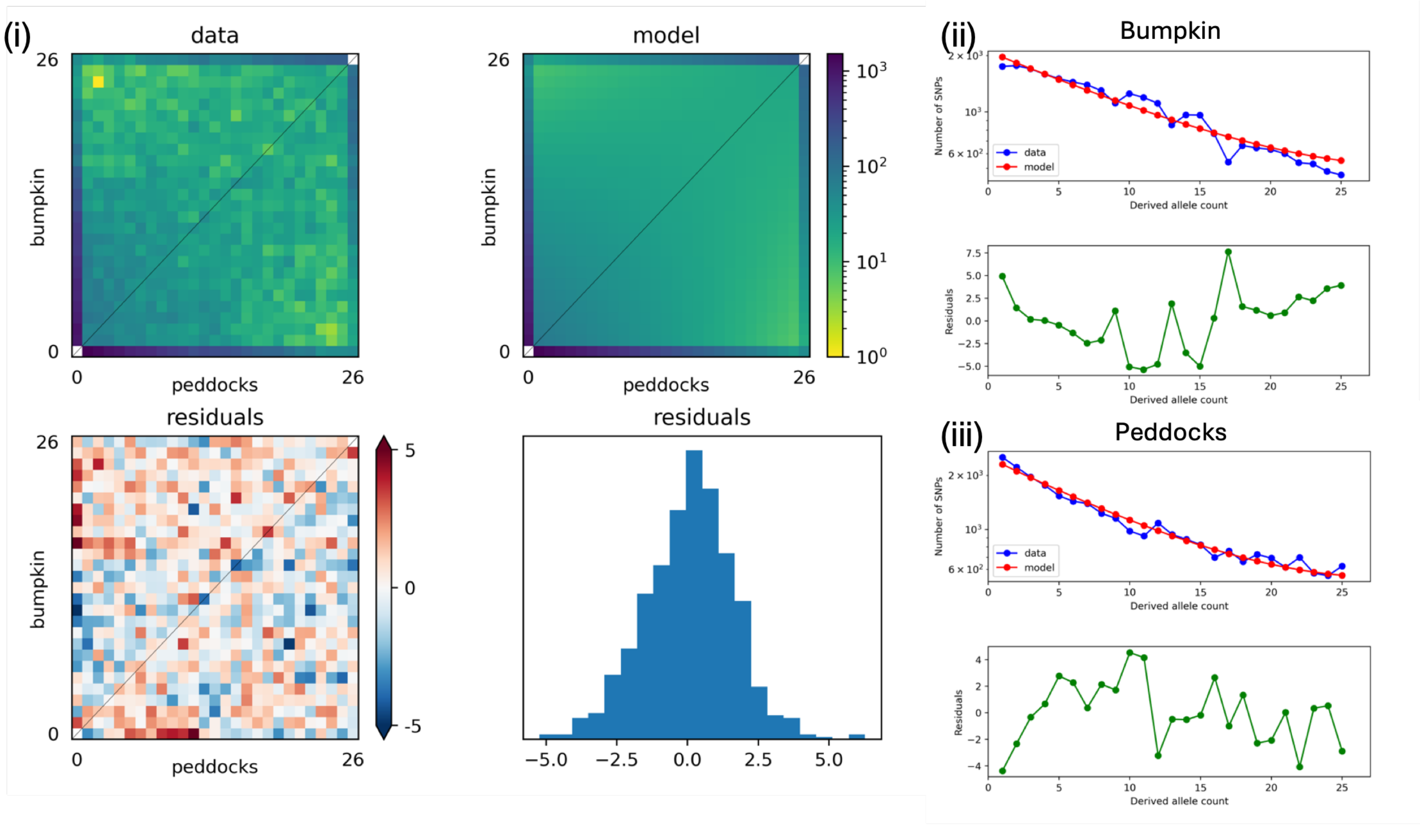


(B) Saturna-Pender residual fit; nonsynonymous mutations

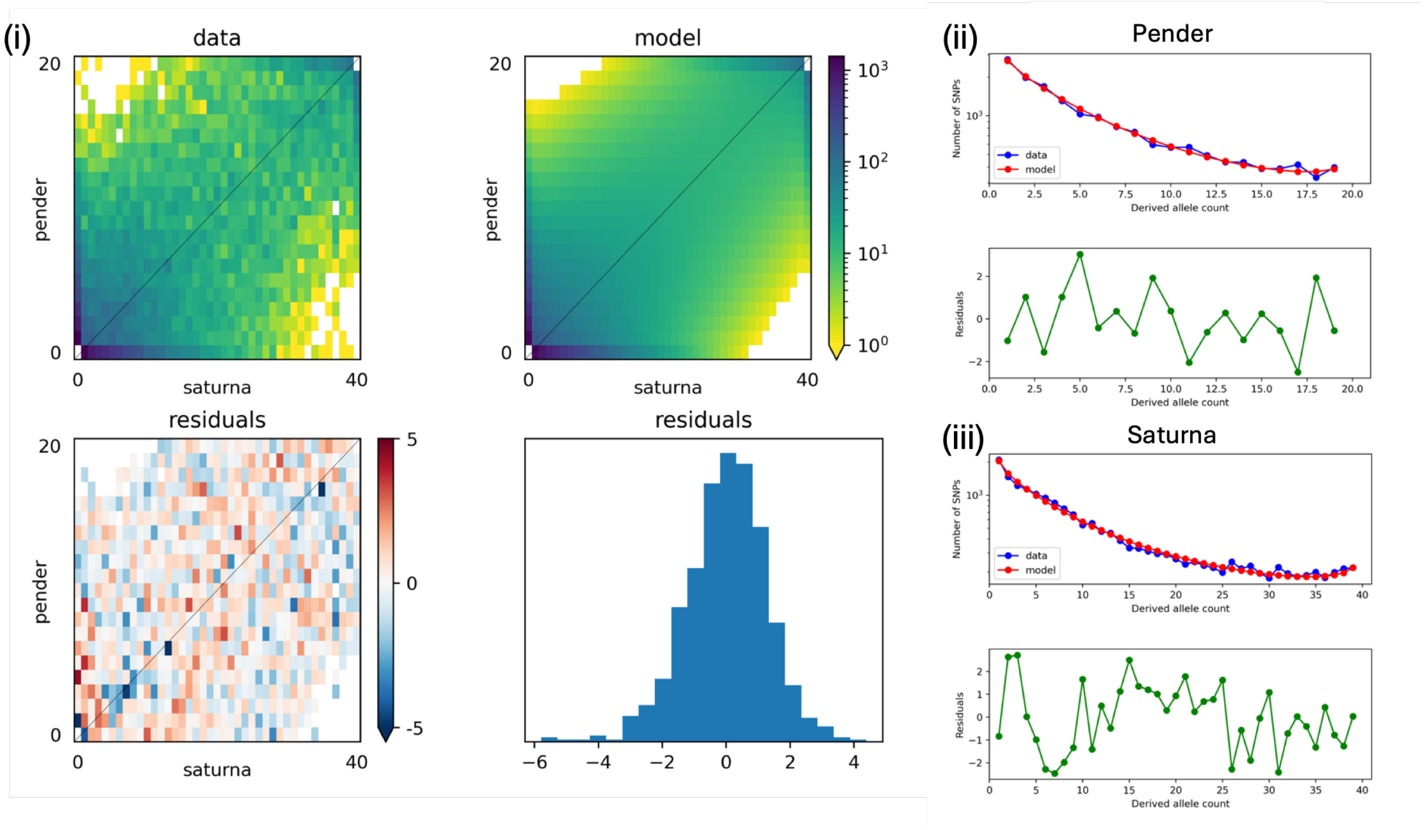

**Figure S18. Candidate joint DFE model fits for nonsynonymous mutations segregating among island populations.** Each panel depicts the fit of the inferred joint DFE model to observed allele frequencies for nonsynonymous mutations segregating among Bumpkin Island and Peddocks Island *P. leucopus* **(A)** and Saturna Island and Pender Island *P. maniculatus* **(B)**. Within each panel, facets **(i-iii)** are arranged as in Supplementary Figure S5. See Supplementary Figure S5 legend for additional details.
